# Multimodal 3D atlas of the macaque monkey motor and premotor cortex

**DOI:** 10.1101/2020.10.05.326579

**Authors:** Lucija Rapan, Sean Froudist-Walsh, Meiqi Niu, Ting Xu, Thomas Funck, Karl Zilles, Nicola Palomero-Gallagher

## Abstract

In the present study we reevaluated the parcellation scheme of the macaque frontal agranular cortex by implementing quantitative cytoarchitectonic and multireceptor analyses, with the purpose to integrate and reconcile the discrepancies between previously published maps of this region.

We applied an observer-independent and statistically testable approach to determine the position of cytoarchitectonic borders. Analysis of the regional and laminar distribution patterns of 13 different transmitter receptors confirmed the position of cytoarchitectonically identified borders. Receptor densities were extracted from each area and visualized as its “receptor fingerprint”. Hierarchical and principal components analyses were conducted to detect clusters of areas according to the degree of (dis)similarity of their fingerprints. Finally, functional connectivity pattern of each identified area was analyzed with areas of prefrontal, cingulate, somatosensory and lateral parietal cortex and the results were depicted as “connectivity fingerprints” and seed-to-vertex connectivity maps.

We identified 16 cyto- and receptor architectonically distinct areas, including novel subdivisions of the primary motor area 4 (i.e. 4a, 4p, 4m) and of premotor areas F4 (i.e. F4s, F4d, F4v), F5 (i.e. F5s, F5d, F5v) and F7 (i.e. F7d, F7i, F7s). Multivariate analyses of receptor fingerprints revealed three clusters, which first segregated the subdivisions of area 4 with F4d and F4s from the remaining premotor areas, then separated ventrolateral from dorsolateral and medial premotor areas. The functional connectivity analysis revealed that medial and dorsolateral premotor and motor areas show stronger functional connectivity with areas involved in visual processing, whereas 4p and ventrolateral premotor areas presented a stronger functional connectivity with areas involved in somatomotor responses.

For the first time, we provide a 3D atlas integrating cyto- and multi-receptor architectonic features of the macaque motor and premotor cortex. This atlas constitutes a valuable resource for the analysis of functional experiments carried out with non-human primates, for modeling approaches with realistic synaptic dynamics, as well as to provide insights into how brain functions have developed by changes in the underlying microstructure and encoding strategies during evolution.

**Highlights:**

- Multimodal analysis of macaque motor and premotor cortex reveals novel parcellation
- 3D atlas with cyto- and multireceptor architectonic features of 16 (pre)motor areas
- Primary motor area 4 is cyto- and receptor architectonically heterogeneous
- (Pre)motor areas differ in their functional connectivity fingerprints

## 1. INTRODUCTION

The primate frontal lobe encompasses two main architectonically and functionally distinct regions: a caudal part, the agranular frontal cortex, composed of motor and premotor areas, and a rostral portion which contains higher associative areas of the prefrontal cortex. The motor and premotor areas of the macaque monkey brain have been subject of multiple cytoarchitectonic, connectivity and functional studies. The ensuing maps not only differ in the nomenclature used, but also reveal considerable differences in the number of areas identified. The least detailed subdivision is that proposed by Brodmann (Brodmann, 1905), where the most caudal area represents the primary motor cortex (Brodmann’s area [BA]4, or area F1 of Matelli et al. (1985)), and the rostrally adjacent cortex is occupied by a single premotor area, BA6 (Fig. 1). Although the primary motor cortex is also described as a homogenous area in most subsequent maps (Fig. 1; Matelli et al., 1985, 1991; Preuss and Goldman-Rakic, 1991; Petrides and Pandya, 2006.; Morecraft et al., 2012; Caminiti et al., 2017), architectonic differences between the portion of BA4 located on the precentral convexity and cortex buried within the central sulcus have also been reported (Rathelot and Strick, 2009). Latter studies agree on the existence of medial, dorsal and ventral subdivisions within BA6, but they differ considerably in the number and location of such subdivisions (Fig. 1). Thus, some maps present a single premotor area on the mesial surface (Preuss and Goldman-Rakic, 1991) whereas others define two areas (Matelli et al., 1985, 1991; Petrides and Pandya, 2006.; Morecraft et al., 2012; Caminiti et al., 2017). Furthermore, whereas some authors subdivide the lateral premotor cortex into dorsal, intermediate and ventral components (Preuss and Goldman-Rakic, 1991; Morecraft et al., 2012), others postulate its subdivision into dorsal and ventral parts (Matelli et al., 1985, 1991; Petrides and Pandya, 2006.; Caminiti et al., 2017). Finally, existing maps also differ in the number of areas defined along the rostro-caudal axis of the lateral aspect of the premotor cortex.

**Figure 1.**
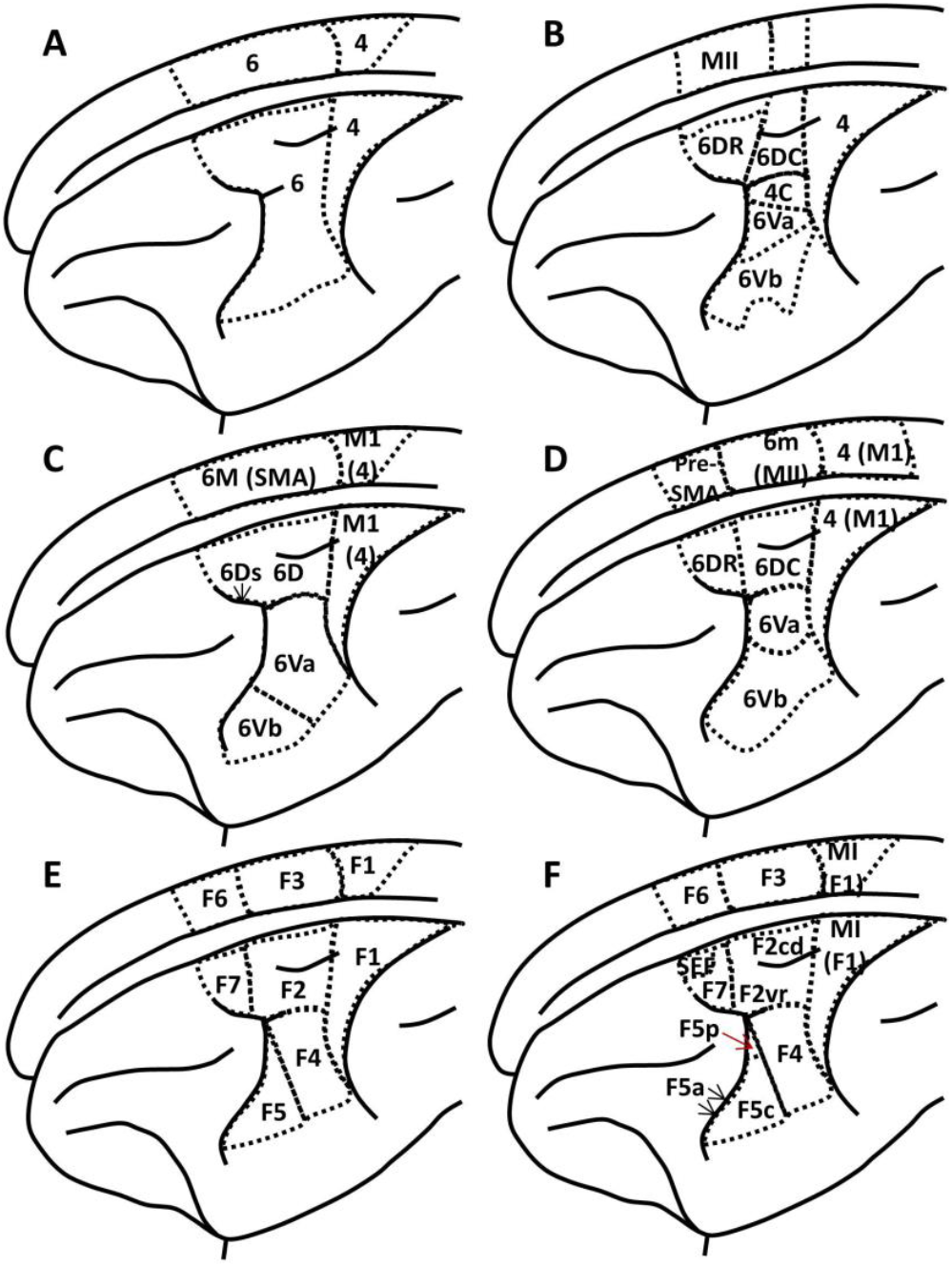
Schematic drawings of the lateral and medial views of a macaque monkey hemisphere depicting the parcellation schemes of the agranular frontal region proposed by (A) Brodmann, 1905; (B) Barbas and Pandya, 1987; (C) Preuss and Goldman-Rakic,1991; (D) Morecraft et al., 2012; (E) Matelli et al.,1985, 1991; and (F) Caminiti et al., 2017. Note, that in the map of Caminiti et al. (2017) cortical areas were defined on the basis of both architectonic and connectional criteria. Red arrow marks a small portion of area F5p on the surface, whereas black arrows indicate area F5s buried within the inferior arcuate sulcus.

The problematic of controversial results concerning the number, location and extent of cortical areas can often be explained by the fact that single different architectonic features were analyzed (e.g., cytoarchitecture or myeloarchitecture), as well as by the lack of objective and reproducible criteria for identification of cortical borders (for reviews see Zilles and Amunts, 2010; Palomero-Gallagher and Zilles, 2018). A crucial step towards overcoming these drawbacks was the development of a method which enabled the quantification of changes in the laminar distribution pattern of cell bodies and the statistical validation of such cortical borders (Schleicher and Zilles 1990; Schleicher et al., 2005; Palomero-Gallagher and Zilles, 2018; Zilles et al., 2002b). The simultaneous analysis of the regional and laminar distribution patterns of multiple transmitter receptor types as visualized by means of receptor autoradiography provides a further quantitative and statistically testable method for identification of cortical borders (Schleicher et al., 2005; Palomero-Gallagher and Zilles, 2018). Indeed, such a multimodal approach combining analysis of cortical cyto- and receptor architecture has been successfully applied in mapping studies of the human (e.g., Caspers et al., 2015; Caspers et al., 2012; Palomero-Gallagher et al., 2013) and macaque monkey (Impieri et al., 2019) brains.

In-vivo neuroimaging in the non-human primate is a promising approach to link between precise electrophysiological and neuroanatomical studies of the cortex and the large-scale networks observed in human neuroimaging. Recently non-human primate imaging has been advancing rapidly, thanks in part to increased collaborating and data-sharing (Milham et al., 2018, 2020). However, integration of neuroimaging data with high-quality postmortem anatomical data has been limited by the two disciplines not reporting results in a common coordinate space. Furthermore, parcellations of macaque cortex that are currently available to the in-vivo neuroimaging researchers do not have information relating to receptor densities. Such information is crucial to understanding the chemical underpinnings of functional activity and connectivity observed in-vivo.

The principal aim of this study is to reassess the organization of macaque motor and premotor cortex. We provide a new parcellation of these regions based on quantitative analysis of their cyto- and receptor architecture. Finally, we determine the characteristic connectivity fingerprint of each area. All data is made available to the community in standard Yerkes19 surface (Donahue et al., 2016) via the Human Brain Project and BALSA platforms.

## 2. MATERIAL AND METHODS

### 2.1 Material

The brain of an adult macaque monkey (*Macaca mulatta*; brain ID: DP1), obtained as a gift from Professor Deepak N. Pandya, was used for cytoarchitectonic analysis. After being deeply anesthetized with sodium pentobarbital, the monkey was transcardially perfused with cold saline followed by 10% buffered formalin. The brain was removed and stored in a buffered formalin solution. The brain was dehydrated in ascending graded alcohols (70% to 100% propanol) followed by chloroform, then embedded in paraffin, serially sectioned (section thickness 20 µm) in the coronal plane with a large-scale microtome, and every fifth section mounted on a gelatin coated slide. Paraffin was removed by a 10-minute incubation in Xem-200 (Vogel, Diatec Labortechnik GmbH), and sections rehydrated in descending graded alcohols (10 minutes each in 100%, 96% and 70% propanol) followed by a final rinse in pure water. Sections were stained for cell-body visualization with a modified silver method (for details, see Merker, 1983; Palomero-Gallagher et al., 2008) that provides a high contrast between cell bodies and background.

For a combined cyto- and receptor architectonic analysis, we used the brains of three adult male macaques (*Macaca fascicularis*; 6±1 years of age) which were obtained from Covance Laboratories (Münster, Germany). Monkeys were sacrificed by a lethal intravenous injection of sodium pentobarbital and the brain was immediately extracted together with meninges and blood vessels, since removing them could damage cortical layer I. Brains were then divided into left and right hemispheres, and cerebellum with brainstem. Each hemisphere was further separated into an anterior and a posterior slab at the height of the most caudal part of the central sulcus. The slabs were shock frozen in N-methylbutane (isopentane) at −40°C for 10 – 15 minutes, after which they were stored in air-tight plastic bags at −80°C until further processing. Slabs were serially sectioned (thickness 20 µm) in the coronal plane in a cryomicrotome at −20°C, thaw-mounted on gelatin-coated glass slides, air dried and stored overnight in air-tight plastic bags at −20°C. Alternating sections were processed for the visualization of cell-bodies (for details, see Merker, 1983; Palomero-Gallagher et al., 2008) or of receptor binding sites (see below).

Macaque fMRI data. A publicly available macaque fMRI dataset from a data sharing consortium PRIMate Data Exchange (PRIME-DE) was used in the present study (Milham et al., 2018, 2020). We opted for one cohort from the Oxford dataset which contains 20 macaque monkeys and 53.33 minutes fMRI scans per animal (TR=2000ms, TE=19ms, resolution=2×2×2 mm, 1600 volumes; Noonan et al., 2014). All the macaques were scanned under anesthesia. During the experiment, atropine (0.05 mg/kg, intramuscular), meloxicam (0.2 mg/kg, intravenous) and ranitidine (0.05 mg/kg, intravenous) were used to maintain the anesthetic conditions. The details of the scan and anesthesia protocols were described in Noonan et al. (2014) and on the PRIME-DE website (http://fcon_1000.projects.nitrc.org/indi/PRIME/oxford.html).

Animal care was provided in accordance with the NIH Guide for Care and Use of Laboratory Animals or the guidelines of European Communities Council Directive for the care and use of animals for scientific purposes.

### 2.2 Quantitative cytoarchitectonic analysis

Cytoarchitectonic analysis was based on an initial identification of cortical areas by visual inspection of histological sections and criteria described in the literature (Brodmann 1905, 1909; Matelli et al. 1985, 1998, 1991; Petrides and Pandya, 1994; Preuss and Goldman-Rakic, 1991; Rizzolatti et al., 1998; Belmailh et al., 2008; Schlag and Schlag-Rey, 1987), followed by the statistical validation of borders between areas. Since existing maps also differ considerably in the nomenclatures used, we here applied that of Brodmann (1909) for the primary motor cortex and that of Matelli et al. (1985, 1998, 1991) for premotor areas.

Visually identified regions of interest (ROI) were scanned by means of a light microscope (Axioplan 2 imaging, ZEISS, Germany) equipped with a motor-operated stage controlled by the KS400® and Axiovision (Zeiss, Germany) image analyzing systems applying a 6.3 x 1.25 objective (Planapo®, Zeiss, Germany), and a CCD camera (Axiocam MRm, ZEISS, Germany) producing frames of 524 x 524 µm in size, 512 x 512-pixel spatial resolution, with an in-plane resolution of 1 µm per pixel, and eight-bit grey resolution. These digitalized images were used for computation of the grey level index (GLI), i.e. the volume density of neurons measured as an areal fraction of all stained cellular forms in square measuring fields of 20-30 µm, by means of the KS400-system and in-house scripts in Matlab (The MathWorks, Inc., Natick, MA). For each area examined, GLI images were generated from three following sections on the same rostro-caudal level.

Quantification of the laminar distribution of the volume fraction of cell bodies was carried out by means of GLI profiles extracted perpendicularly to the cortical surface (for details of the GLI extraction see, Zilles et al., 2002b and Palomero-Gallagher and Zilles, 2018). The shape of a profile can be parametrized as a frequency distribution of ten features which constitute the feature vector of the profile in question, and can be used to measure (dis)similarity in cytoarchitecture (Schleicher et al., 2000). Specifically, the ten features used are the mean GLI across cortical layers (meany.o), the mean cortical depth (meanx.o, which indicates the x coordinate of the center of gravity of the area beneath the profile curve), the standard deviation (std.o), skewness (skew.o) and kurtosis (kurt.o) of the frequency distribution, as well as the corresponding values obtained from the first derivative of the profile (meany.d, meanx.d, std.d, skew.d, kurt.d,), which its local slope (Schleicher et al. 2000; Palomero-Gallagher and Zilles, 2018). We used the Mahalanobis distance (MD; Mahalanobis et al., 1949) to quantify differences in the shape of GLI profiles (Schleicher and Zilles, 1990; Schleicher et al., 1999, 2000, 2005; Zilles et al., 2002b). Profiles were analyzed for cortical borders using a sliding window procedure whereby the sliding window consisted of 10-24 adjacent profiles grouped into a block of profiles, and was moved along the cortical ribbon in single profile increments. For each block size, the MD was calculated and plotted as a distance function for all block positions. This procedure was repeated with increasing block sizes from 10 to of 24 profiles per block to control for the stability of the distance function depending on the number of profiles in a block. Blocks of profiles were used instead of single profiles, since the latter were affected by local structural inhomogeneities which reduced the signal-to-noise ratio of the distance function. To confirm and accept maxima of the distance functions as statistically significant borders, we applied Hotelling’s T^2^ test in combination with a Bonferroni adjustment of the P-values for multiple comparisons, and threshold was set at (P < 0.01) (Schleicher et al., 1999, 2000, 2005; Zilles et al., 2002b). Significant maxima identified with multiple block sizes in one section were biologically evaluated by comparison with maxima at comparable locations in three following sections to exclude maxima caused by artifacts (e.g. ruptures, folds or local discontinues in microstructure due to blood vessels.

### 2.3 Receptor architectonic analysis

We followed previously published protocols (Zilles et al., 2002b; Palomero-Gallagher and Zilles, 2018; see Tab. 1) to conduct the binding process, which includes three main steps: (i) a preincubation, where sections are rehydrated and endogenous ligands that may block the binding site removed, (ii) a main incubation, that consists of two parallel experiments, one to identify total binding of each ligand type and another to visualize non-specific binding of the same ligand, and finally, (iii) a rinsing step, where the binding process is stopped and free ligand and buffer salts are removed. To determine total binding, sections were incubated in a buffer solution with the tritiated ligand, whereas and to determine non-specific binding, neighboring sections were incubated in another buffer solution containing the tritiated ligand with a receptor type-specific displacer in a 1000-fold higher concentration. Thus, we could calculate specific binding for each ligand based on the difference between total and non-specific binding. In the present study, non-specific binding less than 5% of the total binding sites, and therefore, total binding is considered equivalent of specific binding. Finally, the radioactively labelled sections were air-dried and co-exposed against β radiation-sensitive films (Hyperfilm®, Amersham) for 4-18 weeks depending on the analyzed ligand with tritium-standards of known increasing concentrations of radioactivity (Palomero-Gallagher and Zilles, 2018).

**Table 1.** Binding protocols

| Transmitter | Receptor | Ligand (nM) | Displacer (μM) | Incubation buffer | Pre-incubation | Main incubation | Final rinsing |
| --- | --- | --- | --- | --- | --- | --- | --- |
| Glutamate | AMPA | [<sup>3</sup>H]-AMPA (10) | Quisqualate (10) | 50mM Tris-acetate (pH 7.2) [+ 100 mM KSCN]* | 3x 10 min, 4° C | 45 min, 4° C | 1) 4x 4sec<br>2) Acetone/glutaraldehyde (100 ml + 2,5 ml), 2x 2sec, 4° C |
|  | NMDA | [<sup>3</sup>H]-MK-801 (3.3) | (+)-MK-801 (100) | 50mM Tris-acetate (pH 7.2) + 50 μM glutamate [+30 μM glycine + 50 μM spermidine]* | 15 min, 4° C | 60 min, 22° C | 1) 2x 5min, 4° C<br>2) distilled water, 1x 22° C |
|  | KAIN | [<sup>3</sup>H]-Kainate (9.4) | SYM 2081 (100) | 50mM Tris-acetate (pH 7.1) [+10 mM Ca<sup>2+</sup>-acetate]* | 3x 10 min, 4° C | 45 min, 4° C | 1) 3x 4sec<br>2) Acetone/glutaraldehyde (100 ml + 2,5 ml), 2x 2sec, 22° C |
| GABA | GABA<sub>A</sub> | [<sup>3</sup>H]-Muscimol (7.7) | GABA (10) | 50mM Tris-citrate (pH 7.0) | 3 x 5 min, 4° C | 40 min, 4° C | 1) 3x 3 sec, 4° C<br>2) distilled water, 1x 22° C |
|  | GABA<sub>B</sub> | [<sup>3</sup>H]-CGP 54626 (2) | CGP 55845 (100) | 50mM Tris-HCl (pH 7.2) + 2,5 mM CaCl<sub>2</sub> | 3 x 5 min, 4° C | 60 min, 4° C | 1) 3x 2 sec, 4° C<br>2) distilled water, 1x 22° C |
|  | GABA<sub>A</sub>/BZ | [<sup>3</sup>H]-Flumazenil (1) | Clonazepam (2) | 170 mM Tris-HCl (pH 7.4) | 15 min, 4° C | 60 min, 4° C | 1) 2x 1 min, 4° C<br>2) distilled water, 1x 22° C |
| Acetylcholine | M<sub>1</sub> | [<sup>3</sup>H]-Pirenzepine (1) | Pirenzepine (2) | Modified Krebs buffer (pH 7.4) | 15 min, 4° C | 60 min, 4° C | 1) 2x 1 min, 4° C<br>2) distilled water, 1 x 22° C |
|  | M<sub>2</sub> | [<sup>3</sup>H]-Oxotremorine-M (1.7) | Carbachol (10) | 20 mM HEPES-Tris (pH 7.5) + 10 mM MgCl<sub>2</sub>+ 300 nM Pirenzepine | 20 min, 22° C | 60 min, 22° C | 1) 2x 2 min, 4° C<br>2) distilled water, 1 x 22° C |
|  | M<sub>3</sub> | [<sup>3</sup>H]-4-DAMP (1) | Atropine sulfate (10) | 50 mM Tris-HCl (pH 7.4) + 0.1 mM PSMF + 1mM EDTA | 15 min, 22° C | 45 min, 22° C | 1) 2x 5 min, 4° C<br>2) distilled water, 1 x 22° C |
| Noradrenaline | α<sub>1</sub> | [<sup>3</sup>H]-Prazosin (0.2) | Phentolamine Mesylate (10) | 50mM Na/K-phosphate buffer (pH 7.4) | 15 min, 22° C | 60 min, 22° C | 1) 2x 5 min, 4° C<br>2) distilled water, 1 x 22° C |
|  | α<sub>2</sub> | [<sup>3</sup>H]-UK 14,304 (0.64) | Phentolamine Mesylate (10) | 50 mM Tris-HCl + 100 μM MnCl<sub>2</sub> (pH 7.7) | 15 min, 22° C | 90 min, 22° C | 1) 5 min, 4° C<br>2) distilled water, 1 x 22° C |
| Serotonin | 5-HT<sub>1A</sub> | [<sup>3</sup>H]-8-OH-DPAT (1) | 5-Hydroxy-tryptamine, (1) | 170 mM Tris-HCl (pH 7.4) [+4 mM CaCl<sub>2</sub>+ 0.01% ascorbate]* | 30 min, 22° C | 60 min, 22° C | 1) 5 min, 4° C<br>2) distilled water, 3 x 22° C |
|  | 5-HT<sub>2</sub> | [<sup>3</sup>H]-Ketanserin (1.14) | Mianserin (10) | 170 mM Tris-HCl (pH 7.7) | 30 min, 22° C | 120 min, 22° C | 1) 2x 10 min, 4° C<br>2) distilled water, 3 x 22° C |

\* substances in brackets only included in the main incubation, \*\* substances in brackets only included in the pre-incubation

After films were developed, autoradiographs were digitized with an image analysis system consisting of a source of homogenous light and a CCD-camera (Axiocam MRm, Zeiss, Germany) with an S-Orthoplanar 60-mm macro lens (Zeiss, Germany) corrected for geometric distortions, connected to the image acquisition and processing system Axiovision (Zeiss, Germany), in order to carry out densitometric analysis of binding site concentrations in the autoradiographs (Zilles et al., 2002b; Palomero-Gallagher and Zilles, 2018). Spatial resolution of the resulting images was 3000 x 4000 pixels; 8-bit gray value resolution. Since the gray values of the digitized autoradiographs represent concentration levels of radioactivity, a scaling (i.e. a linearization of the digitized autoradiographs) had to be performed in which the gray values were transformed into fmol binding sites/mg protein using in house developed Matlab (The MathWorks, Inc. Natrick, MA) scripts. This process required two steps: (i) the gray value images of the plastic tritium-standards were used to compute the calibration curve, which defines the non-linear relationship between gray values and concentrations of radioactivity; (ii) radioactivity concentration R was then converted to binding site concentration Cb in fmol/mg protein using equation 1:

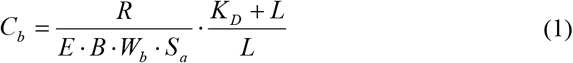

where E is the efficiency of the scintillation counter used to determine the amount of radioactivity in the incubation buffer (depends on the actual counter), B is the number of decays per unit of time and radioactivity (Ci/min), Wb the protein weight of a standard (mg), Sa the specific activity of the ligand (Ci/mmol), KD the dissociation constant of the ligand (nM), and L the free concentration of the ligand during incubation (nM). The result was a linearized image in which the gray value of each pixel in the autoradiograph is converted into a receptor density in fmol/mg protein (for details see Zilles et al., 2002b; Palomero-Gallagher and Zilles 2018). To visualize the distribution pattern of each receptor type throughout the cortex, we applied pseudo-color coding of autoradiographs by means of linear contrast enhancement, which preserves the scaling between gray values and receptor concentrations. Equally spaced density ranges were assigned to a spectrum of eleven colors, where red was assigned to highest and black to lowest receptor concentration levels. If five or more different receptor types showed transition from higher to lower concentration levels, or vice versa, at the same cortical position, we confirmed the presence of a receptor architectonic border.

Measurement of receptor densities was performed by computing the surface below receptor profiles, which were extracted from the linearized autoradiographs using in house developed scripts for Matlab (The MathWorks, Inc. Natrick, MA) in a manner analog to the procedure described above for GLI profiles. Unlike for cytoarchitectonic analysis, where the outer contour line was placed at border between layers I and II, for receptor profiles, the outer contour line followed the pial surface. We thus calculated mean densities (i.e., averaged over all cortical layers) of each of the 13 different receptors in all 16 cytoarchitectonically defined areas for each of the three left hemispheres. The ensuing densities were visualized as “receptor fingerprints”, i.e., as polar coordinate plots simultaneously depicting the concentrations of all examined receptor types within a given cortical area (Zilles et al., 2002a). In order to perform accurate sampling of each microscopically defined area, we compared autoradiographs with the adjacent sections which had been processed for the visualization of cell bodies.

### 2.4 2D and 3D maps of the macaque agranular frontal cortex

In order to display the spatial relationship between all defined areas, including those located deep in sulci, we created a 2D framework based on the macroanatomical landscape of the brain processed solely for the visualization of cell bodies (i.e. DP1). Every 40^th^ section was presented as a simple geometrical pattern by means of Adobe Illustrator CS6, thus creating a “scaffold” on which the position of cytoarchitectonic borders could be traced relative to the macroscopic landmarks (sulci and dimples). Within a section, each area was labeled with a specific color, which was further connected to the same color portion on the following sections, creating a continuous shape of each area. Thus, the 2D parcellation scheme not only enables visualization of areas and borders even when located inside sulci, but also reveals interhemispheric differences.

Location and extent of the motor and premotor areas were delineated in the 3D space of the Yerkes19 surface (Donahue et al., 2016) by LR, using the connectome workbench software (https://www.humanconnectome.org/software/connectome-workbench) by carefully aligning boundaries to macroanatomical landmarks identified using the cytoarchitecture. The location of all regions on the Yerkes19 surface were independently checked and verified by MN, SFW and NPG. 3D reconstruction of the hemisphere was obtained using the Connectome Workbench software. Additionally, the mean receptor densities of all 13 receptor types have been projected onto the corresponding area on the Yerkes19 surface for visualization. Color bars in the ensuing figures code for receptor densities in fmol/mg protein.

### 2.5 Analysis of functional connectivity

The structural and functional data were preprocessed using the Human Connectome Project-style pipeline for Nonhuman Primate and described previously (Autio et al., 2020; Xu et al., 2019). For each macaque, the structural preprocessing includes denoising, skull-stripping, tissue segmentation, surface reconstruction and surface registration to align to Yerkes19 macaque surface template. The functional preprocessing includes temporal compressing, motion, correction, global mean scaling, nuisance regression (Friston’s 24 motion parameters, white matter, cerebrospinal fluid), band-pass filtering (0.01-0.1Hz), and linear and quadratic detrending. The preprocessed data then were co-registered to the anatomy T1 and projected to the middle cortical surface. Finally, the data were smoothed (FWHM=3mm) on the high-resolution native surface, aligned and down resampled to a 10k surface (10,242 vertices per hemisphere).

The pre-processed BOLD activity timecourses for each monkey were demeaned and then concatenated in time. In order to parsimoniously describe the pattern of functional connectivity across the cortex, we calculated ‘connectivity fingerprints’ of each area. Connectivity fingerprints, originally inspired by receptor fingerprints (Passingham et al., 2002; Mars et al., 2018) aim to describe the unique pattern of connectivity of each cortical area with other areas across the cortex.

Here we chose to investigate the connectivity of each of the newly defined premotor and motor areas with 23 areas of prefrontal, cingulate, somatosensory and lateral parietal cortex, as defined by the Lyon atlas of Kennedy and colleagues (Markov et al., 2014). We calculated a representative time course for each of the 16 newly defined premotor and motor areas and the 23 prefrontal, cingulate, somatosensory and lateral parietal areas, giving 39 areas in total. For each of the 39 areas, we performed a principal components analysis on activity across all vertices within the area. The first principal component was taken as the representative activity timecourse for each area.

We used the representative timecourses of each of the 16 motor/premotor areas as seeds for functional connectivity analysis. The representative timecourses were correlated with the activity timecourses for each vertex on the surface using a Pearson correlation. A Fisher’s r-to-Z transformation was then applied to each of the correlation coefficients. This was visualized on the cortical surface. To quantify the connectivity fingerprint, we performed Pearson correlations between activity in each of the 16 premotor/motor areas and the 23 prefrontal, cingulate, somatosensory and lateral parietal areas. A Fisher’s r-to-Z transformation was also applied to each of the correlation coefficients. This was then displayed as a spider plot in order to visualize the connectivity fingerprint.

### 2.6 Statistical analyses

#### 2.6.1. Receptor densities

Statistical testing was used to determine if there were significant differences in receptor densities between adjacent regions. Testing was performed using linear mixed-effects models because these can account for repeated measures within the same subject. Prior to statistical analysis, receptor density values were normalized within each receptor type. All statistical analysis was conducted using the R programming language (version: 3.6.3) (R Core Team, 2020).

A first omnibus test of all regions and receptors was performed to establish if there were any significant differences in receptor density between all regions and receptor types. The model consisted of fixed effects for area, receptor type, the interaction between area and receptor type. The random effects in the model consisted in a random intercept for each macaque brain and receptor type (Equation 2).

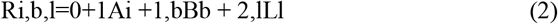

where R is the receptor density, A is motor area, B is macaque brain, and L is ligand.

A second set of tests were used to determine if pairs of adjacent regions were significantly different from one another over all receptor types. The linear mixed effect model used for the second series of tests had the same form as the omnibus test, but was only applied to pairs of adjacent regions. The p-values for the main effect “area” were corrected for multiple comparisons using the Benjemani-Hochberg correction for false-discovery rate (Benjamini and Hochberg, 1995) and significance threshold was set at p < 0.05.

Finally, for pairs of areas that were significantly different from one another in the second level tests after correction for multiple comparisons, a third linear mixed-effect model was used to test for a difference in the density between paired regions for each receptor type, respectively. The model was composed of a fixed effect for area and a random interceptor for each macaque brain (Equation 3). The p-values for the fixed-effect “area” from each of these tests were again corrected using the Benjemani-Hochberg correction for false-discovery rate (Benjamini and Hochberg, 1995) and significance threshold was set at p < 0.05.

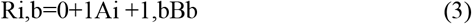

where R is the receptor density, A is motor area, B is macaque brain.

Additionally, principal components and hierarchical cluster analysis were carried out to determine degree of similarity of the receptor fingerprints. Receptor densities were normalized by z-scores prior to multivariate analyses since absolute densities vary considerably among receptors, and without a normalization step, receptors exhibiting high absolute densities would dominate the computation of Euclidean distances. For the hierarchical cluster analysis, the Euclidean distance was used as a measure of (dis)similarity since it takes both differences in the size and in the shape of receptor fingerprints into account. The Ward linkage algorithm was chosen as the linkage method, since in combination with the Euclidean distance it resulted in the maximum cophenetic correlation coefficient as compared to any combination of alternative linkage methods and measurements of (dis)similarity. The number of stabile clusters was determined by a k-means analysis and the elbow method (Rousseeuw, 1987).

#### 2.6.2 Functional connectivity

Linear mixed-effect models were also used to test if motor regions differed based on the strength of functional connectivity with other brain regions. Prior to statistical analysis, connectivity values to a specific region were normalized by dividing the standard deviation of all connectivity values to that region.

A first omnibus test of all regions and receptors was performed to establish if there were any significant differences in connectivity strength between all motor regions. The model consisted of a fixed effect for the motor areas and a random effect for the macaque brains from which the connectivity measures were acquired. Similar to the second tests for receptor density, post-hoc testing was performed by using the same model as for the omnibus test but only comparing 2 neighboring regions at a time (Equation 4). This made it possible to test if any two adjacent regions differed on the basis of the strength of functional connectivity to a set of target regions. As before, the p-values for the fixed-effect “area” from each of these tests were corrected using the Benjemani-Hochberg correction for false-discovery rate (Benjamini and Hochberg, 1995) and significance threshold was set at p < 0.05.

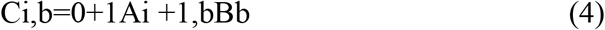

where C is the connectivity strength, A is the motor area, B is the brain from which a particular connectivity strength measure was acquired.

## 3. RESULTS

We identified 16 cyto- and receptor architectonically distinct areas within the macaque agranular frontal region, three of which were classified as primary motor areas (4a, 4p and 4m), and 13 as premotor areas (F2d, F2v, F3, F4d, F4v, F4s, F5d, F5v, F5s, F6, F7d, F7i, F7s). A 2D flat map scheme (Fig. 2) displays their distribution and location relative to macroanatomic landmarks on the medial and dorsolateral surfaces of monkey brain DP1. The main advantage of this map is that it not only shows the extent of areas found on the brain surface, but also those located within sulci. Furthermore, it presents the actual spatial relationship between cortical borders and macroanatomic features such as sulci and dimples in both hemispheres, and highlights the low degree of interhemispheric variability.

**Figure 2.**
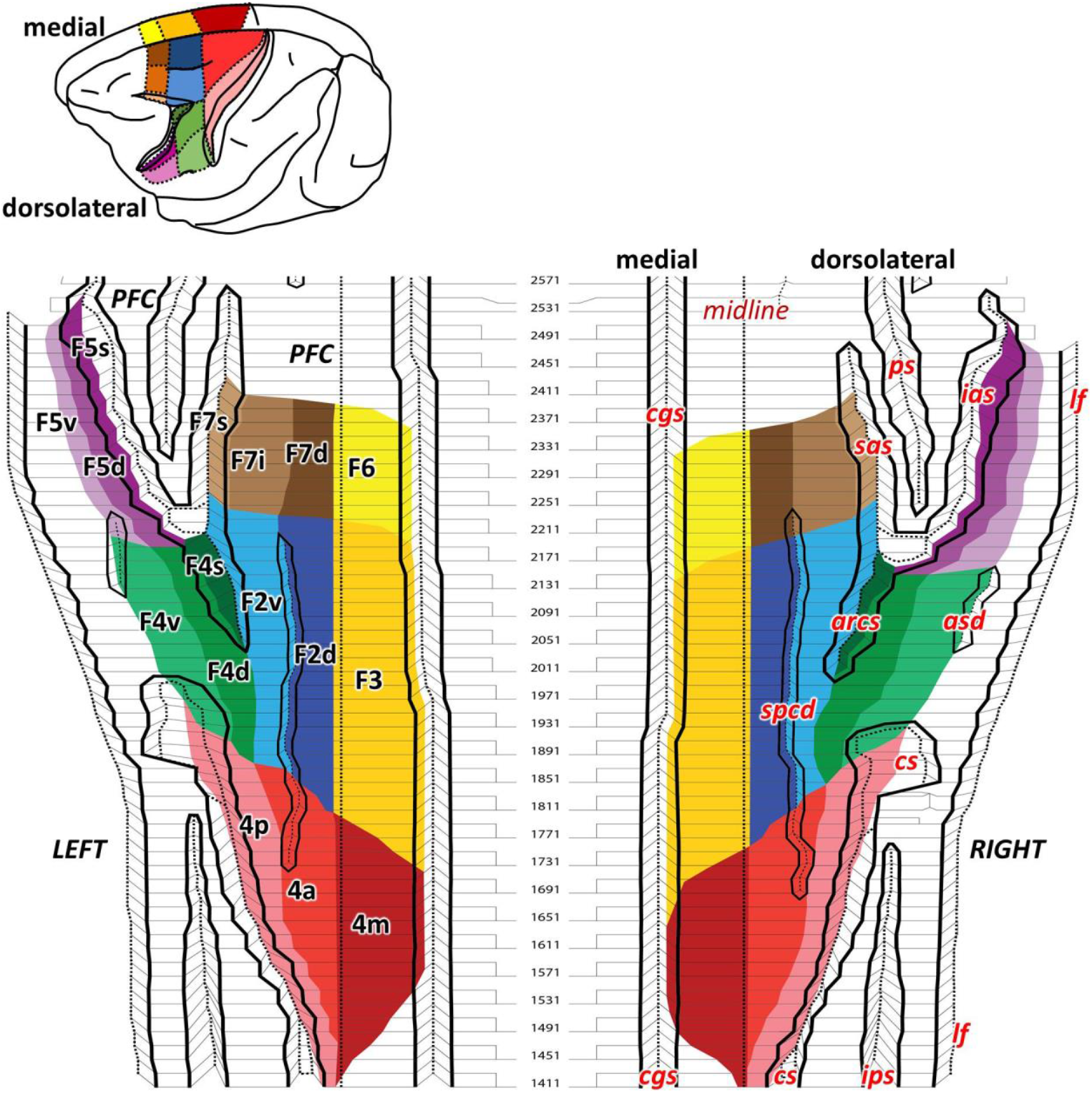
2D flat map depicting all identified areas on the medial and dorsolateral premotor surfaces (a total of 13 premotor and 3 motor areas). Areas are marked on the left hemisphere and microanatomical features on the right hemisphere. Black full lines mark the sulci and dimple borders on the surface, whereas dashed black lines represent fundus. The only dashed black line on the surface marks the midline, which segregates medial and dorsolateral cortical surface. Section number (every 40^th^) indicated between the hemispheres. arcs – spur of the arcuate sulcus, asd – anterior supracentral dimpl, cgs – cingulate sulcus, cs – central sulcus, ias – inferior arcuate branch, ips – inferior parietal sulcus, lf – lateral fissure, ps – principal sulcus, sas – superior arcuate branch, spcd – superior precentral dimple.

The central sulcus (*cs*) serves as a clear landmark to locate the most posterior border of the macaque motor cortex, where it abuts the somatosensory cortex (Fig. 2). Specifically, the border between primary motor area 4p and somatosensory area 3a was always found in the fundus of the cs. On the lateral surface, the superior (*sas*) and inferior (*ias*) arcuate sulci, together with the spur of the arcuate sulcus (*arcs*), form a letter Y that has the appearance of a physical border between the generally granular prefrontal cortex and the agranular motor region. The arcs served as a landmark to separate dorsal and ventral premotor areas. On the medial surface of the hemisphere, the border between premotor and motor areas and the ventrally adjacent cingulate cortex is found on the dorsal bank of the cingulate sulcus (*cgs*). Other useful macroanatomical features, though not as deep as sulci, and more prone to individual variability, are dimples. The most prominent one is the superior precentral dimple (*spcd*) that extends from primary motor cortex to the rostral premotor areas on the dorsal convexity in both hemispheres, although it is longer in the right hemisphere than in left one. Finally, the anterior subcentral dimple (*asd*) roughly indicates the ventral extent of the premotor areas.

### 3.1 Cytoarchitecture

#### 3.1.1. Primary motor cortex

Area 4 is characterized by the presence of unusually large pyramidal cells (known as Betz cells; Betz, 1874) in sublayer Vb (Fig. 3). Although the macaque primary motor cortex has generally been described as a homogenous area, we identified three subdivisions based on their distinct cyto- and receptor architecture: area 4m on the medial aspect of the hemisphere, area 4a on the dorsolateral surface, and area 4p extending from the edge of the dorsal surface of the hemisphere to the fundus of the *cs*, where it abuts somatosensory area 3a (Fig. 2).

**Figure 3.**
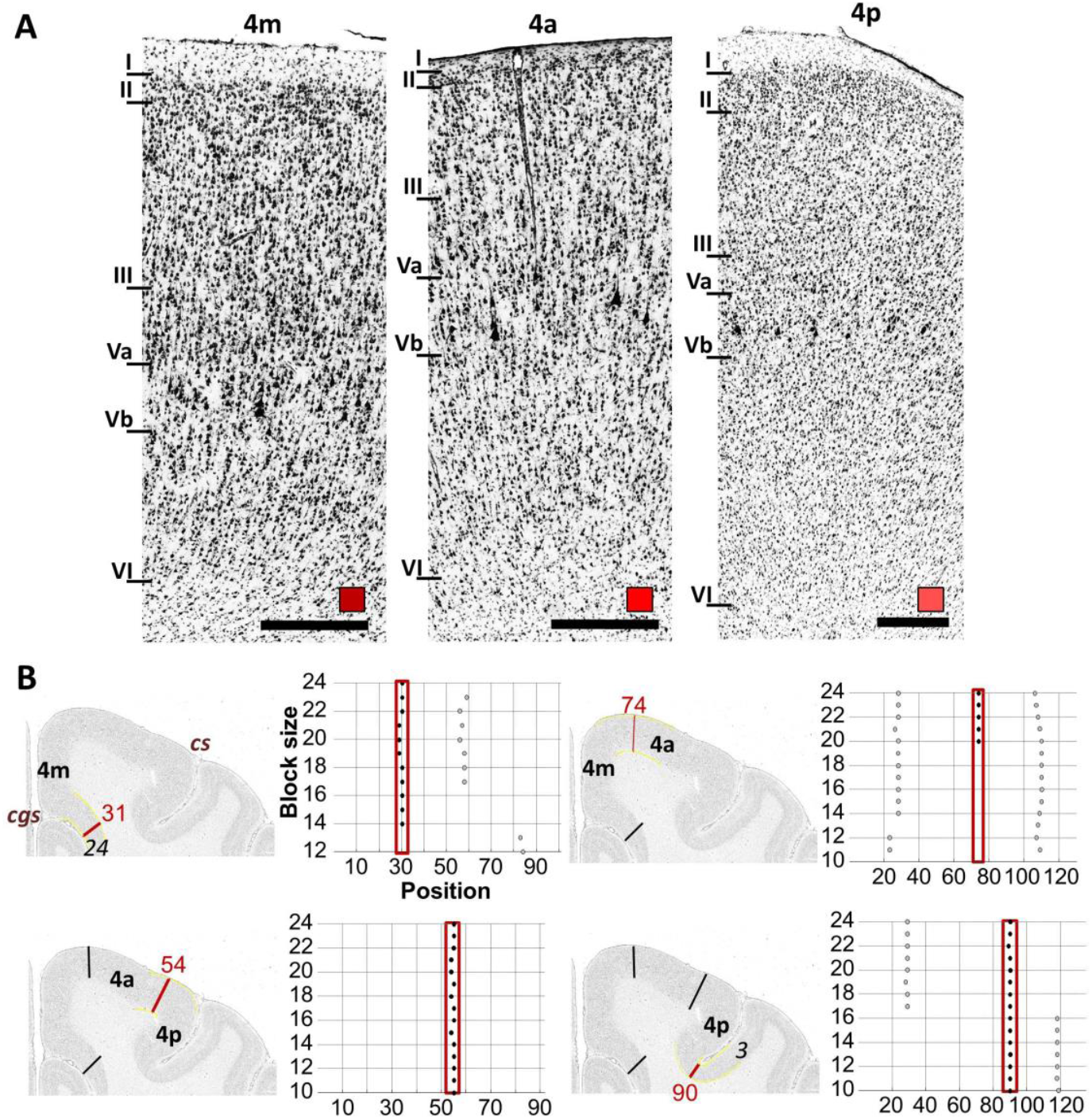
(<u>A)</u> Cytoarchitecture of the medial (4m), anterior (4a) and posterior (4p) subdivisions of the primary motor cortex, area 4. Colored square over the scale bar indicates the color used in Figure 2 to code the area in question. (<u>B)</u> Quantitative analysis of cytoarchitectonic borders. The position of each border verified by the statistical analysis of Mahalanobis distances is highlighted by a red line (and corresponding profile index) on the GLI-image, and the corresponding dot plot (depicted to the right of the GLI-image) reveals that the location of significant maxima in the distance function (indicated by each dot) does not depend on the block size, but remains constant over large block size intervals (highlighted by the red frame). Roman numerals indicate cortical layers. Scale bars 1 mm. cgs – cingulate sulcus, cs - central sulcus.

Area 4a has the strongest laminar appearance of all subdivisions, due to a lower cell-body packing density in layers III and VI. This difference is particularly apparent when compared to 4p, where only layers II and Vb (due to the presence of Betz cells) are prominent. Additionally, 4a has a significantly thinner layer I in regard to surrounding areas (Fig. 3A). Area 4m is distinguishable by a prominent vertical cell organization in layers Vb and VI. The same columnar pattern is also visible in adjoining area 4a, but only in layer V. Furthermore, the border between layers III and Va is not as clear in 4m as in 4a. Figure 3B provides an example of the statistical confirmation of visually identified cortical borders.

#### 3.1.2. Medial premotor cortex

Dorsally above the *cgs*, on the medial surface of the hemisphere we identified two premotor areas: F3 (Fig. 4) and F6 (Fig. 5). F3 is found caudal to F6 and shares a border with area 4m, while F6 is delimited rostrally by prefrontal area 8. Both F3 and F6 expand a little above the midline and encroach onto the dorsal surface of the hemisphere, where they abut areas F2d and F7d, respectively. Finally, ventrally the border between medial premotor areas and the cingulate cortex was consistently found on the dorsal bank of the of the cingulate sulcus, though very close to the fundus.

**Figure 4.**
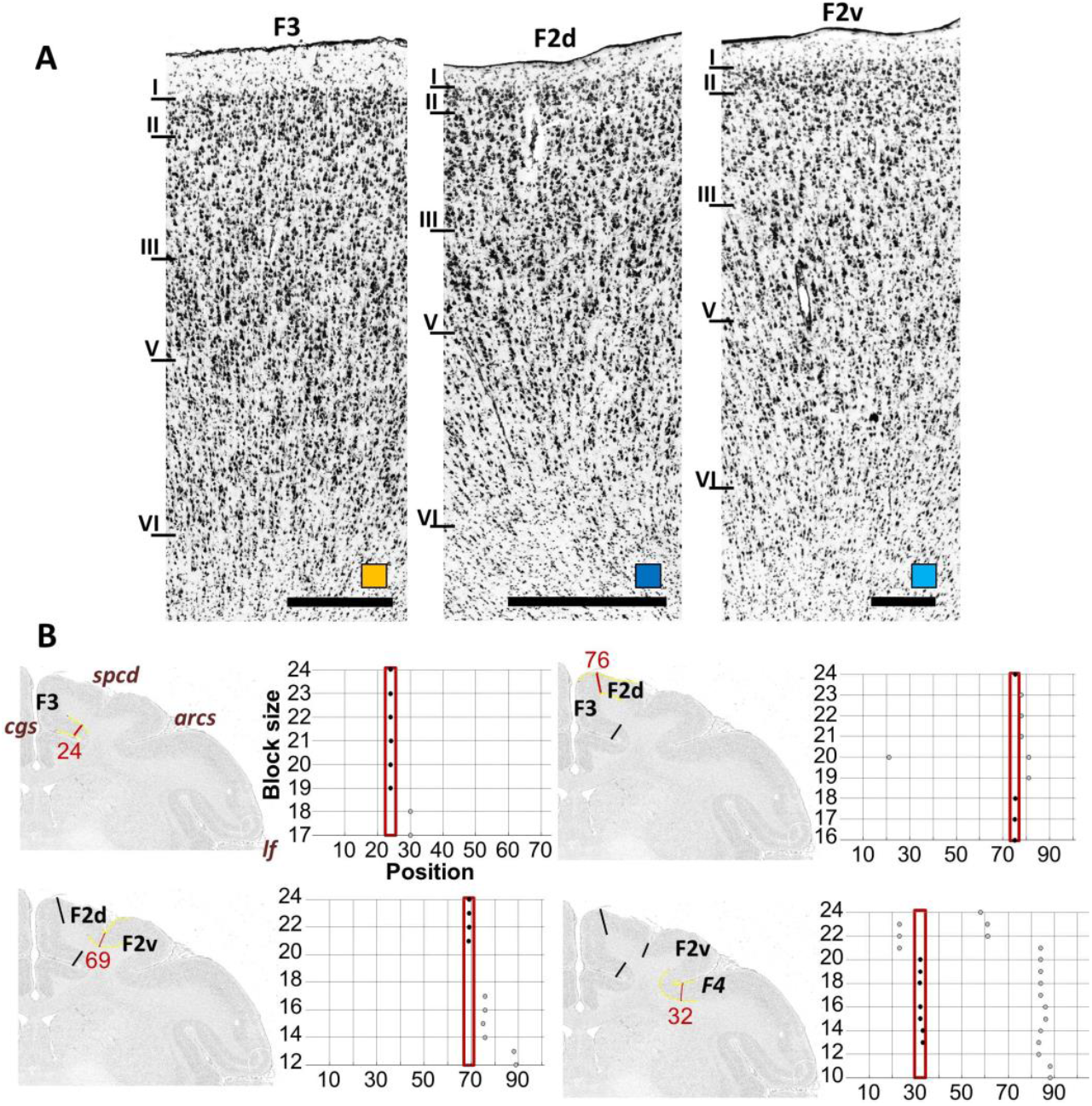
(A) Cytoarchitecture of caudal medial premotor area F3, as well as of the subdivisions of caudal dorsolateral premotor area F2 (F2d and F2v). Colored square over the scale bar indicates the color used in Figure 2 to code the area in question. (B) Quantitative analysis of cytoarchitectonic borders. The position of each border verified by the statistical analysis of Mahalanobis distances is highlighted by a dark red line (and corresponding profile index) on the GLI-image, and the corresponding dot plot (depicted to the right of the GLI-image) reveals that the location of significant maxima in the distance function (indicated by each dot) does not depend on the block size, but remains constant over large block size intervals (highlighted by the red frame). Roman numerals indicate cortical layers. Scale bars 1 mm. cgs – cingulate sulcus, ias –inferior arcuate branch, lf – lateral fissure, sas – superior arcuate branch.

**Figure 5.**
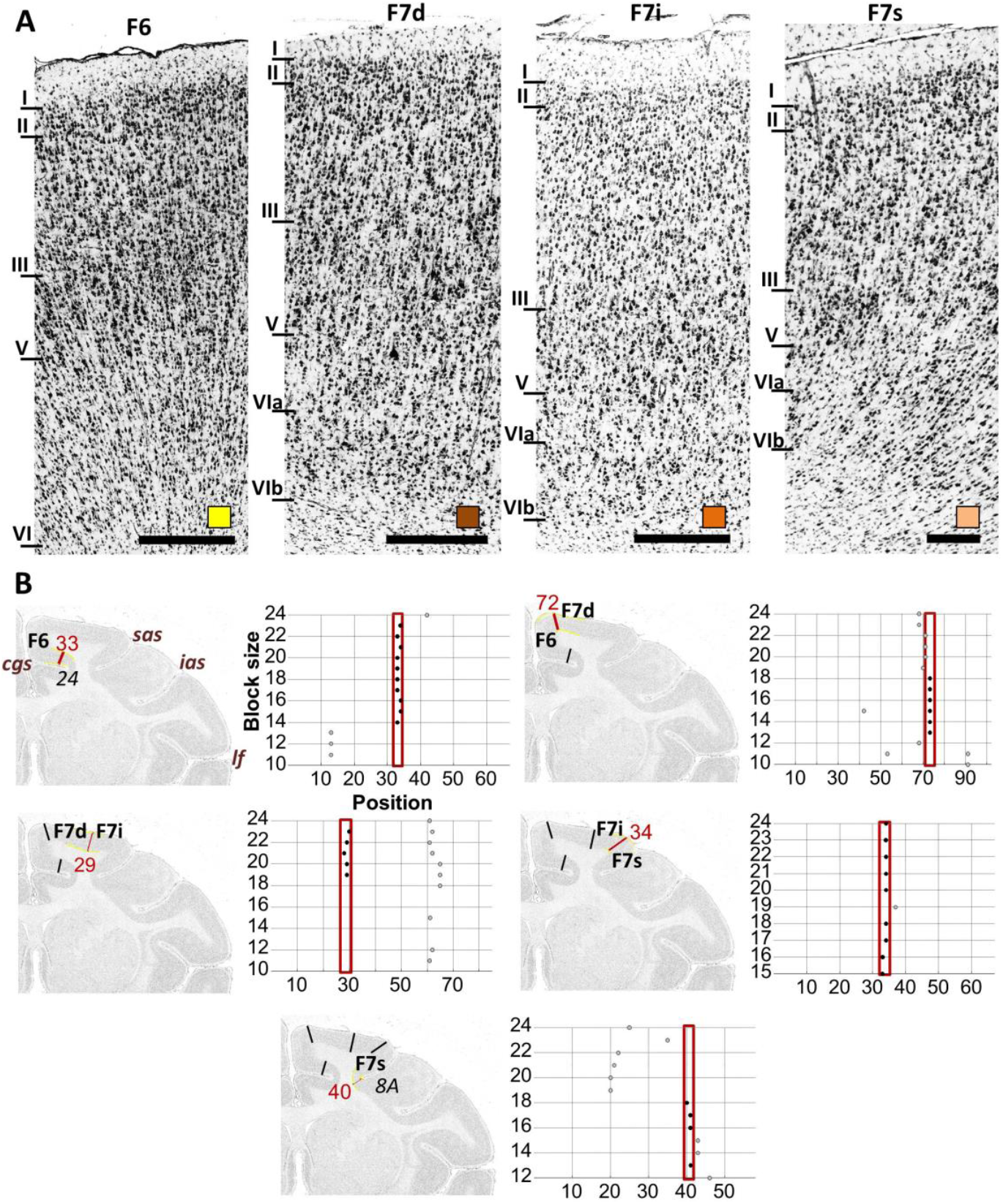
(A) Cytoarchitecture of rostral medial premotor area F6, as well as of the subdivisions of rostral dorsolateral premotor area F7 (F7d, F7i and F7s). Colored square over the scale bar indicates the color used in Figure 2 to code the area in question. (B) Quantitative analysis of cytoarchitectonic borders. The position of each border verified by the statistical analysis of Mahalanobis distances is highlighted by a dark red line (and corresponding profile index) on the GLI-image, and the corresponding dot plot (depicted to the right of the GLI-image) reveals that the location of significant maxima in the distance function (indicated by each dot) does not depend on the block size, but remains constant over large block size intervals (highlighted by the red frame). Roman numerals indicate cortical layers. Scale bars 1 mm. cgs – cingulate sulcus, ias – inferior arcuate branch, lf – lateral fissure, sas – superior arcuate branch.

Area F3 (Fig. 4A) can be distinguished from area F6 (Fig. 5A) by the numerous conspicuously large pyramids scattered throughout layer V of F3, but not of F6. Furthermore, F3 is characterized by an overall lower cell-packing density than F6, and this is particularly obvious in layers II, III and V. Thus, layer VI in F6 has an overall lower cell body density in regard to the superficial layers. Figures 4B and 5B provide an example of the statistical confirmation of visually identified cortical borders for areas F3 and F6, respectively.

#### 3.1.3 Dorsolateral premotor cortex

We identified five premotor areas on the dorsal convexity of the hemisphere: areas F2d and F2v (Fig. 4), which abuts the primary motor cortex, as well as areas F7d, F7i and F7s (Fig. 5), that are delimited rostrally and ventrally, at the fundus of the *sas*, by prefrontal area 8. The *spcd* serves as a partial landmark to identify the border between F2d and F2v, since it does not always cover the entire rostro-caudal extent of these two areas due to intersubject and interhemispheric differences in length (e.g., compare left and right hemispheres in Fig. 2). The *arcs* constitutes a reliable macroscopic landmark only to identify for the rostral portion of the border between F2v and the ventrolateral premotor areas (Fig. 2). F2v extends into the most caudal portion of the *sas* and close to its fundus is replaced by granular prefrontal cortex (Fig. 2).

Layer II of area F2d is slightly thinner but more densely packed than that of medially abutting F3. Furthermore, layer V of F2d is thinner than that of F3, though it seems more prominent due to the presence of cell aggregates (Fig. 4A). Layer V of F2d is also more prominent than that of F2v. Indeed, in F2v cell body packing density of layer V is comparable to that of the surrounding layers, making F2v appear less laminar than F2d. Finally, layer II is wider in F2v than in F2d (Fig. 4A). Statistical confirmation of visually identified cortical borders between F2 subdivisions is shown in Figure 4B.

Dorsal premotor area F7 can be clearly distinguished from neighboring areas by the subdivision of its layer VI into a pale, cell-sparse VIa and a cell-dense, darkly stained Vib (Fig. 5A). Furthermore, differences within layers VIa and VIb also enabled the definition of three subdivisions within this area: F7d is located on the most dorsal aspect of the hemisphere and is followed laterally by intermediate area F7i, that occupies the rest of the dorsal surface above *sas*, and by ventral area F7s located on the dorsal bank of the *sas* (Fig. 2). Areas F7d and F7i have an evidently sublaminated layer VI, which is not as apparent in F7s (Fig. 5A). Sublamina VIa is much wider in F7d than in F7i. Area F7s can also be distinguished from the other two areas by its denser layer II (Fig. 5A). The observer-independent analysis also confirmed the existence of these novel subdivisions of area F7 as presented in the exemplary section shown in Fig. 5B.

#### 3.1.4. Ventrolateral premotor cortex

We identified three areas occupying the posterior portion of the ventrolateral premotor cortex, i.e. areas F4d, F4v and F4s (Fig. 6), and three further areas in its rostral part, i.e. areas F5d, F5v and F5s (Fig. 7). Areas F4d and F4v are delimited caudally by primary motor area 4p and rostrally by premotor areas F5d and F5v (Fig. 2). Area F4s is located on the ventral wall of the *arcs*, where it reaches the sulcal fundus, and area F5s occupies the outer half of the ventral wall of *ias* (Fig. 2), where it neighbors granular prefrontal cortex.

**Figure 6.**
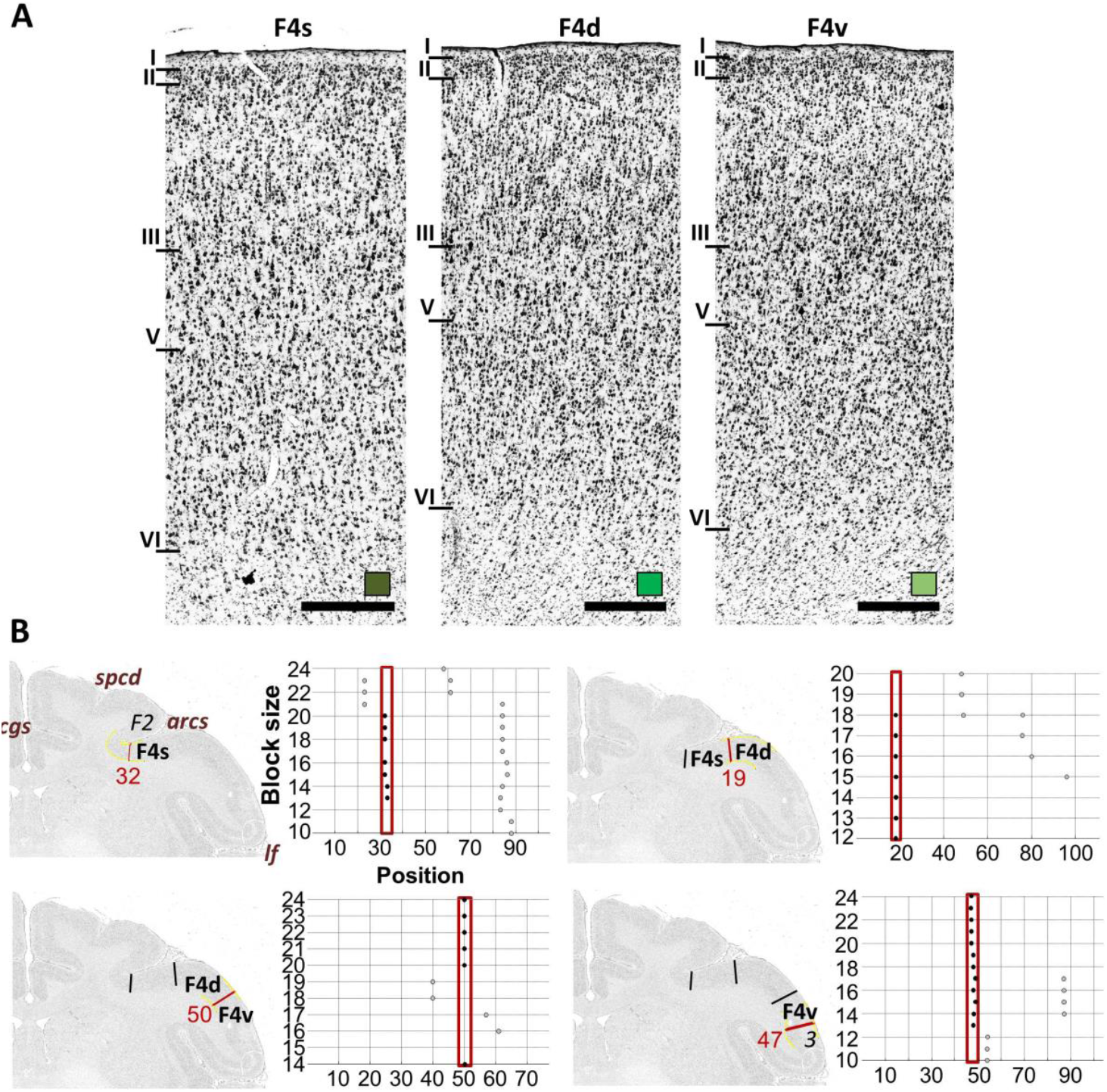
(A) Cytoarchitecture of the subdivisions of caudal ventrolateral premotor area F4 (F4s, F4d and F4v). Colored square over the scale bar indicates the color used in Figure 2 to code the area in question. (B) Quantitative analysis of cytoarchitectonic borders. The position of each border verified by the statistical analysis of Mahalanobis distances is highlighted by a dark red line (and corresponding profile index) on the GLI-image, and the corresponding dot plot (depicted to the right of the GLI-image) reveals that the location of significant maxima in the distance function (indicated by each dot) does not depend on the block size, but remains constant over large block size intervals (highlighted by the red frame). Roman numerals indicate cortical layers. Scale bars 1 mm. arcs – spur of the arcuate sulcus, cgs – cingulate sulcus, lf – lateral fissure, spcd – superior precentral dimple.

**Figure 7.**
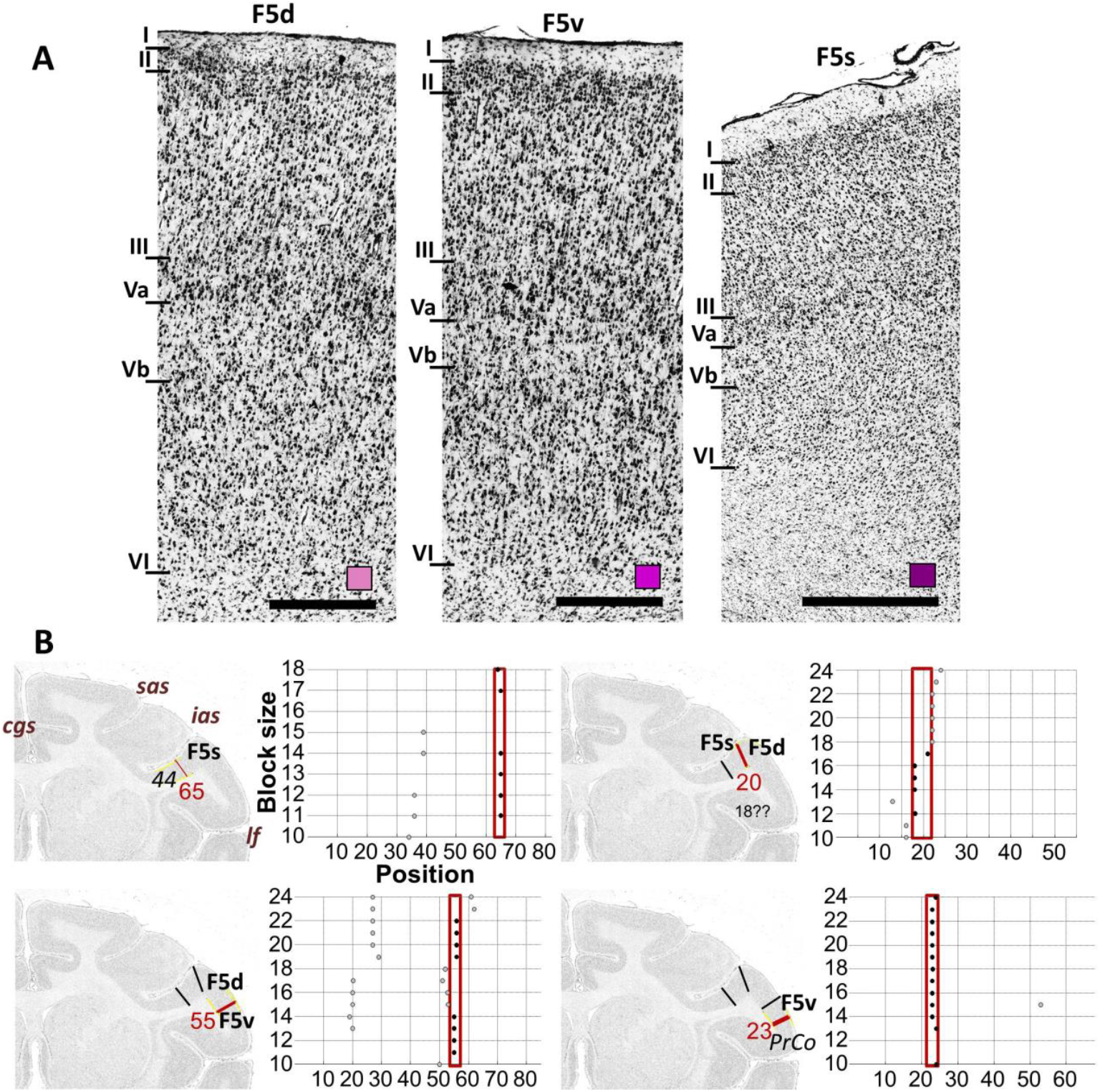
(A) Cytoarchitecture of the subdivisions of rostral ventrolateral premotor area F5 (F5s, F5d and F5v). Colored square over the scale bar indicates the color used in Figure 2 to code the area in question. (B) Quantitative analysis of cytoarchitectonic borders. The position of each border verified by the statistical analysis of Mahalanobis distances is highlighted by a dark red line (and corresponding profile index) on the GLI-image, and the corresponding dot plot (depicted to the right of the GLI-image) reveals that the location of significant maxima in the distance function (indicated by each dot) does not depend on the block size, but remains constant over large block size intervals (highlighted by the red frame). Roman numerals indicate cortical layers. Scale bars 1 mm cgs – cingulate sulcus, ias – inferior arcuate branch, lf – lateral fissure, sas – superior arcuate branch.

The cytoarchitecture of F4 (Fig. 6A) can be easily distinguished from that of neighboring areas due to the absence of Betz cells, as in the primary motor cortex, and of a sublamination of layer V, as in F5 (Fig. 7A). Three distinct subdivisions could be defined: F4s (sulcal), F4d (dorsal) and F4v (ventral; Fig. **6**). Layer II and upper layer III of F4s present a characteristic columnar organization that can’t be recognized in the lateral subdivisions F4d or F4v (Fig. 6A). Furthermore, F4d and F4v have smaller pyramids than F4s, and they are particularly small and densely packed in F4v. Finally, the border between layer VI and the white matter is sharper in F4s than in lateral areas F4d or F4v. In contrast, F4d has a noticeable columnar organization in layer VI, thus this layer blends gradually with the white matter, whereas F4v had slightly clearer border with the white matter, similar to F4s. Additionally, layer V of F4d has a lower cell packing density and is much wider than that of F4v, whereas layer I is thinner in F4v than in F4d (Fig. 6A). Newly defined borders within area F4 were also confirmed by the observer-independent analysis as presented in the exemplary section shown in Fig. 6B.

Area F5s has a prominent layer Va with a high cell packing density, and scattered medium-sized pyramids in layer Vb, which is much thinner than in the lateral subdivisions F5d and F5v (Fig. 7A). In F5s there is no distinct border between layers V and VI, but layers II and III can be clearly distinguished from each other. The main difference between F5 and neighboring prefrontal area 44 on the inner half of the ventral wall of *ias*, is the lack of an inner granular layer IV in the former area. Laterally neighboring area F5d is characterized by darkly stained small-sized pyramids with a horizontal organization in the lower part of layer III, and prominent medium-sized pyramids in layer V. Areas F5d and F5v also present a subdividable layer V, but in F5v border between Vb and VI is clearer than in F5d. Moreover, F5v lacks the horizontal organization in the lower part of the layer III (Fig. 7A). An example of section depicting the confirmation of border positions by the observer-independent analysis is presented in Fig. 7B.

### 3.2 Receptor architecture

We characterized the regional and laminar distribution patterns of 13 different receptor types in each cytoarchitectonically identified area by means of receptor profiles extracted perpendicularly to the cortical surface (Tab. 2) and identified significant differences in mean densities (i.e., averaged over all cortical layers) for specific receptors between bordering areas (Supplementary Fig. 1).

**Supplementary Figure 1.**
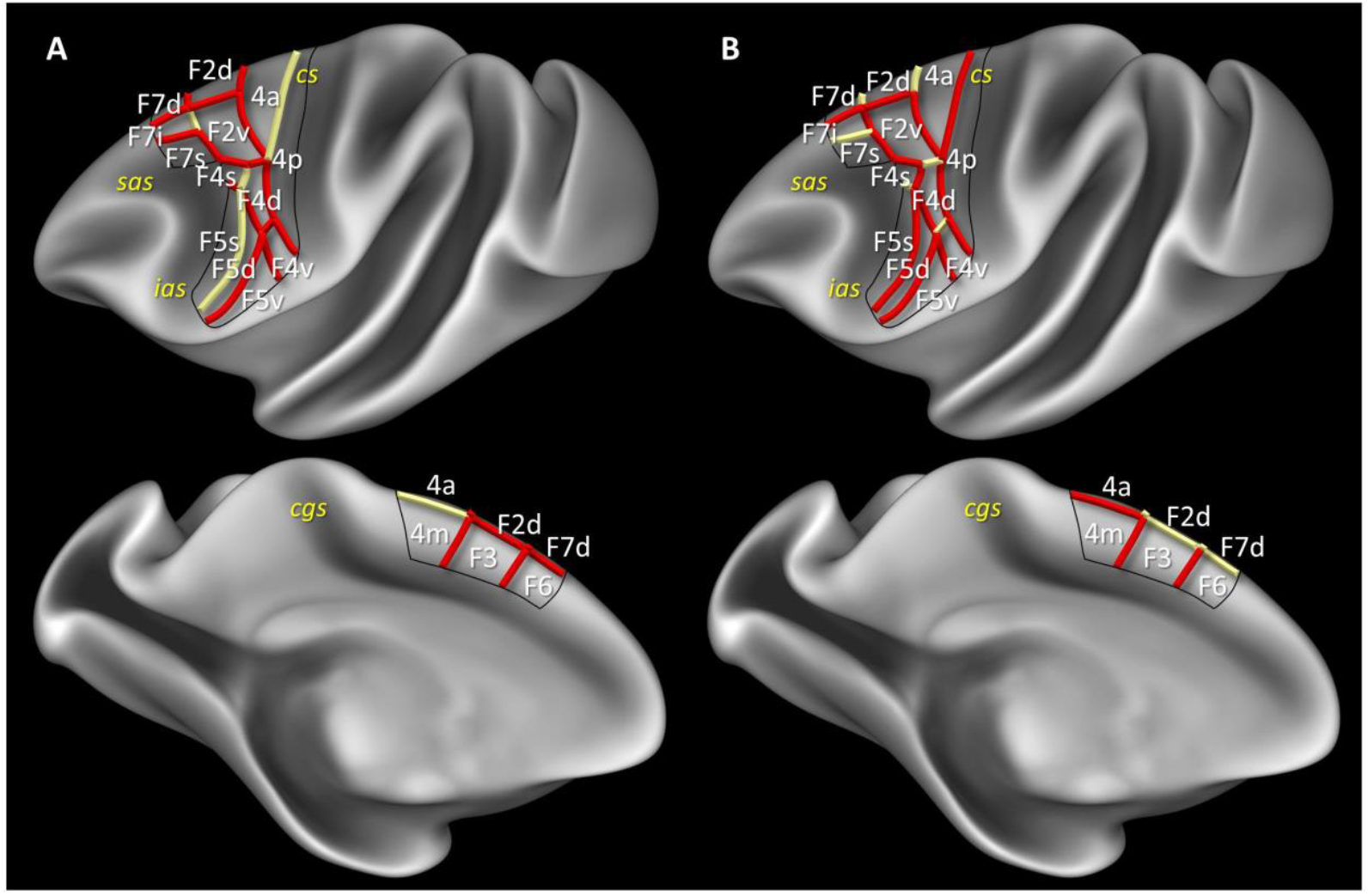
Summary scheme of the neurochemical (receptor) and functional (connectivity) correlation among neighboring motor and premotor areas, projected onto the lateral and medial views of the Yerkes19 surface (Donahue et al., 2016). Borders marked with red show significant differences in either the receptor (A) or the functional connectivity (B) analysis. Grey identifies borders with no significant differences between areas.

**Table 2.** Absolute receptor densities (mean ±s.d.) in fmol/mg protein. PMC premotor cortex, SMA supplementary motor area.

|  | Primary motor |  |  | SMA | Pre-SMA | Caudal dorsolateral PMC |  | Rostral dorsolateral PMC |  |  | Caudal ventrolateral PMC |  |  | Rostral ventrolateral PMC |  |  |
| --- | --- | --- | --- | --- | --- | --- | --- | --- | --- | --- | --- | --- | --- | --- | --- | --- |
|  | 4a | 4p | 4m | F3 | F6 | F2d | F2v | F7d | F7i | F7s | F4d | F4v | F4s | F5s | F5d | F5v |
| **AMPA** | 301 | 289 | 334 | 377 | 440 | 368 | 345 | 338 | 341 | 409 | 344 | 449 | 342 | 563 | 777 | 748 |
| s.d. | 70 | 81 | 60 | 140 | 103 | 51 | 44 | 81 | 101 | 63 | 38 | 118 | 82 | 239 | - | 73 |
| **Kainate** | 292 | 255 | 324 | 709 | 723 | 671 | 575 | 597 | 587 | 568 | 424 | 662 | 424 | 664 | 683 | 776 |
| s.d. | 75 | 76 | 86 | 42 | 79 | 81 | 51 | 94 | 66 | 76 | 41 | 221 | 17 | 22 | 43 | 50 |
| **NMDA** | 936 | 832 | 924 | 896 | 966 | 825 | 673 | 723 | 743 | 856 | 709 | 1021 | 728 | 1176 | 1153 | 1455 |
| s.d. | 257 | 228 | 241 | 307 | 302 | 195 | 154 | 299 | 240 | 289 | 62 | 337 | 154 | 257 | 166 | 121 |
| **GABA<sub>A</sub>** | 781 | 724 | 795 | 885 | 1070 | 934 | 920 | 1015 | 880 | 1023 | 868 | 1349 | 881 | 1292 | 1198 | 1359 |
| s.d. | 178 | 237 | 148 | 340 | 128 | 252 | 203 | 141 | 71 | 64 | 133 | 367 | 103 | 70 | 137 | 200 |
| **GABA<sub>B</sub>** | 1116 | 1255 | 1084 | 1749 | 1839 | 1910 | 1442 | 1776 | 1512 | 1580 | 1476 | 2095 | 1522 | 2078 | 2065 | 2450 |
| s.d. | 18 | 37 | 133 | 100 | 67 | 78 | 44 | 135 | 159 | 226 | 264 | 97 | 136 | 412 | 298 | 328 |
| **GABA<sub>A</sub>/BZ** | 1606 | 1489 | 1597 | 1554 | 1492 | 1804 | 1429 | 1892 | 1475 | 1470 | 1639 | 1931 | 1632 | 1802 | 1983 | 2087 |
| s.d. | 621 | 432 | 581 | 256 | 173 | 67 | 130 | 241 | 255 | 148 | 430 | 243 | 308 | 300 | 269 | 200 |
| **M<sub>1</sub>** | 488 | 461 | 417 | 815 | 861 | 877 | 808 | 831 | 718 | 807 | 562 | 747 | 456 | 766 | 705 | 849 |
| s.d. | 88 | 174 | 134 | 265 | 304 | 217 | 221 | 215 | 244 | 284 | 198 | 428 | 33 | 440 | 389 | 381 |
| **M<sub>2</sub>** | 85 | 83 | 88 | 188 | 180 | 177 | 171 | 206 | 160 | 165 | 100 | 135 | 133 | 146 | 146 | 168 |
| s.d. | 72 | 60 | 83 | 35 | 44 | 44 | 30 | 44 | 36 | 32 | 24 | 49 | 38 | 33 | 58 | 96 |
| **M<sub>3</sub>** | 563 | 540 | 534 | 564 | 563 | 583 | 534 | 583 | 545 | 579 | 472 | 578 | 492 | 707 | 646 | 736 |
| s.d. | 52 | 15 | 40 | 152 | 212 | 144 | 122 | 172 | 194 | 179 | 69 | 32 | 86 | 215 | 218 | 197 |
| **$\alpha_1$** | 352 | 334 | 359 | 489 | 479 | 512 | 448 | 440 | 421 | 422 | 386 | 502 | 377 | 469 | 467 | 504 |
| s.d. | 17 | 18 | 16 | 68 | 86 | 58 | 70 | 76 | 78 | 86 | 17 | 2 | 91 | 78 | 42 | 34 |
| **$\alpha_2$** | 141 | 166 | 141 | 333 | 319 | 362 | 284 | 287 | 292 | 307 | 258 | 326 | 235 | 363 | 348 | 420 |
| s.d. | 53 | 52 | 54 | 43 | 41 | 43 | 50 | 42 | 34 | 55 | 37 | 96 | 8 | 53 | 28 | 17 |
| **5HT<sub>1A</sub>** | 194 | 204 | 215 | 566 | 565 | 606 | 393 | 430 | 410 | 425 | 273 | 422 | 320 | 513 | 510 | 611 |
| s.d. | 51 | 25 | 25 | 71 | 134 | 88 | 60 | 60 | 90 | 90 | 7 | 12 | 73 | 140 | 146 | 144 |
| **5HT<sub>2</sub>** | 275 | 282 | 250 | 342 | 368 | 362 | 352 | 378 | 354 | 366 | 317 | 345 | 314 | 379 | 360 | 361 |
| s.d. | 57 | 53 | 48 | 63 | 83 | 75 | 87 | 76 | 88 | 75 | 89 | 56 | 64 | 67 | 35 | 48 |

Although not all receptors show each areal border, and not all borders are equally clearly defined by all receptor types, nevertheless, if a border was detected by at least five (or sometimes by all) receptor types, and this happened at a comparable position in at least three neighboring rostro-caudal levels, we confirmed the existence of a cytoarchitectonically identified border.

The cytoarchitectonically identified subdivisions of the primary motor cortex are revealed by differences in the laminar distribution patterns (Fig. 8, Supplementary Fig. 2) and mean absolute densities (Tab. 2) of multiple receptors. Although there are no significant differences in mean receptor densities among subdivisions of area 4, the border between areas 4m and 4a is clearly revealed by the higher kainate and α_1_ receptor densities in the infragranular layers of the former area as well as by the higher NMDA, but lower M_1_ and M_3_ densities in the supragranular layers (Fig. 8, Supplementary Fig. 2). Additionally, area 4p contained a higher GABA_B_ receptor density in the infragranular layers than areas 4a and 4m (Fig. 8, Supplementary Fig. 2).

**Figure 8.**
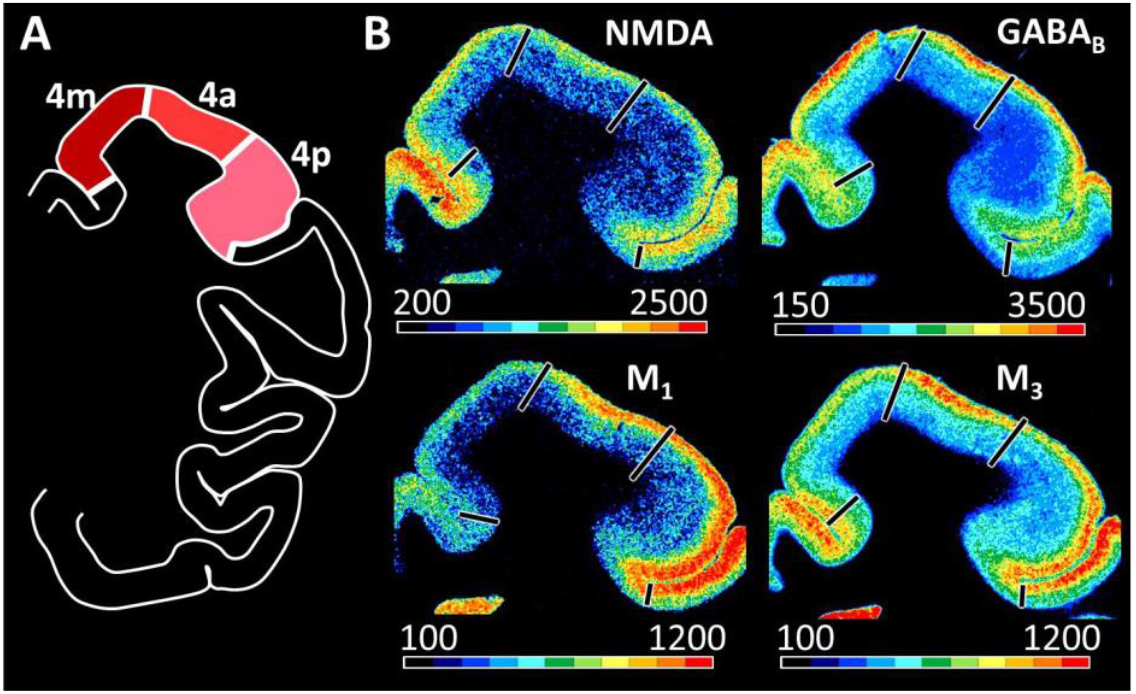
(A) Schematic drawing of a coronal section processed for receptor labeling at the level of the primary motor cortex showing the position of its medial (4m;), dorsolateral (4a;) and sulcal (4p;) subdivisions. Areal color coding as in Fig. 2. (B) Exemplary sections depicting the distribution of NMDA, GABA_B_, M_1_ and M_3_ receptors. Lines represent borders between defined premotor areas. Scale bars code for receptor densities in fmol/mg protein. Distribution patterns of all 13 receptors are shown in Supplementary Figure 1.

The subdivisions of area 4 can be distinguished from their rostrally neighboring areas (i.e. F3, F2d, F2v, F4d and F4v; Supplementary Fig. 1) by their significantly lower receptor concentrations in almost all examined receptors, especially in α_2_ and 5HT_1A_ receptors.

**Supplementary Figure 2.**
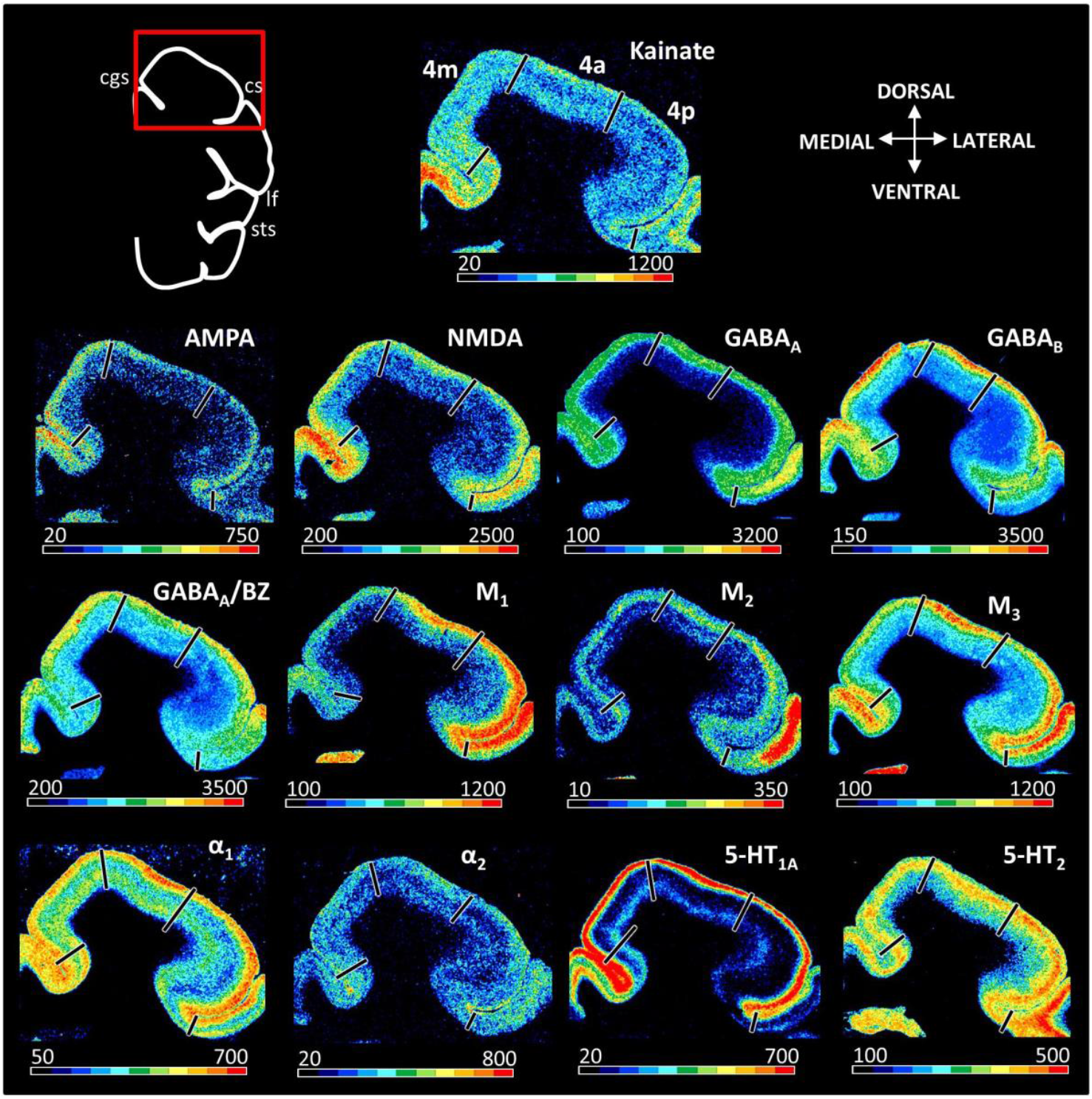
Neighboring coronal sections processed for receptor autoradiography and color coded to visualize the regional and laminar distribution patterns of all 13 examined receptor types in the macaque primary motor cortex. Color bars code for receptor densities in fmol/mg protein. cgs – cingulate sulcus, cs – central sulcus, lf – lateral fissure, sts – superior temporal sulcus.

Figures 9, 10 and Supplementary Figs. 3 and 4 display the receptor distribution patterns of medial premotor areas and of their bordering areas. As revealed by statistical analysis of mean areal densities for medial premotor areas (F3 and F6), inter-area differences were significantly indicated only by the 5-HT_2_ receptor, with higher levels in F6 compared to F3. Furthermore, GABA_A_/BZ binding site densities are higher in the superficial layers, conversely α_1_ receptor densities are higher in the deep layers of F3 than in those of F6 (Supplementary Figs. 3 and 4). Area F3 could be distinguished from area F2d most clearly by 5HT_2_ receptor. In contrast, rostral medial area F6 has higher concentration levels for almost all receptors, except for M_2_, M_3_ and 5HT_2_ as well as GABA_A_/BZ binding sites, when compared to dorsally bordering area F7d. Thus, we found significantly higher levels of kainate, NMDA and 5HT_1A_ receptors, but lower of GABA_A_/BZ binding sites, in F6 than in F7d.

**Figure 9.**
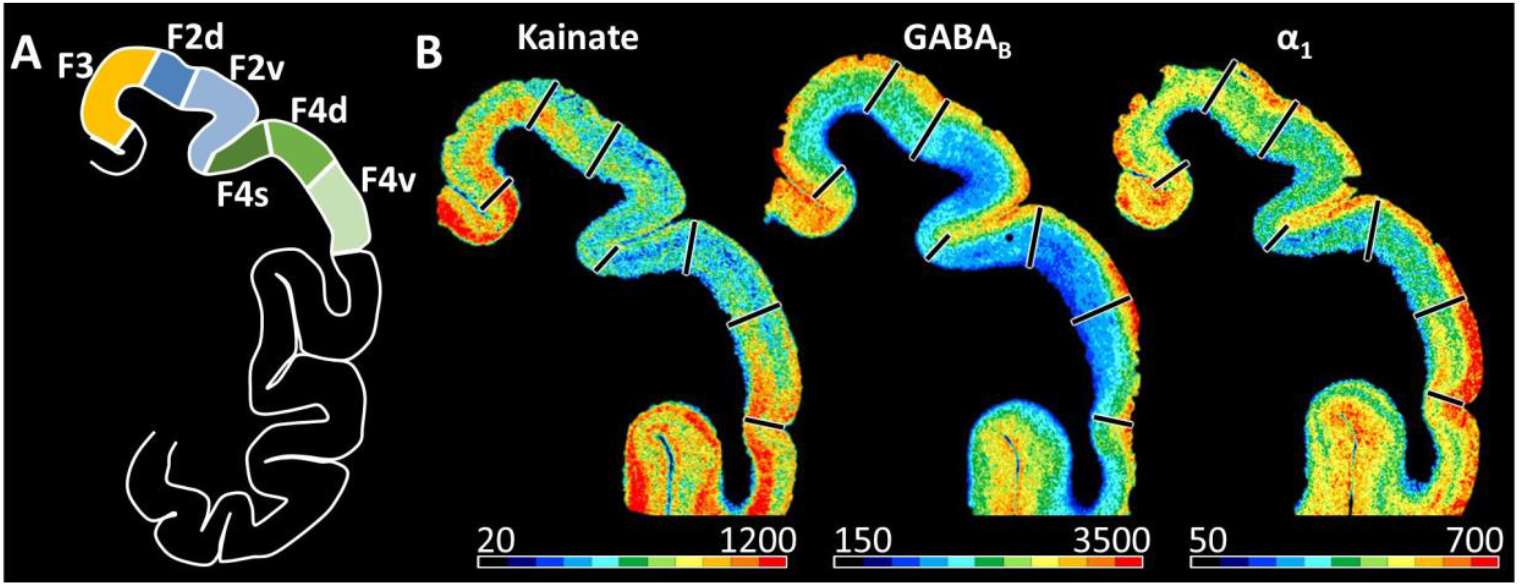
(A) Schematic drawing of a coronal section processed for receptor labeling at the level of the posterior premotor region showing the position of medial (F3), dorsolateral (F2d and F2v) and ventrolateral (F4s, F4d and F4v) premotor areas. Areal color coding as in Fig. 2. (B) Exemplary sections depicting the distribution of kainate, GABA_B_ and α_1_ receptors. Lines represent borders between defined premotor areas. Scale bars code for receptor densities in fmol/mg protein. Distribution patterns of all 13 receptors are shown in Supplementary Figures 2 and 4.

**Figure 10.**
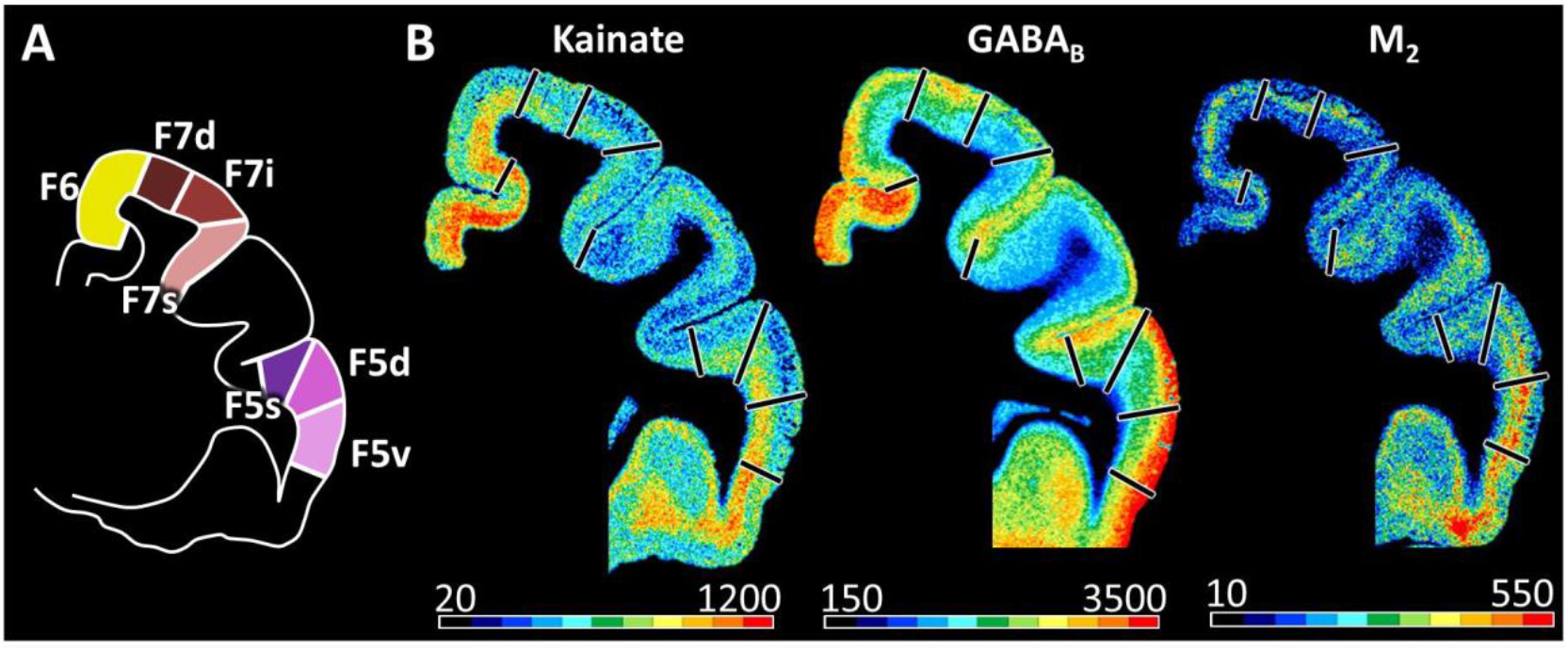
(A) Schematic drawing of a coronal section processed for receptor labelling at the level of the anterior premotor region showing the position of medial (F6), dorsolateral (F7d, F7i and F7s) and ventrolateral (F5s, F5d and F5v) premotor areas. Areal color coding as in Fig. 2. (B) Exemplary sections depicting the distribution of kainate, GABA_B_ and M_2_ receptors. Lines represent borders between defined premotor areas. Scale bars code for receptor densities in fmol/mg protein. Distribution patterns of all 13 receptors are shown in Supplementary Figures 3 and 5.

Regarding caudolateral premotor areas, the absolute densities of most receptor types decreased when moving from F2d through F2v to F4s and then increased when moving further ventrally into F4v (Tab. 2).

As shown in Fig. 9 and Supplementary Fig. 3, F2d has significantly higher densities of all receptors except AMPA, GABA_A_, 5HT_2_, and M_2_ compared to F2v. Furthermore, the infragranular layers of F2d presented higher kainate, NMDA, GABA_A_/BZ, M_1_, α_1_ and 5-HT_1A_ densities than those of F2v, whereas the supragranular layers of F2d presented higher AMPA, α_1_ and α_2_, but lower GABA_A_/BZ densities than those of F2v (Fig. 9, Supplementary Fig. 3). In contrast to the numerous significant differences between subdivisions of F2, only a few significant differences were found when compared to their bordering premotor areas, i.e. F7d, F7i, F7s and F4s. Here, F2d showed significantly higher concentrations for the kainate, α_1_, α_2_ and 5HT_1A_ receptors and lower concentrations for the M_2_ receptor as compared with F7d. Significantly lower densities of NMDA receptor were found in F2v than in F7s. No significant differences in receptor densities are found between F7i and F2v (Supplementary Fig. 1).

**Supplementary Figure 3.**
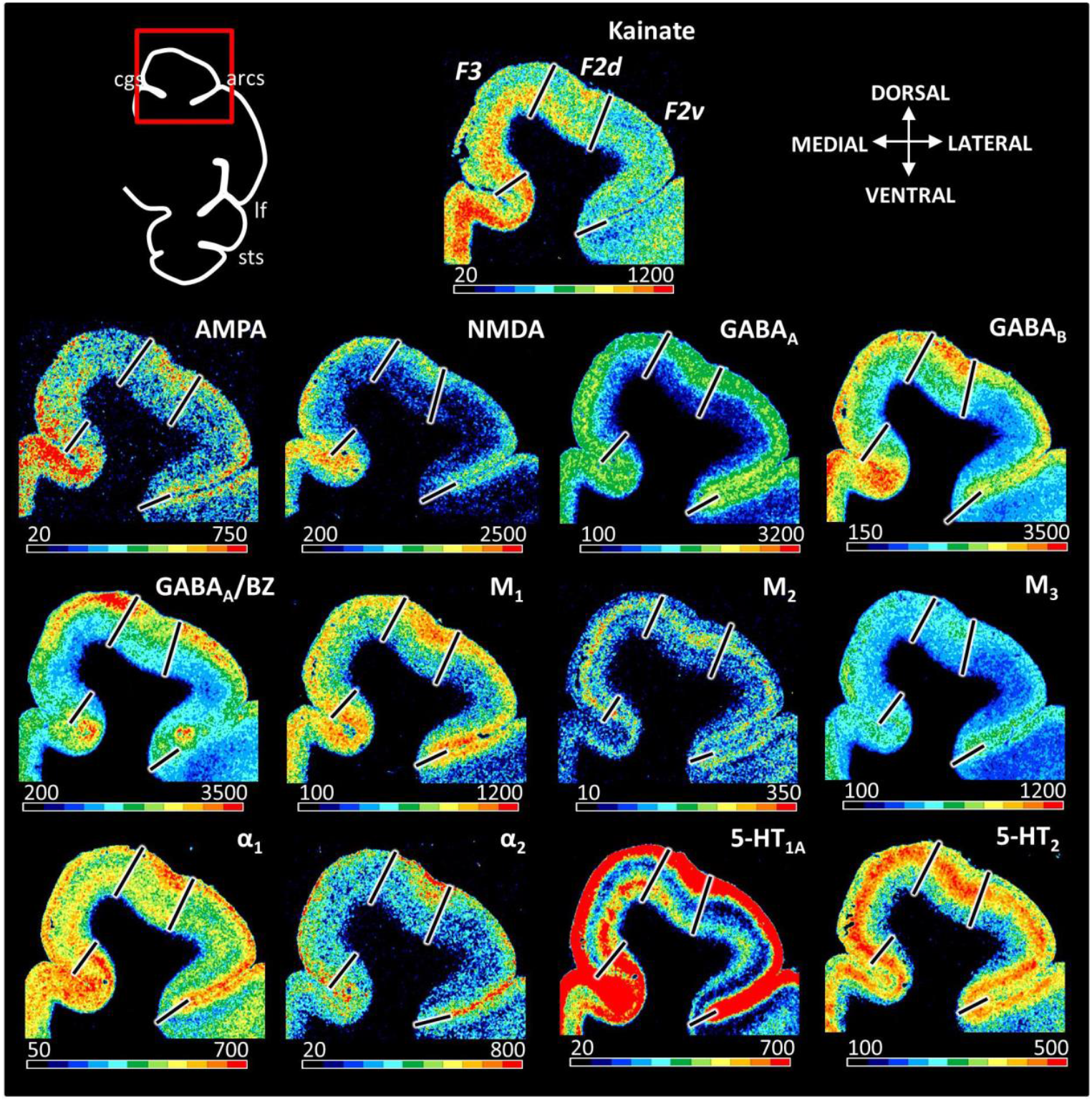
Neighboring coronal sections processed for receptor autoradiography and color coded to visualize the regional and laminar distribution patterns of all 13 examined receptor types in the macaque dorso-caudal premotor areas. Color bars code for receptor densities in fmol/mg protein. arcs – spur of the arcuate sulcus, cgs – cingulate sulcus, lf – lateral fissure, sts – superior temporal sulcus.

Although F4s and F4d are cytoarchitectonically different, they present comparable receptor fingerprints, and no significant differences are found between these two areas regarding their receptor densities. Layer-specific differences between F4d and F4v were found mainly in the infragranular layers, which presented lowest densities in F4d and highest in F4v (Fig. 9, Supplementary Fig. 5). Areas F4d and F4v differ significantly in their NMDA and GABA_A_ receptor densities, which are higher in the latter subdivision of F4. Additionally, the subdivisions of area F4 can be easily distinguished from their rostrally F5 areas (F5s, F5d and F5v) by their lower receptor concentrations in all examined receptors, especially in glutamatergic (AMPA, kainate, NMDA) receptors, which were statistically significant.

Receptor distribution patterns in the rostrolateral premotor areas also confirmed the position of cytoarchitectonically identified borders. Area F7d presented significantly higher GABA_A_/BZ, M_1_, M_2_ and 5HT_2_ densities than F7i, especially in the supragranular layers (Fig. 10, Supplementary Fig. 4). As revealed by statistical analysis of mean areal densities, the most obvious differences between areas F7i and F7s appeared in GABA_A_ receptor. Regarding the laminar distribution, the densities of GABAergic and α_2_ receptors were lower in the supragranular layers of F7i than in those of F7s, whereas the opposite holds true for NMDA receptors. Additionally, kainate and 5HT_1A_ receptor densities in the infragranular layers of F7i were higher than those of F7d or F7s (Fig. 10, Supplementary Fig. 4).

**Supplementary Figure 4.**
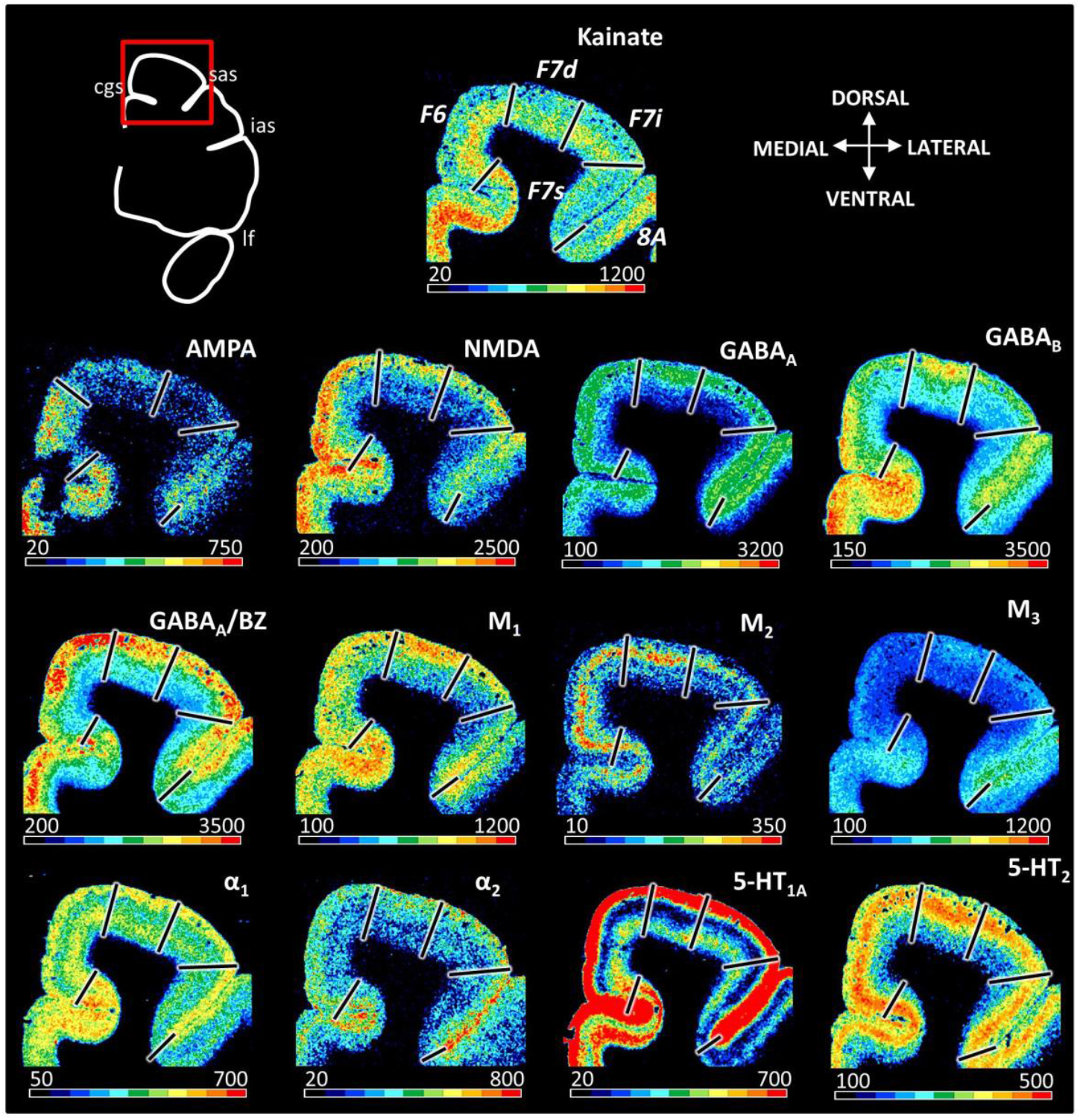
Neighboring coronal sections processed for receptor autoradiography and color coded to visualize the regional and laminar distribution patterns of all 13 examined receptor types in the macaque rostro-dorsal premotor areas and in the rostrally adjacent prefrontal area 8A. Color bars code for receptor densities in fmol/mg protein. arcs – spur of the arcuate sulcus, cgs – cingulate sulcus, ias – inferior arcuate branch, lf – lateral fissure, sas – superior arcuate branch.

Within subdivisions of area F5, a gradual increase in the densities of kainate, NMDA, GABA_B_, M_2_, α_1_ and 5-HT_1A_ receptors is noticed when moving from F5s through F5d to F5v (Fig. 10, Supplementary Fig. 6). These changes were more prominent in the infragranular layers for M_2_, α_1_ and 5HT_1A_ receptors and in the supragranular layers for NMDA and GABA_B_ receptors (Fig. 10, Supplementary Fig. 6). Area F5s presents lower AMPA and GABA_A_/BZ, but higher GABA_B_ densities than does F5d (Tab. 2). Area F5d contains significantly lower NMDA, GABA_B_, M_3_, α_2_ and 5HT_1A_ receptor densities than does F5v (Tab. 2).

**Supplementary Figure 5.**
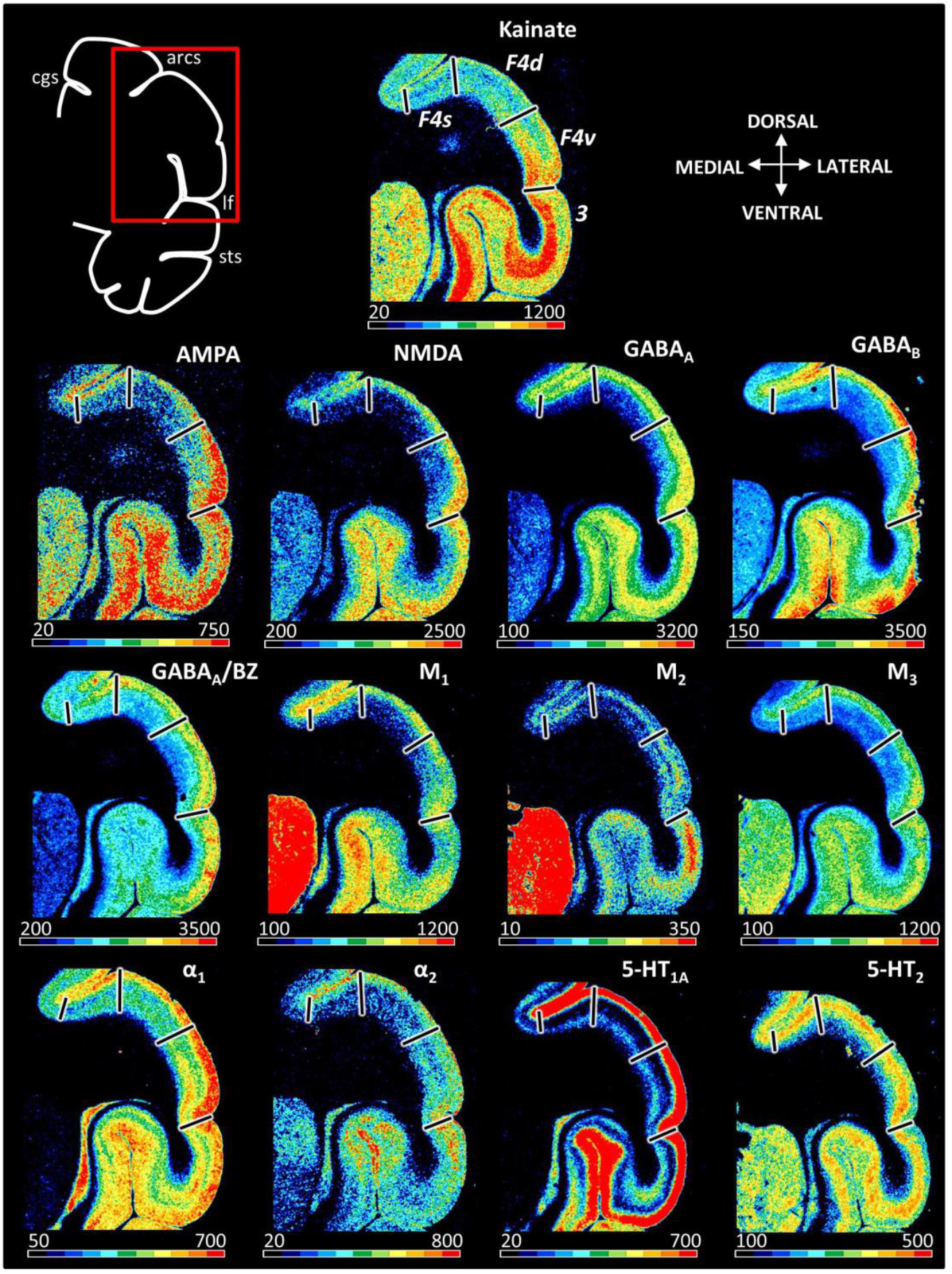
Neighboring coronal sections processed for receptor autoradiography and color coded to visualize the regional and laminar distribution patterns of all 13 examined receptor types in the macaque ventro-caudal premotor areas and in caudally adjacent somatosensory area 3. Color bars code for receptor densities in fmol/mg protein. arcs – spur of the arcuate sulcus, cgs – cingulate sulcus, lf – lateral fissure, sts – superior temporal sulcus.

**Supplementary Figure 6.**
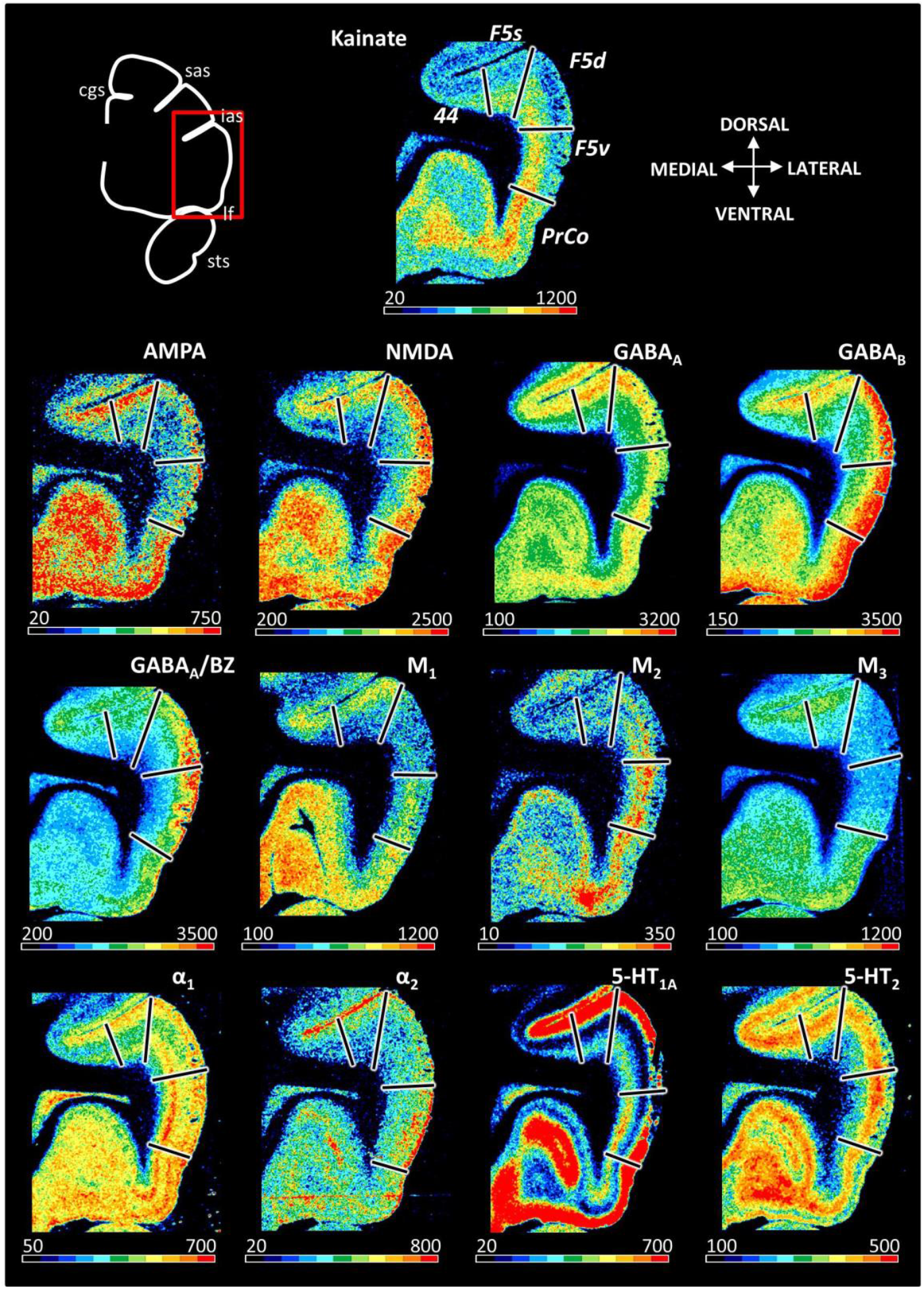
Neighboring coronal sections processed for receptor autoradiography and color coded to visualize the regional and laminar distribution patterns of all 13 examined receptor types in the macaque rostro-dorsal premotor areas, rostrally adjacent prefrontal area 44 and ventrally adjacent precentral opercular area (PrCo). Color bars code for receptor densities in fmol/mg protein. cgs – cingulate sulcus, ias – inferior arcuate branch, lf – lateral fissure, sas – superior arcuate branch, sts – superior temporal sulcus.

#### 3.2.1 Receptor fingerprints in stereotaxic space

Given the close relationship between the position of cortical borders and macroanatomical landmarks presented in the 2D flat map (Fig. 2), as well as the extremely low degree of interindividual variability of both features in the macaque brain, we were able to draw the relative position and extent of motor and premotor areas on the Yerkes19 surface using the Workbench software, and thus provide a spatial visualization of the parcellation scheme (Fig. 11) and of the differences in receptor densities throughout the monkey agranular frontal cortex (Fig. 11; Supplementary Fig. 7). These figures not only reveal the clear differences between the primary motor cortex (4m, 4a and 4p) and the premotor region, but also the existence of rostro-caudal and dorso-ventral trend within the premotor cortex. The mean densities of all receptor types are lower in the primary motor than in the premotor cortex. Areas F4s and F4d of the lateral premotor surface have generally lower receptor densities compared to the remaining premotor areas, and are thus more comparable to the primary motor areas.

**Figure 11.**
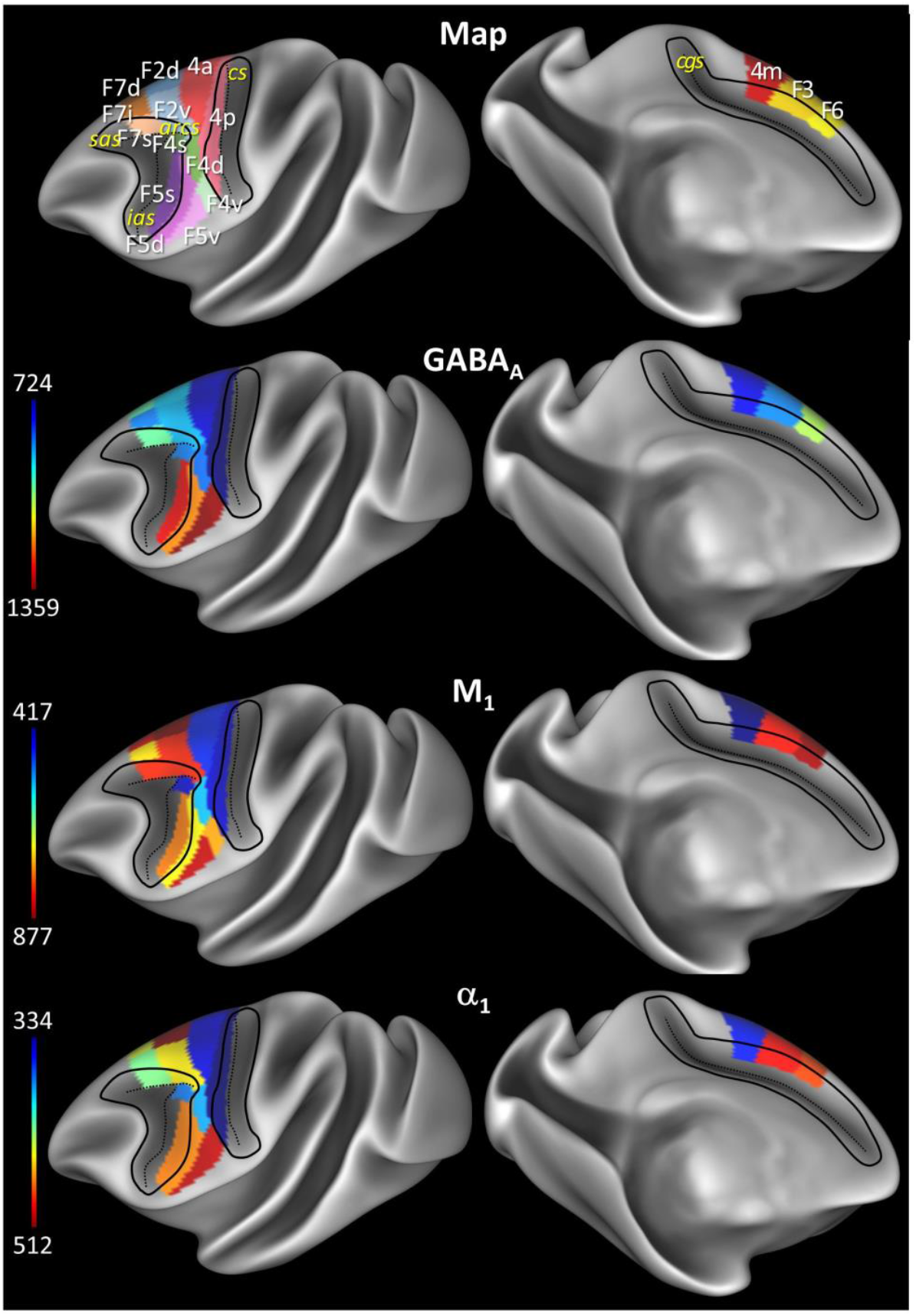
Position and extent of the motor and premotor areas on lateral and medial views of the Yerkes19 surface (Donahue et al., 2016). The mean receptor densities of three exemplary receptor types (α_1_, GABA_A_ and M_1_) have been projected onto the corresponding area. Color bars code for receptor densities in fmol/mg protein. The projections of all receptor types onto the Yerkes 19 surface are shown in Supplementary figure 6. The file coding for the densities of all 13 receptors in each area is provided as Supplementary data 1 made available to the neuroscientific community via the HBP and PRIME-DE repositories.

Within the lateral premotor region, α_1_ receptors show a rostro-caudal gradient, with caudal (F3, F2 and F4v) areas containing higher concentrations than rostral (F6, F7 and F5) ones (Fig. 11). Furthermore, the opposite trend has been observed for kainate, α_2_, 5HT_1A_ and 5HT_2_ receptors, these receptors have rather higher concentrations in the rostral premotor areas than caudal ones (Supplementary Fig. 7). The GABA_A_ and M_1_ receptors, on the other hand, present a clear dorso-ventral trend (Fig. 11). Similar trends are observed for AMPA, NMDA, GABA_B_, M_2_ and M_3_ receptors as well as GABA_A_/BZ binding sites (Supplementary Fig. 7). Most of the receptors show lower receptor densities in dorsal (subdivisions of areas F2 and F7) than in ventral (subdivisions of areas F4 and F5) areas, as presented for GABA_A_ in Fig. 11. The opposite trend holds true for M_1_ and M_2_ receptors (Fig. 11; Supplementary Fig. 7).

Areas on the medial surface (4m, F3 and F6) show for most receptors, i.e. AMPA, kainate, GABA_A_, GABA_B_, M_1_ and 5HT_2_, a clear rostro-caudal gradient in receptor densities, with highest concentrations found in rostral area F6 and lowest ones in caudal area 4m. The opposite trend is observed only in GABA_A_/BZ binding sites (Fig. 11; Supplementary Fig. 7). However, the rostro-caudal trend isn’t present in all receptors. Instead, area F3 has higher concentration levels of M_2_, α_1_ and α_2_ than surrounding areas 4m and F6, or lower levels, as in case of NMDA receptors, than areas 4m and F6 (Fig. 11; Supplementary Fig. 7). Finally, premotor areas F3 and F6 show no significant differences in M_3_ and 5HT_1A_ receptor concentration levels (Supplementary Fig. 7).

**Supplementary Figure 7.**
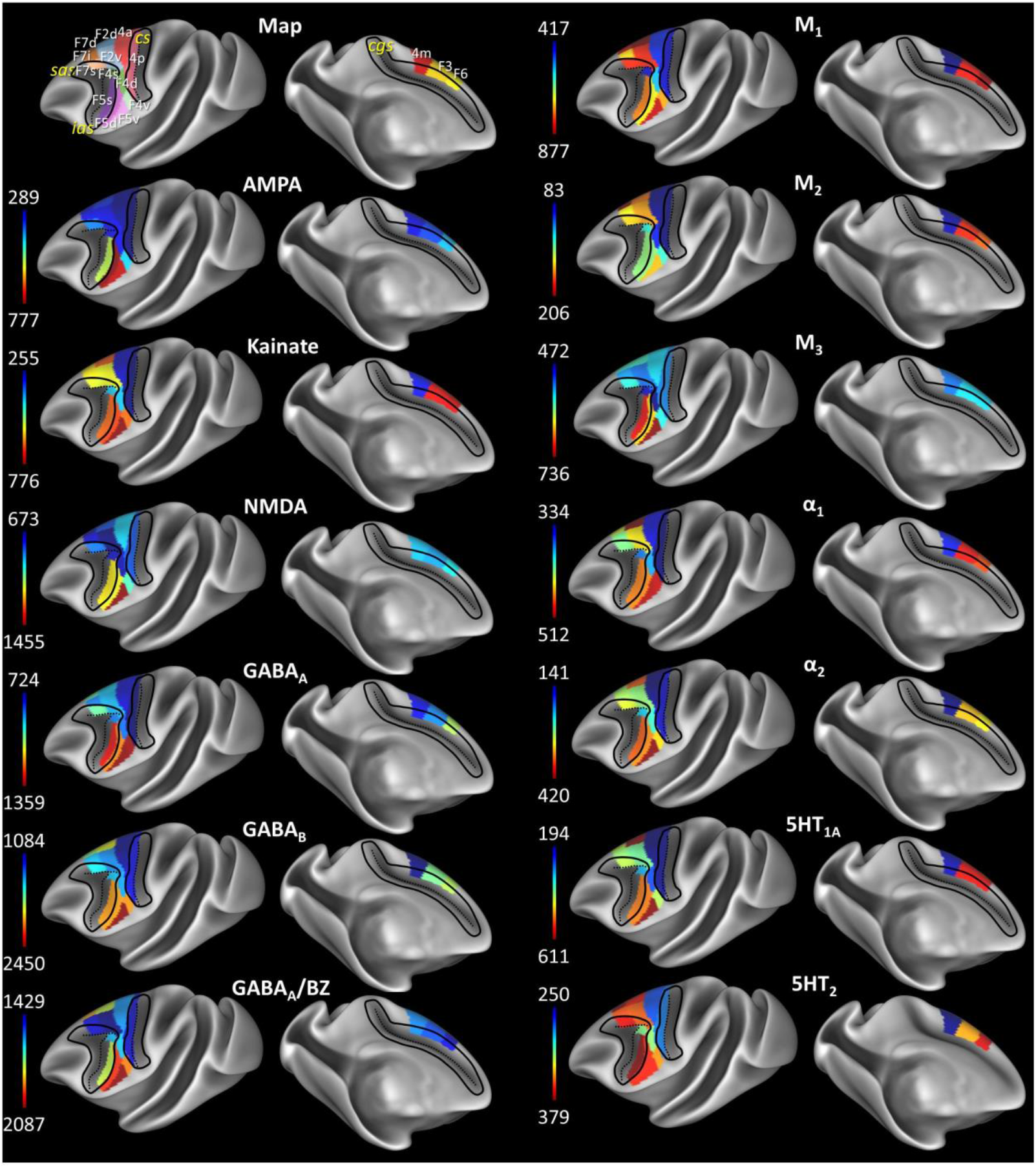
Position and extent of the areas in ROI and mean densities of 13 receptors projected onto the lateral and medial views of the Yerkes19 surface (Donahue et al., 2016). See map in Fig.11. Color bars code for receptor densities in fmol/mg protein. The file coding for the densities of all 13 receptors in each area is provided as Supplementary data 1

The novel cortical parcellation based on cytoarchitecture and receptor architecture will be made available on the Human Brain Project and BALSA neuroimaging sites, along with all receptor data used to make the receptor fingerprints.

#### 3.2.2 Cluster analyses of receptor fingerprints

Finally, for each area and subarea, mean receptor densities (averaged over all cortical layers) were visualized as a ‘receptor fingerprint’ (Supplementary Fig. 8), which illustrates the specific receptor balance of the identified area. Receptor fingerprints can vary in shape and size, and those of the primary motor cortex are obviously different when compared to those of the premotor areas due to the overall lower absolute receptor concentration values in areas 4a, 4p and 4m than in premotor areas.

**Supplementary Figure 8.**
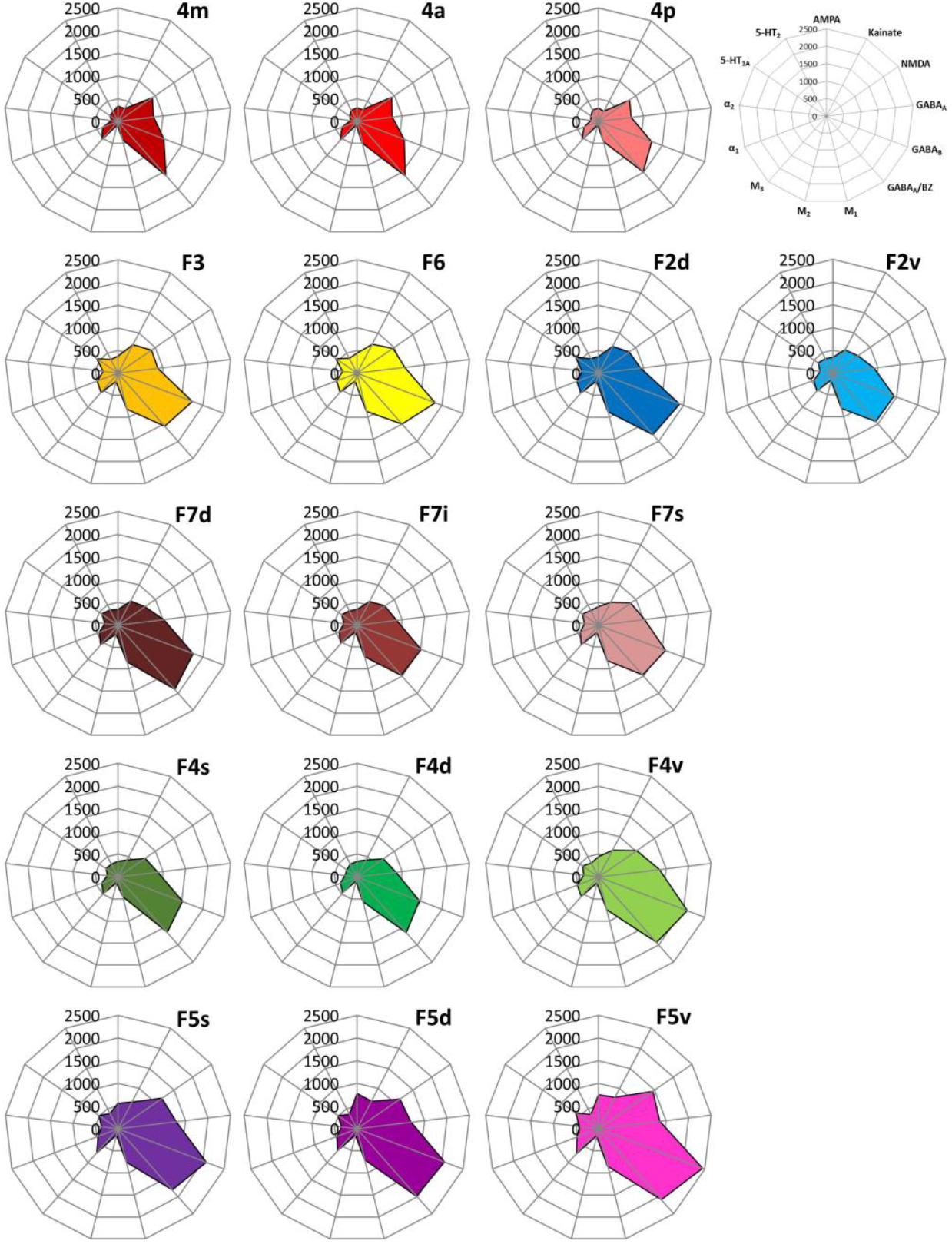
Receptor fingerprints of the examined areas for each subdivision Receptor fingerprints of the examined brain regions, i.e., polar coordinate plots depicting the absolute densities (in fmol/mg protein) of 13 receptors in each area. The positions of the different receptor types and the axis scaling are identical in all polar plots, and specified in the polar plot at the top right corner of the figure. The colored area represents the mean absolute receptor densities. Color coding as in Fig. 2.

A hierarchical cluster analysis was conducted in order to reveal (dis)similarities between receptor fingerprints of the macaque monkey motor and premotor areas, and the k-means clustering and elbow-analysis determined 3 as the optimal number of clusters. A fundamental branching separated a “caudal cluster” containing areas of the primary motor cortex, as well as areas F4d and F4s, from all other premotor areas (Fig. 12A). However, the three primary motor areas and the two premotor areas are located on separate branches within this cluster. Areas 4m and 4a are more similar to each other than to 4p, as revealed by the size and shape of their receptor fingerprints. In a second step further branching resulted in a clear segregation of ventral premotor areas, grouped into a “ventral cluster”, from those found on the dorsomedial and dorsolateral hemispheric surfaces, constituting a “dorsal cluster” (Fig. 12A). Interestingly, within the ventral cluster F4v is more similar to F5s and F5d than does F5v. Within the dorsal cluster, dorso-medial premotor areas F3 and F6 associate with the most dorsal portions of areas F2 and F7, i.e. F2d and F7d. In the principal component analysis, the 1^st^ principal component confirmed the segregation of the primary motor areas (4m, 4a and 4p) and the most dorsal subdivisions of ventral premotor area F4 (F4s and F4d) from the remaining premotor areas (vertical dashed line in Fig. 12B). Further, the 2^nd^ principal component separated the ventral from the dorsal premotor areas (horizontal dashed line in Fig. 12B).

**Figure 12.**
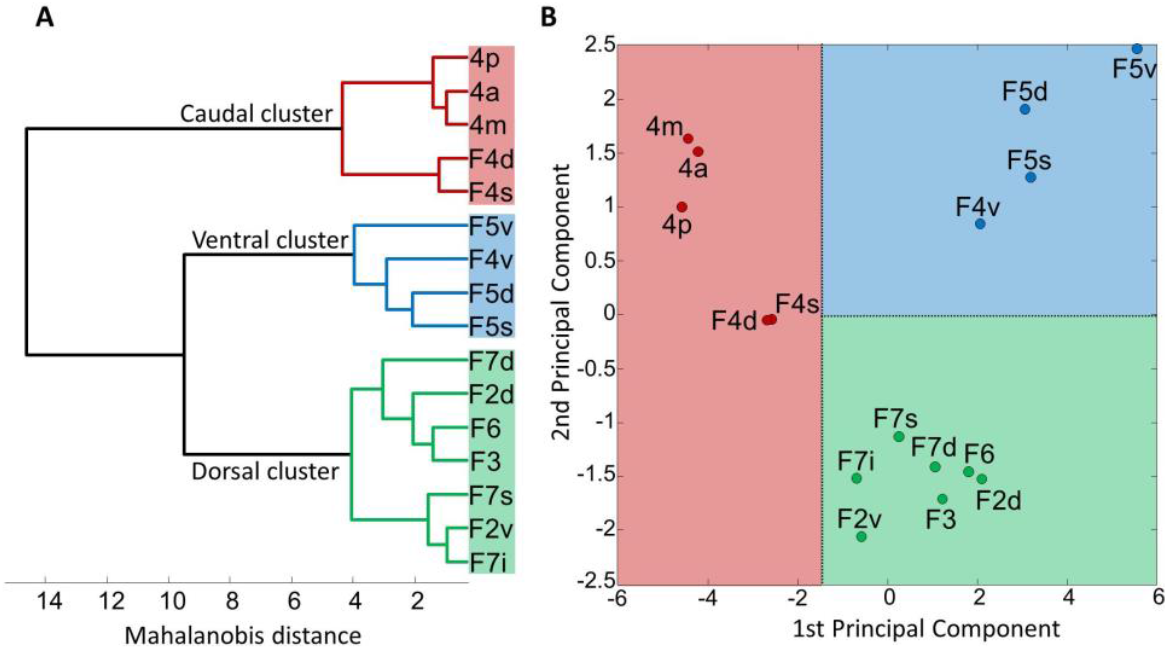
Hierarchical cluster (A) and principal component (B) analyses of the receptor fingerprints of macaque primary motor and premotor areas. K-means clustering and elbow analysis showed three as the optimal number of clusters.

### 3.3 Functional Connectivity analyses

Cortical areas not only have a unique cyto- and receptor architecture, but can also be characterized by their distinctive pattern of connectivity. Thus, we analyzed the functional connectivity of each of the identified motor and premotor areas with areas of prefrontal, cingulate, somatosensory and lateral parietal cortex and depicted the result as connectivity fingerprints (Fig. 13; Table 3; Supplementary Fig. 1) and as seed-to-vertex connectivity maps (Supplementary Fig. 9).

**Table 3.** Functional connectivity between motor and premotor areas defined in the present study and areas identified by resting state fMRI. PMC premotor cortex, SMA supplementary motor area.

|  | Primary motor |  |  | SMA | Pre-SMA | Caudal dorsolateral PMC |  | Rostral dorsolateral PMC |  |  | Caudal ventrolateral PMC |  |  | Rostral ventrolateral PMC |  |  |
| --- | --- | --- | --- | --- | --- | --- | --- | --- | --- | --- | --- | --- | --- | --- | --- | --- |
|  | 4a | 4p | 4m | F3 | F6 | F2d | F2v | F7d | F7i | F7s | F4d | F4v | F4s | F5s | F5d | F5v |
| **10** | 0.001 | 0.007 | -0.002 | 0.026 | 0.021 | 0.030 | 0.027 | 0.034 | 0.036 | 0.015 | 0.029 | 0.025 | 0.011 | 0.051 | 0.063 | 0.078 |
| **9** | 0.056 | 0.062 | 0.021 | 0.084 | 0.103 | 0.066 | 0.103 | 0.078 | 0.095 | 0.083 | 0.041 | 0.047 | 0.049 | 0.076 | 0.070 | 0.102 |
| **9/46d** | 0.222 | 0.222 | 0.150 | 0.310 | 0.317 | 0.227 | 0.327 | 0.217 | 0.317 | 0.339 | 0.212 | 0.136 | 0.293 | 0.305 | 0.175 | 0.192 |
| **46d** | 0.163 | 0.178 | 0.125 | 0.233 | 0.216 | 0.180 | 0.246 | 0.152 | 0.222 | 0.265 | 0.135 | 0.104 | 0.231 | 0.220 | 0.127 | 0.132 |
| **9/46v** | 0.086 | 0.119 | 0.143 | 0.204 | 0.206 | 0.138 | 0.205 | 0.192 | 0.212 | 0.170 | 0.207 | 0.232 | 0.200 | 0.345 | 0.287 | 0.263 |
| **46v** | 0.102 | 0.125 | 0.088 | 0.172 | 0.173 | 0.128 | 0.195 | 0.151 | 0.173 | 0.171 | 0.126 | 0.122 | 0.134 | 0.216 | 0.182 | 0.186 |
| **8m** | 0.261 | 0.295 | 0.263 | 0.351 | 0.293 | 0.267 | 0.355 | 0.231 | 0.298 | 0.482 | 0.229 | 0.184 | 0.408 | 0.330 | 0.150 | 0.147 |
| **8l** | 0.178 | 0.231 | 0.250 | 0.309 | 0.269 | 0.203 | 0.324 | 0.192 | 0.276 | 0.317 | 0.276 | 0.273 | 0.341 | 0.435 | 0.266 | 0.250 |
| **8r** | 0.221 | 0.255 | 0.223 | 0.357 | 0.331 | 0.246 | 0.360 | 0.248 | 0.326 | 0.382 | 0.231 | 0.205 | 0.375 | 0.384 | 0.226 | 0.210 |
| **45A** | 0.010 | 0.010 | 0.008 | 0.048 | 0.037 | 0.028 | 0.077 | 0.029 | 0.077 | 0.053 | 0.077 | 0.092 | 0.065 | 0.172 | 0.196 | 0.155 |
| **ProM** | 0.050 | 0.015 | 0.127 | 0.048 | 0.046 | 0.037 | 0.100 | 0.048 | 0.067 | 0.045 | 0.194 | 0.246 | 0.066 | 0.227 | 0.297 | 0.643 |
| **24c** | 0.090 | 0.095 | 0.080 | 0.159 | 0.165 | 0.132 | 0.159 | 0.139 | 0.163 | 0.158 | 0.104 | 0.089 | 0.131 | 0.168 | 0.124 | 0.134 |
| **32** | 0.049 | 0.055 | 0.037 | 0.114 | 0.092 | 0.080 | 0.112 | 0.070 | 0.104 | 0.098 | 0.072 | 0.062 | 0.083 | 0.131 | 0.097 | 0.102 |
| **1** | 0.211 | 0.202 | 0.435 | 0.212 | 0.146 | 0.158 | 0.182 | 0.104 | 0.111 | 0.147 | 0.213 | 0.190 | 0.183 | 0.166 | 0.097 | 0.113 |
| **2** | 0.131 | 0.099 | 0.530 | 0.119 | 0.086 | 0.096 | 0.144 | 0.075 | 0.063 | 0.130 | 0.341 | 0.450 | 0.185 | 0.267 | 0.273 | 0.308 |
| **3** | 0.234 | 0.282 | 0.662 | 0.261 | 0.165 | 0.183 | 0.222 | 0.104 | 0.130 | 0.243 | 0.245 | 0.281 | 0.273 | 0.225 | 0.145 | 0.163 |
| **5** | 0.318 | 0.299 | 0.331 | 0.314 | 0.233 | 0.263 | 0.332 | 0.170 | 0.203 | 0.312 | 0.170 | 0.093 | 0.302 | 0.204 | 0.073 | 0.101 |
| **7A** | 0.274 | 0.323 | 0.223 | 0.348 | 0.274 | 0.272 | 0.366 | 0.227 | 0.320 | 0.402 | 0.170 | 0.112 | 0.336 | 0.268 | 0.113 | 0.123 |
| **7B** | 0.123 | 0.163 | 0.388 | 0.208 | 0.170 | 0.124 | 0.169 | 0.135 | 0.116 | 0.176 | 0.291 | 0.293 | 0.226 | 0.310 | 0.195 | 0.195 |
| **7m** | 0.279 | 0.361 | 0.269 | 0.359 | 0.264 | 0.257 | 0.312 | 0.175 | 0.216 | 0.339 | 0.136 | 0.077 | 0.318 | 0.174 | 0.050 | 0.064 |
| **LIP** | 0.247 | 0.273 | 0.298 | 0.271 | 0.188 | 0.218 | 0.295 | 0.161 | 0.222 | 0.347 | 0.152 | 0.120 | 0.316 | 0.227 | 0.081 | 0.087 |
| **MIP** | 0.208 | 0.202 | 0.171 | 0.184 | 0.125 | 0.170 | 0.232 | 0.117 | 0.152 | 0.239 | 0.074 | 0.029 | 0.200 | 0.104 | 0.019 | 0.042 |
| **VIP** | 0.162 | 0.176 | 0.204 | 0.149 | 0.095 | 0.123 | 0.201 | 0.082 | 0.121 | 0.191 | 0.091 | 0.055 | 0.190 | 0.108 | 0.024 | 0.022 |

**Figure 13.**
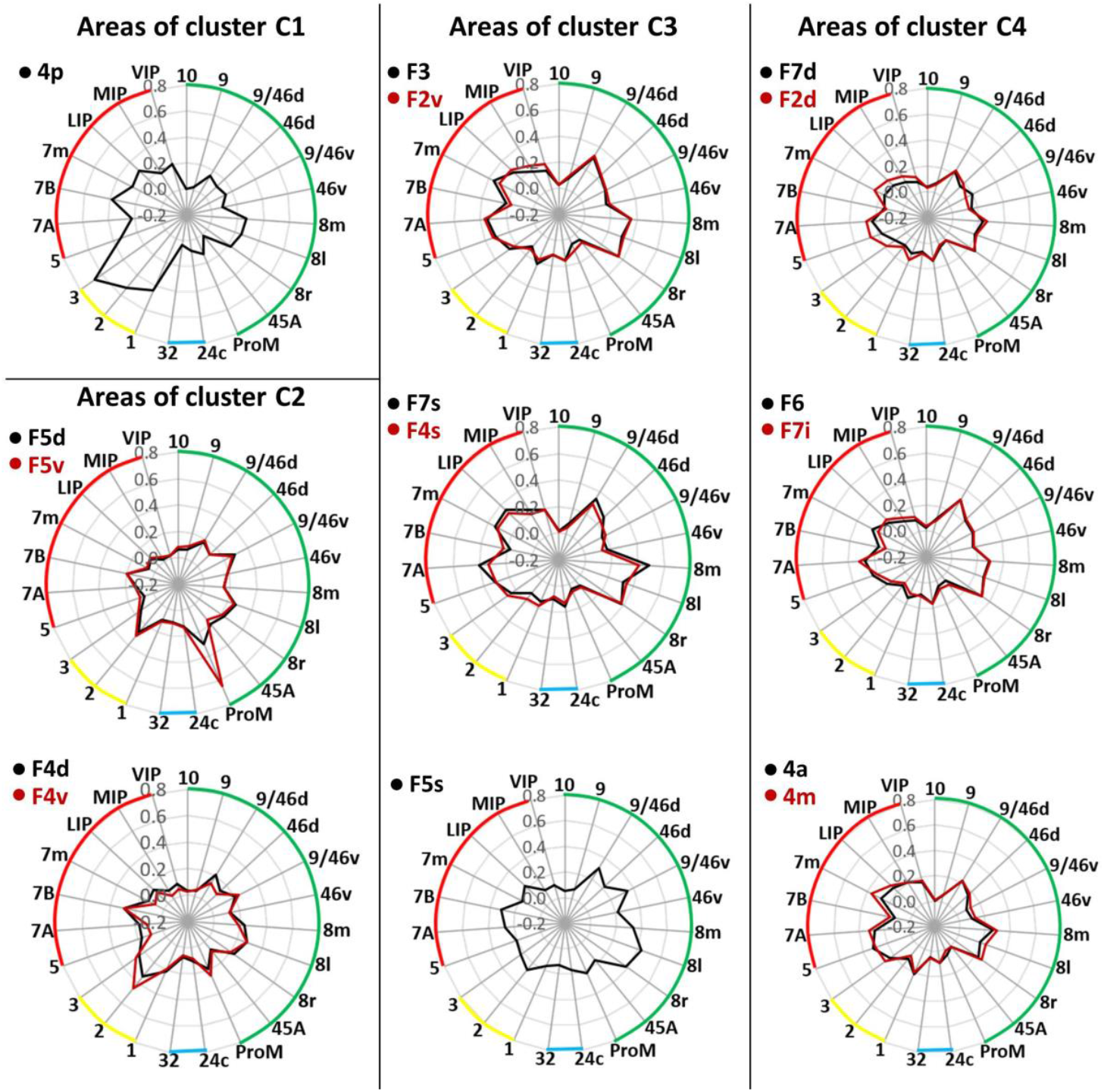
Connectivity fingerprints of each examined area and showing their connectivity strength to areas of the resting state network. Green codes for prefrontal areas, blue for cingulate, yellow for somatosensory and red for parietal areas. Nomenclature of targeted areas has is based on the Kennedy atlas (Markov et al., 2014), axis scaling is identical in all polar plots.

**Supplementary Figure 9.**
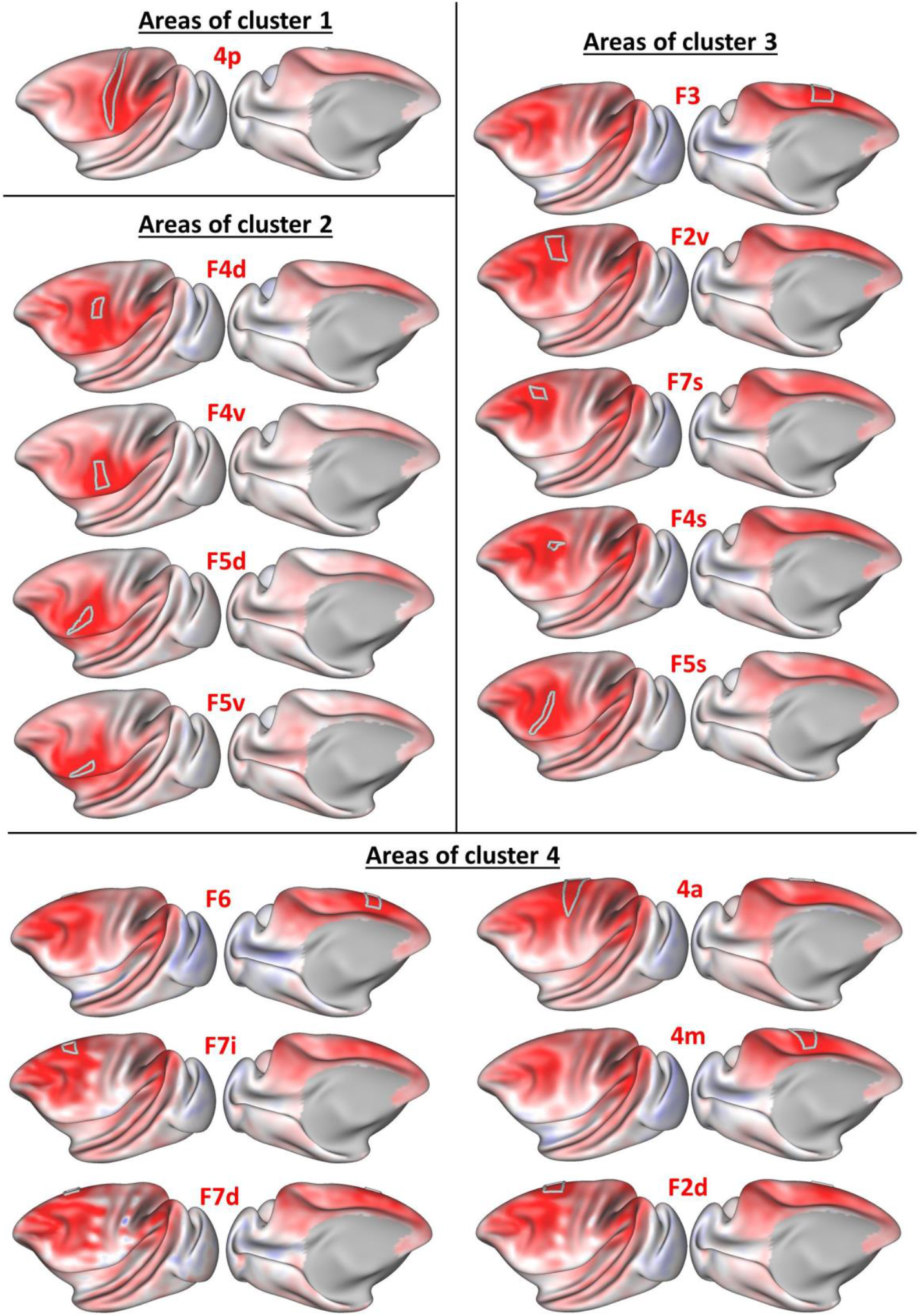
Seed-to-voxel connectivity maps resulting from the analysis of the functional connectivity between the cyto- and receptor architectonically distinct areas defined in the present study and cortical areas revealed by resting state fMRI analysis in the macaque monkey. Red regions present positive correlation, while blue regions show negative correlation.

Functional connectivity fingerprints of primary motor areas 4a and 4m differed significantly from each other, and, in particular, from that of primary motor area 4p, but resembled those of supplementary (F3) and pre-supplementary (F6) areas, as well as of dorsolateral premotor areas (F2d, F2v, F7d, F7i and F7s), and ventrolateral premotor area F4s (Fig. 13). The connectivity fingerprint of area 4p stood out by its particularly conspicuous strong connectivity with somatosensory areas, in particular with area 3, whereas the ventral premotor areas (F4d, F4v, F5s, F5d and F5v) had stronger connectivity to area 2. However, similarity of their fingerprints is displayed by connectivity with proisocortical motor area ProM (most notable in the connectivity fingerprint of F5v) and parietal area 7B, rather than 7A. Interestingly, the functional connectivity fingerprints of areas F2v and F7i are larger than those of their dorsally located counterparts (F2d and F7d, respectively; Fig. 13).

#### 3.2.2 Cluster analyses of functional connectivity fingerprints

Based on the functional connectivity fingerprints of the macaque monkey motor and premotor areas, a hierarchical cluster analysis was conducted, where the k-means clustering and elbow-analysis determined 4 as the optimal number of clusters.

The fundamental branching revealed clear differentiation of the primary motor subdivision 4p, which solely presented cluster 1 (C1, Fig. 14A) and ventral premotor areas (F5d, F5v, F4d and F4v) grouped into cluster 2 (C2, Fig. 14A) from the areas located on the dorsolateral and medial cortical surfaces. Further branching separated areas located within the arcuate sulcus (F2v, F4s, F7i and F4s) and medial area F3, which grouped into cluster 3 (C3, Fig. 14A), and areas arranged as cluster 4 (C4, Fig. 14A), i.e. dorsolateral areas (F7d, F7i and F2d) and medial area F6, grouped together with the primary motor subdivisions 4a and 4m.

**Figure 14.**
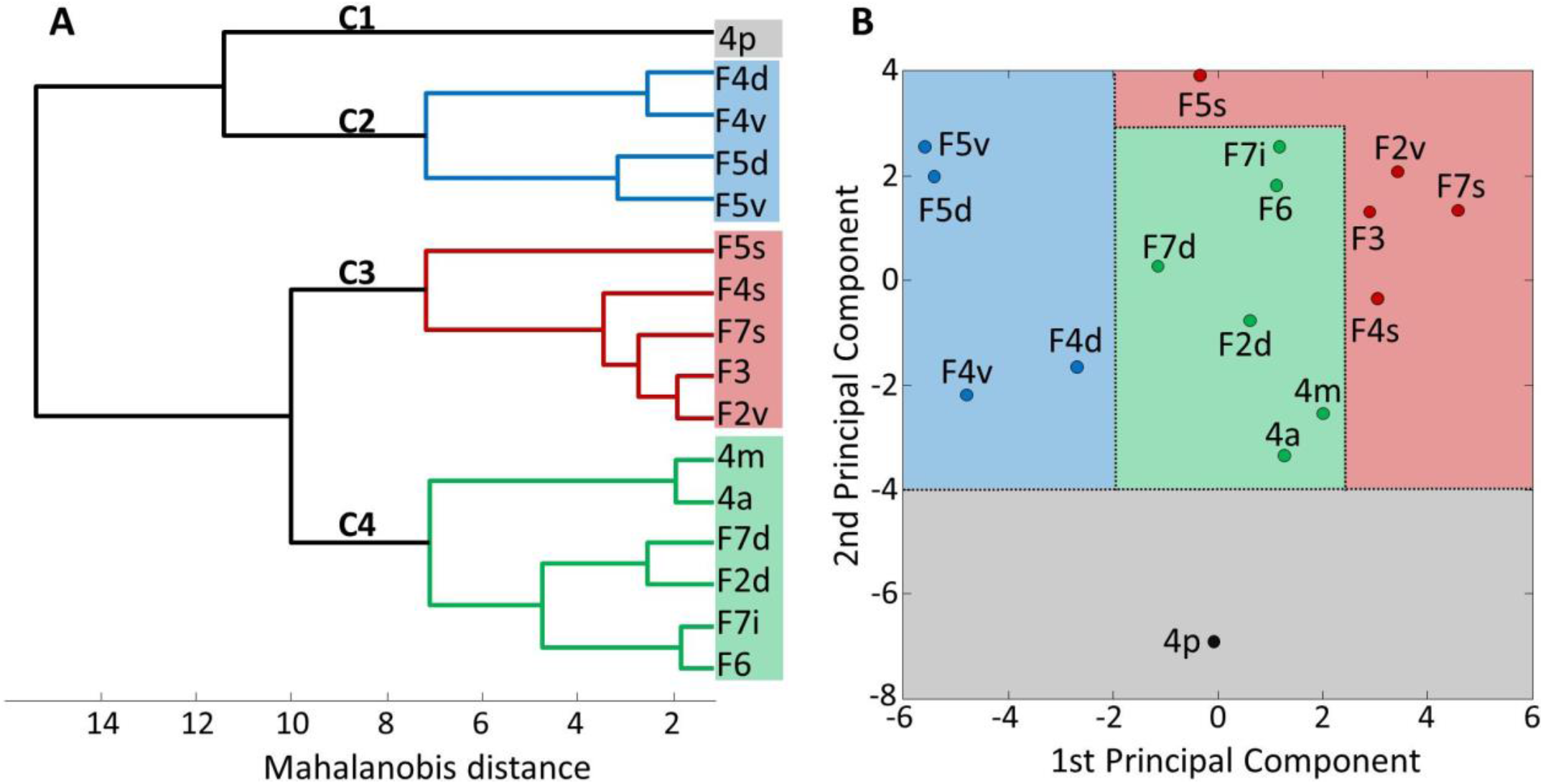
Hierarchical cluster (A) and principal component (B) analyses of the functional connectivity fingerprints of macaque primary motor and premotor areas. K-means clustering and elbow analysis showed four as the optimal number of clusters.

As for the principal component analysis, most notably, the 1^st^ principal component confirmed the segregation of the ventral premotor areas (F5d, F5v, F4d and F4v) from the remaining premotor and motor areas (vertical dashed line in Fig. 14B). Whereas the 2^nd^ principal component clearly set apart area 4p from the all the other cluster groups, emphasizing its unique connectivity pattern among motor and premotor areas (horizontal dashed line in Fig. 14B).

## 4. DISCUSSION

We conducted a multimodal and statistically testable analysis of the monkey agranular frontal cortex which resulted in the definition of 16 cyto- and receptor architectonically distinct areas. We identified three areas within the primary motor cortex (4a, 4p and 4m), and confirmed the existence of areas F3 and F6 (supplementary motor and pre-supplementary motor cortex, respectively). We also propose a novel parcellation scheme for lateral premotor areas F4 (divided into areas F4d, F4v, and F4s), F5 (divided into areas F5d, F5v, and F5s), and F7 (divided into areas F7d, F7i, and F7s). The identified areas were mapped to the Yerkes19 surface (Donahue et al., 2016) and the mean density of each of the 13 receptors in a given area was projected onto the corresponding area in the ensuing parcellation scheme. We then described the functional “connectivity fingerprint” of each of the newly defined areas. Thus, we provide for the first time a 3D atlas of macaque motor and premotor areas in stereotaxic space, which integrates information of their cytoarchitecture receptor architecture and functional connectivity. Furthermore, cluster analyses of the receptor fingerprints revealed a closer association of premotor areas F4d and F4s with primary motor areas than with the remaining premotor areas, as well as a segregation of ventral from dorsal premotor areas based on differences in their receptor fingerprints. Additionally, we analyzed the strength of the functional connectivity of each defined area with components of the resting state network. The functional connectivity fingerprints of areas 4a and 4m were found to be less similar to that of area 4p than to those of medial or dorsolateral premotor areas. Within the dorsolateral premotor cortex, an increasing gradient in the strength of functional connectivity patterns was found when moving from dorsal to ventral parts of areas F2 and F7.

As previously described for the human brain (for a recent comprehensive review see Palomero-Gallagher and Zilles, 2018), not all receptors reveal all cytoarchitectonic borders, but when the border of a given area within the macaque agranular frontal cortex was highlighted by changes in the regional and/or laminar distribution pattern of a specific receptor type, we found its position to be concordant with that demonstrated by other receptors, as well as by changes in cytoarchitecture. Analysis of receptor architecture provides crucial improvements to that of the cytoarchitecture. First, the possibility of precise quantification due a high specificity of the radioactively labeled ligand binding to individual receptor types, and second, the analysis of multiple receptor types in neighboring sections through entire human or monkey hemispheres, which allows a multimodal and statistically testable approach to validate the distinction between areas or subareas defined during cytoarchitectonic mapping (Schleicher and Zilles, 1988; Zilles et al., 2002b; Palomero-Gallagher and Zilles, 2018). Furthermore, neurotransmitters and their receptors constitute key molecules of signal transmission, and receptor fingerprints provide information on the hierarchical aspect of cortical functional organization (Zilles et al., 2002a; Palomero-Gallagher and Zilles, 2018). Finally, each cytoarchitectonic area is characterized by a unique pattern of local inputs and outputs, i.e. a ‘connectional fingerprint’, that underlies an overall regional connectivity and also subserves its function (Passingham et al., 2002). In order to understand the specific role of a cortical area in complex cognitive functions, it is necessary to integrate insights obtained from structural analyses (cyto- and receptor architecture as well as connectivity patterns) and functional imaging studies. Therefore, focus of the present discussion is twofold: i) the comparison of our parcellation scheme with existing maps of macaque motor and premotor cortex, and ii) the implication of the results of our multimodal statistical analysis, where we emphasize not just structural, but also functional aspects of receptor architecture. Branching of hierarchical clusters has been examined in regard to the framework of previously published connectivity and electrophysiological studies, pertaining the agranular frontal cortex.

### 4.1 Comparison with previous subdivisions of macaque motor and premotor cortex

The macaque primary motor cortex occupies the cortex along the dorsal wall of the central sulcus, as well as the rostrally adjacent precentral convexity, and extends onto the medial surface of the hemisphere. It is characterized by the lack of a visible layer IV and the presence of prominent giant pyramids (Betz cells; Betz, 1874) in layer Vb. The primary motor cortex is functionally heterogeneous, and representations of movements in specific anatomical divisions of the body have been mapped in a somatotopic-like medio-lateral cortical sequence in the human (Woolsey et al., 1952) and non-human primate (Gould et al., 1986; Strick and Preston, 1982a,b; Stepniewska et al., 1993) brain. However, the ensuing motor homunculus is not considered to be reliable parcellation criteria because regions overlap and have multiple representation locations (Park et al., 2001). To date, most maps of the monkey brain illustrate the primary motor cortex as being cytoarchitectonically homogenous, although it has been assigned different names: area 4 (Brodmann 1905; Barbas and Pandya 1987), M1(4) (Preuss and Goldman-Rakic, 1991), 4(M1) (Morecraft et al 2012), F1 (Matelli et al., 1985), or MI(F1) (Caminiti et al., 2017). Conversely, some authors have proposed that the primary motor cortex may be composed of architectonically distinct parts (Preuss et al., 1997; Rathelot and Strick, 2009; Stepniewska et al., 1993), as is the case for the human (Geyer et al., 1996). The present results of the quantitative multimodal analysis revealed the existence of three cyto- and receptor architectonically distinct subdivisions within the macaque primary motor cortex: area 4m on the medial surface, area 4a occupying the precentral convexity, and area 4p, located mostly within the central sulcus where, in the fundus, it adjoins somatosensory area 3a. Stepniewska et al. (1993) identified a rostral (M1r) and a caudal (M1c) subdivision of the owl monkey primary motor cortex based on structural and functional differences, whereby pyramids of M1c are larger than those of M1r. Preuss et al. (1997) identified caudal (area 4c), intermediate (area 4i), and rostral (area 4r) subdivisions within the lateral portion of macaque area 4 based on differences in the size and packing density of layer III and layer V SMI-32-immunoreactive pyramids, whereby they found most obvious differences between the sulcal cortex (area 4c) and that of the precentral convexity (areas 4i and 4r). Area 4c would be equivalent of our area 4p, whereas our area 4a encompasses areas 4i and 4r. The differential distribution of a specific population of corticospinal neurons innervating forelimb muscles, the cortico-motoneuronal cells, enabled the definition of two distinct subregions within the primary motor cortex, i.e. the ‘old’ M1 and the ‘new’ M1 (Rathelot and Strick, 2009). Most cortico-motoneuronal cells were found within the central sulcus (correspond to the ‘new’ M1), at a location comparable to that of our area 4p, whereas the surface of the precentral gyrus, where we identified area 4a, only presented a few scattered cortico-motoneuronal cells (the so-called ‘old’ M1). Finally, and although it was not described by the authors (Matelli et al., 1985), differences in the intensity of staining for cytochrome oxidase also enable definition of areas 4a and 4p in the primary motor cortex of *Macaca nemestrina*. Within the area identified as F1, staining intensity for cytochrome oxidase is clearly weaker in the cortex of the rostral wall of the central sulcus than in that of the adjoining precentral convexity (see Fig. 1, section 34 in Matelli et al., 1985). Previous architectonic studies have reported consistent medio-lateral variations in the size of layer V pyramids, with the largest ones found within the medial portion of the primary motor cortex (Stepniewska et al., 1993; Wiesendanger, 1981), and our present results confirm these observations. However, since cytoarchitectonic differences were accompanied by differences in laminar distribution patterns of multiple receptors, we defined the mesial portion of the primary motor cortex as a distinct region, i.e. 4m.

Although initially described as a single area (i.e. area 6; Brodmann 1905), the premotor cortex is now known to be a complex mosaic composed of structurally and functionally distinct areas responsible for processing different aspects of motor behavior (Rizzolatti et al., 1987; Wise et al., 1985; Barbas and Pandya, 1987; Preuss and Goldman-Rakic, 1991; Matelli et al., 1985, 1991, 1998; Dum and Strick, 2002; Geyer et al., 2000). The medial portion of area 6 has been subdivided into caudal (area F3 or SMA of Matelli et al., 1985, 1991 and Caminiti et al., 2017; area 6m of Morecraft et al., 2012) and rostral areas (area F6 of Matelli et al., 1985, 1991 and Caminiti et al., 2017; pre-SMA of Morecraft et al., 2012). Similar to the primary motor cortex, electrical stimulation of SMA in monkey revealed an additional complete somatotopical map of the body motor representation (Woolsey et al., 1952), whereas movements in area F6 are mostly related to the arm control and orientation (Mitz and Wise, 1987; Luppino et al., 1991). The existence of areas F3 and F6 without additional subdivisions was confirmed by histochemical (Matelli et al., 1985; Geyer et al., 1998), cytoarchitectonic (Matelli et al., 1991; Geyer et al., 1998), connectivity (Luppino et al., 1993), and electrophysiological data (Rizzolatti et al., 1996), and is further supported by the results of the present study. Expanding on a previous receptor architectonic study (Geyer et al., 1998), we were able to show that areas F3 and F6 also differ in their GABA_A_/BZ concentration levels, which were lower in F6 than F3.

Medial premotor areas extend a little over the hemispheric midline and encroach onto the dorsal premotor convexity, where they are delimited by dorsal premotor area 6D of Preuss and Goldman-Rakic (1991), which encompasses a caudal (F2 of Matelli et al., 1985, 1991; or 6DC of Petrides and Pandya, 2006; Morecraft et al., 2012) and a rostral premotor region (F7 of Matelli et al., 1985, 1991; or 6DR of Petrides and Pandya, 2006; Morecraft et al., 2012). We identified dorsal and ventral subdivisions of F2, F2d and F2v respectively, with regard to the *spcd*, and our parcellation is in accordance with the results of previously published immunohistochemical (Geyer et al., 2000), connectivity (Caminiti et al., 2017), cytoarchitectonic and functional (Matelli et al., 1998) analyses. Within rostral area F7 (Matelli et al., 1985, 1991), we identified three areas, i.e. dorsal F7d, intermediate F7i and sulcal F7s, based on cytoarchitectonic differences mainly in layer VI. Area F7d partly corresponds in position to that of the rostro-dorsal oculomotor area SEF (Schlag and Schlag-Rey, 1987), but extends more caudally than does SEF. In addition, Preuss and Goldman-Rakic (1991) referred to the cortex on the dorsal wall of the *sas* as area 6Ds, which largely corresponds to our area F7s.

Parcellation schemes of the ventral convexity, which is occupied by ventral premotor area 6V of Petrides and Pandya (2006), differ considerably from each other, since in some cases it was subdivided dorso-ventrally (dorsal 6Va and ventral part 6Vb; Preuss and Goldman-Rakic, 1991; Morecraft et al., 2012; dorsal 6DC, intermediate 4C and ventral 6V; Barbas and Pandya, 1987) and in others rostro-caudally (rostral F4 and caudal F5; Matelli et al., 1985, 1991). Our results reconcile, at least in part, these diverging parcellation schemes, since we could not only identify areas F4 and F5, but also dorso-ventral distinctions within them in both cyto- and receptor architecture. Thus, area F4 of Matelli et al. (1985, 1991) would encompass our areas F4s, occupying the ventral wall of the arcuate sulcus spur caudal to F5, and two more areas on the free surface of the hemisphere, i.e. F4d dorsally and F4v ventrally. Preuss and Goldman-Rakic (1991) reported a differentiation of their area 6V, with a lighter myelination of the cortex within the arcuate sulcus (occupied by our area F4s) than the adjoining cortex on the convexity (encompassing our areas F4d and F4v). Furthermore, functionally identified areas F4d and F4v (Maranesi et al., 2012) are comparable in extent and location to our areas F4d and F4v, respectively. Finally, our areas F4s and F4d coincide in location and architecture with area 4C of Barbas and Pandya (Barbas and Pandya, 1987).

Within area F5 (Matelli et al., 1985, 1991) we identified three subdivisions based on differences in cyto- and receptor architecture: area F5s occupies the outer portion of the ventral wall of the inferior arcuate branch and is delimited ventro-laterally by F5d, which in turn is located dorsal to area F5v. Belmalih et al. (2009) also identified three areas within F5 based on cyto-, myelo- and chemoarchitectonic observations. However, they describe two areas along the ventral wall of the inferior arcuate branch, i.e. posterior F5p and anterior F5a, and a single area F5c on the lateral surface below the inferior arcuate branch (Belmalih et al., 2009). Area F5a would be the equivalent of our area F5s, area F5c encompasses our areas F5d and F5v, whereas the location of F5p corresponds to that of our area F4s. We found the cyto- and receptor architecture of area F4s to be more similar to that of areas F4d and F4v, than to that of our areas F5s, F5d and F5v. Similarly, Maranesi et al. (2012) also consider that F5p (Belmalih et al., 2009) is part of F4, based on the analysis of intracortical microstimulation and extracellular recordings. Furthermore, the fact that hand movements were only represented in the most dorsal part of area F5 (Maranesi et al., 2012) further supports our subdivision of the postarcuate ventral convexity into dorsal F5d and ventral F5v. Areas F5d and F5v correspond by location, size and cytoarchitecture to areas F5c and DO of Belmalih et al., 2009, respectively.

### 4.2 Correlation between structural, neurochemical and functional organization

Studies in both humans and monkeys, revealed a rostro - caudal, as well as dorso - ventral distinction of the functional organization within the premotor cortex. (Passingham, 1993; Rizzolatti et al., 1998; Geyer et al., 2000). Posterior areas, neighboring the primary motor cortex, are active during more simple movements, e.g. when task is routine, whereas anterior areas are involved in the control of more complex movements, e.g. when additional or new motor/cognitive input is introduced (Passingham, 1993; Rizzolatti et al., 1998; Geyer et al., 2000). As noted in Fig. 11, the distribution of α_1_ receptors follows a caudo-rostral gradient, since caudal areas (F3, F2 and F4) have higher concentration levels than rostral ones (F6, F7 and F5). In contrast, we observed a rostro-caudal gradient (i.e. the opposing trend) for kainate, α_2_, 5HT_1A_ and 5HT_2_ receptors, i.e. higher concentrations have been recorded in the rostral premotor areas than caudal ones (Supplementary Fig. 7).

Functional differences can also be recognized in a dorso-ventral direction, where dorsal (and medial) areas are active when movement is guided by internal inputs, referring to the internal feedback loops (e.g. basal ganglia) and/or proprioception. On the other hand, activation of the ventral areas is guided by the exteroceptive inputs, e.g. visual or auditory stimuli (Passingham, 1993; Rizzolatti et al., 1998; Geyer et al., 2000). A similar trend can be identified for GABA_A_ and M_1_ receptors, as their concentration levels showed clear dorso-ventral differences. For GABA_A_ lower concentration levels were recorded in medio-dorsal areas (F3, F6, F2 and F7), whereas, concentration levels for M_1_ were reversed, with lower values measured in ventral areas (F5 and F4v). Most receptors, i.e. AMPA, NMDA, GABA_B_, and M_3_ as well as GABA_A_/BZ binding sites (Supplementary Fig. 7), show lower receptor densities in dorsal subdivisions of areas F2 and F7 than in ventral subdivisions of areas F4 and F5, as presented for GABA_A_ in Fig. 11. The opposite trend holds true only for M_1_ and M_2_ receptors (Fig. 11; Supplementary Fig. 7).

Furthermore, areas on the medial surface (4m, F3 and F6) show a clear rostro - caudal gradual trend for most receptors, i.e. AMPA, kainate, GABA_A_, GABA_B_, M_1_ and 5HT_2_, where receptor levels were highest in rostral area F6 and lowest in caudal area 4m. The opposite trend is observed only in GABA_A_/BZ binding sites (Fig. 11; Supplementary Fig. 7). Finally, premotor areas F3 and F6 show no significant differences in M_3_ and 5HT_1A_ receptor concentration levels (Supplementary Fig. 7), but rostro - caudal differences are obvious when compared to motor area 4m.

We will here discuss the differences in cyto- and receptor architecture as well as in functional connectivity patterns in the framework of the clusters revealed by the multivariate analyses of functional connectivity fingerprints, and whether significant differences in functional connectivity patterns are associated with significant differences in receptor architecture.

#### 4.2.1 Primary motor cortex

The uniqueness of area 4 with regard to pre-motor areas is reflected in its molecular composition, since the receptor fingerprints of the subdivisions of area 4 (see Supplementary Fig. 8) had a clearly distinct shape and were the smallest of all examined areas. In particular, as observed in humans (Zilles and Palomero-Gallagher, 2017), in macaques the densities of almost all examined receptor types were lower in the primary motor areas than in any premotor area, resulting in an early segregation in the hierarchic cluster analysis.

We found the receptor and functional connectivity fingerprints of area 4p to differ conspicuously from those of areas 4a and 4m. Across all motor and premotor areas, area 4p has the strongest functional connectivity to the somatosensory cortex, in particular to area 3, and to parietal area 7B, related to the somatomotor responses (Andersen et al., 1990a). Conversely, areas 4a and 4m clustered with the dorsolateral premotor areas as those areas revealed to have similar connectivity patterns. They present a stronger functional connectivity with areas 7A and 7m, which have been associated with the control of visuomotor coordination (Andersen et al., 1990a,b; Leichnetz, 2001), than does area 4p (Fig. 13).

These findings correspond with the two subdivisions defined within the monkey primary motor cortex, the so-called ‘old’ and ‘new’ M1, based on ontogenetic and evolutionary aspects as well as on differences in the packing density of cortico-motoneuronal neurons (Rathelot and Strick, 2009). Based on their topographic location, ‘new’ M1 would correspond to our area 4p and ‘old’ M1 to our area 4a. Cortico-motorneuronal cells are mainly found within ‘new’ M1 (Rathelot and Strick, 2009), thus enabling control of the finest movements, such as independent finger movements (Porter and Lemon, 1993), as well as programing novel patterns of motor output in order to acquire a new skill (Rathelot and Strick, 2009). A rostro-caudal structural and functional segregation has also been described within the human primary motor cortex (Geyer et al., 1996; Binkofski et al., 2002). Human 4a is responsible for maintaining the execution of the motor plan, independently of the motor attention, whereas activity of area 4p is modulated by motor-directed attention. These results indicate that area 4p, though not 4a, plays a role in the acquisition of new and/or more precise motor skills that require a higher degree of attention. The lateral portions of human areas 4a and 4p would be equivalent of our areas 4a and 4p.

Multivariate analyses of receptor fingerprints resulted in a clustering of lateral premotor areas F4s and F4d with the subdivisions of the primary motor cortex (Fig. 12). Indeed, out of all premotor areas, F4s and F4d had the lowest receptor concentration levels for most of studied receptor types, and thus display a higher similarity with the primary motor areas. Interestingly, the region occupied by our areas F4s and F4d coincides with area 4C of Barbas and Pandya (Barbas and Pandya, 1987), which they consider to be a subregion of the primary motor cortex, and not of premotor area 6. However, although the receptor architecture of F4s resembled that of F4d, these two areas differed significantly in their functional connectivity.

Primary motor cortex plays a crucial role in voluntary movements. Thus, in order to control movement of limbs and other body parts, area 4 is thought to integrate information from area 5 on the spatial location of the body parts (Lacquaniti et al., 1995). Our functional connectivity analysis provides further support for this hypothesis, as all three subdivisions of the primary motor cortex have a strong functional connectivity with area 5 (Fig. 13), a higher-order somatosensory area involved in the analysis of proprioceptive information (Bakola et al., 2013), where most neurons encode the location of the arm in space relative to the body posture (Lacquaniti et al., 1995). This may underlie the functional synchronization required within the areas of cluster 4 to plan voluntary limb movements based on visual, auditory and/or somatosensory guidance, when the animal moves toward the object to accomplish the reaching distance.

#### 4.2.2 Coordination of hand-to-mouth movements

Both ventral premotor areas, F4 (Gentilucci et al., 1988) and F5 (Rizzolatti et al., 1988), have similar motor representations of the hand and of the mouth. However, F4 neurons are associated with proximal hand movement, the activation of the hand region in area F5 is related to distal hand orientation and movement (Gentilucci et al., 1988), guiding the goal-directed hand tasks for reaching or grasping food and bring it to the mouth (Rizzolatti et al., 1988). Within F5, hand movement was evoked on the bank of *ias* and on the dorsal portion of the lateral convexity, where it overlaps with the ventrally located mouth activation (Kurata and Tanji, 1986; Gentilucci et al., 1988; Rizzolatti et al., 1988; di Pellegrino et al., 1992, Ferrari et al., 2003; Maranesi et al., 2012), which seems to extend over the fronto-opercular region (Ferrari et al., 2003; Maranesi et al., 2012). This is clearly comparable with our three subdivisions of the rostral ventral premotor cortex, with area F5s within the *ias*, followed by F5d on the dorsal portion of the lateral convexity and F5v (comparable to area DO of Ferrari et al., 2017) on its ventral portion.

Area F4d as identified in the present study is comparable in location and extent to the functionally defined area F4d of Maranesi et al. (2012), which encodes hand and face movements (Maranesi et al., 2012). While the ventral portion of area F4, i.e. where we identified our area F4v, has been associated with mouth movements, in particular the control of tongue and simple oro-facial movements (Maranesi et al., 2012). Additionally, compared to dorsal parts of F4, the ventral sector showed distinct sensory properties. The majority of projections were revealed to be nonvisual, i.e. somatosensory and proprioceptive responses were widely represented, receiving sensory information from the oro-facial body parts (Maranesi et al., 2012). This is in correspondence with the result of our functional connectivity analysis as F4v shows stronger connection to the somatosensory cortex then F4d. Furthermore, caudal (F4) and rostral (F5) potions of the ventral premotor cortex share strong reciprocal connections (Matelli et al., 1998), and receive most prominent projections from areas involved in somatosensory and somatomotor responses, such as area 7B (Andersen et al., 1990a) and primary somatosensory cortex (Fig. 13). However, areas F5, more notably F5v, are strongly connected to area ProM, which has previously also been associated with the gustatory, orbitofrontal, insular and somatosensory cortex and plays role in the feeding process (Cipolloni and Pandya, 1999), whereas subdivisions of F4, in particular F4v, have a strong functional connectivity with primary somatosensory areas 2 and 3. Additionally, mirror neurons have been identified within area F5 mainly on the postarcuate convexity, although some have also been found within *ias* (Rizzolatti and Craighero, 2004; Rizzolatti and Fogassi, 2014). Mirror neurons fire not only during movement execution, but also during the observation of an object-directed action being performed by another individual (Rizzolatti et al., 1996, Gallese et al., 1996). Although most of neurophysiological studies focused on mirror neurons in relation to hand actions, some also analyzed mouth mirror neurons, which display similar visuomotor properties to those of hand mirror neurons (Ferrari et al., 2003). Interestingly, only a small population of mouth mirror neurons responds to communicative gestures, such as lipsmacking, and are not restricted to the ventral portion of F5, but are also found within F4v (Ferrari et al., 2003). Thus, both areas, F5v and F4v, could play complementary roles, at different hierarchical levels, in the control of monkey vocalization, although their main role is thought to be related to food-processing behavior (Hoshi and Tanji, 2004).

#### 4.2.3 Postarcuate region codes peripersonal space

Neuronal activity recorded within and around the spur of the arcuate sulcus showed that this region is visually responsive and activated during saccades (Baker et al., 2006; Koyama et al., 2004) and it has been denominated as premotor eye field (Amiez and Petrides, 2009). In particular, the dorsal portion of the arcuate sulcus, corresponding to our area F2v, contains a forelimb representation, as well as different types of visually responsive neurons responsible of coding a peripersonal space, similar to areas F4s and F4d, only they hold face, hand and mouth representation (Fogassi et al., 1999). Hence it has been suggested that areas in the postarcuate region, constitute a somatocentered map, used for visual navigation and control of different actions (Fogassi et al., 1999).

Area F4s is associated with areas LIP, 7m and 7A, all related to the saccadic eye movement and visuospatial perception (Andersen et al., 1990a,b; Leichnetz, 2001). Multivariate analysis of functional connectivity fingerprints revealed it to cluster with rostral areas F7s (located in the *sas*) and F5s (extending along the *ias*). The ventral part of F7 (which encompasses our areas F7i and F7s) is reported to use information from medial parietal area PGm (or 7m) to locate the object in space for orientation, as well as to coordinate arm-body movements (Matelli et al., 1998; Luppino and Rizzolatti, 2000). Area F5s contains neurons with unique responses to visual stimuli, so-called ‘canonical’ neurons (Rizzolatti et al., 1998; Fogassi et al., 2001, Kakei et al., 2001), which are active when monkey observes a visual object and executes a hand-based action (Rizzolatti et al., 1996, Gallese et al., 1996).

It is worth noting, that within cluster 3 we find area F2v closely related to area F3, i.e. the supplementary motor area (SMA). The electrical stimulation of this area revealed a complete somatotopical map of the body representation, in addition to the one in the primary motor cortex (Woolsey at al., 1952). F3 is active when motor task demands certain conditions or retrieval of the motor memory (Tanji, 1994). Additionally, it plays important role in organizing movements, especially when action requires performing a set of serial movements (Tanji, 1994).

F3 has strong reciprocal connections with rostrally neighboring area F6 (Luppino and Rizzolatti, 2000). Although these two areas show a great similarity of their receptor fingerprints, they differ considerably in their functional connectivity patterns. F3 is the source of dense, topographically organized corticospinal projections and strong cortico-cortical connections to area 4 and other premotor areas (F2, F4 and F5; Luppino and Rizzolatti, 2000), whereas F6 does not control movement directly, but serves as the major input of limbic and prefrontal information to all caudal and rostral premotor areas (Luppino and Rizzolatti, 2000). This is in accordance with our functional connectivity findings, where we can see segregation of these areas into different cluster groups. Indeed, whereas F3 is found in cluster C3 with caudal lateral premotor area F2v, area F6 is in cluster C4 with rostral lateral premotor area F7i (Fig. 14).

#### 4.2.4 Integration of different sensory inputs

Neurons in the dorsal premotor cortex are involved in integrating information about which arm to use or which target to reach (Hoshi and Tanji, 2004). Thus, it is no surprise that area F6 correlated with these areas. as activations in area F6 (or pre-SMA) are mostly related to arm movements (Mitz and Wise, 1987; Luppino et al., 1991) and target localization (Hoshi and Tanji, 2000, 2004). However, it is interesting that area F7i showed highest similarity with the medial area F6, although neurochemically resembled area F7d more closely.

The cluster analysis based on the receptor fingerprints showed a clear segregation of the subdivisions within areas F2 and F7, since areas dorsal to the *spcd* (i.e. F2d and F7d) were revealed to be more similar from the neurochemical point of view to medial areas F3 and F6, than to their corresponding ventral subdivisions (i.e. F2v, F7i and F7s) (Fig. 12). Thus, we here provide further results demonstrating that areas F2 and F7 each consist of at least two functionally distinct sectors, as suggested by Rizzolatti et al. (1998). Area F2d, previously described as dimple area F2dc (Matelli et al., 1998), receives projections from areas PEip and PEc, two higher-order areas involved in the amplification of somatosensory stimuli, in order to plan and coordinate, mostly, leg movements (Matelli et al., 1998). The dorsal portion of rostral premotor area F7 (F7d) has been associated with oculomotor and visuospatial functions, and receives main inputs from the inferior parietal cortex, e.g. area PG, which encodes eye orientation (Sakata et al., 1980), and from the intraparietal cortex, e.g. area LIP, which encodes eye movement (Andersen et al., 1990b; Snyder et al., 1997; Huerta and Kaas, 1990).

### 4.3 Integration of in-vivo functional data with post-mortem anatomy

Functional imaging of the macaque has great potential to aid in the translation between cutting edge anatomy and physiology that is available in the macaque brain, and the in-vivo imaging descriptions of the functional anatomy of the human brain in health and disease. Several groups have used invasive tract-tracing data to test and validate diffusion MRI tractography methods (Dauget et al., 2007; Jbabdi et al., 2013; van den Heuvel et al., 2015). However, integration of high-quality anatomical data with macaque functional imaging has been slow, with few exceptions (Scholtens et al., 2015; Froudist-Walsh et al., 2018). In part this is due to post-mortem anatomical and in-vivo imaging data not being reported in a common stereotaxic space. Here we demonstrate how by combining cytoarchitecture, receptor and functional imaging data we can enrich our understanding of cortical anatomy. Furthermore, we openly share our parcellation and receptor data in a standard neuroimaging space order to make it easier for cytoarchitecture and receptor anatomy to inform future imaging studies.

## 5 CONCLUSIONS

We here present a 3D atlas of macaque motor and premotor areas based on a quantifiable and statistically testable analysis of its cyto- and receptor architecture. Multivariate analyses of the receptor fingerprints revealed the existence of a caudal cluster (encompassing primary motor areas and ventral premotor areas F4s and F4d), a dorsal cluster (encompassing areas F3 and F6 on the medial surface of the hemisphere as well as areas F2d, F2v, F7d, F7i and F7s on its dorsolateral surface), and a ventral cluster (encompassing area F4v and all subdivisions of area F5). Motor and premotor areas are involved in the integration of sensory information in order to plan and execute task-related movements. Interestingly, our functional connectivity analysis revealed that areas of the dorsal cluster show a stronger functional connectivity with areas involved in spatial processing than do areas of the ventral cluster. Conversely, areas of the ventral cluster show a stronger functional connectivity with areas involved in the processing of somatosensory input than do areas of the ventral cluster.

The proposed parcellation scheme, which integrates and reconciles the discrepancies between previously published maps of this region, was projected onto the Yerkes19 surface (Donahue et al., 2016) together with the receptor fingerprints of each identified area. Thus, we provide the neuroscientific community for the first time with a 3D map of the macaque agranular frontal cortex integrating information on its cyto- and receptor architecture and demonstrate how it constitutes a valuable resource for the analysis of functional experiments carried out with non-human primates. Furthermore, the receptor fingerprints provide valuable data for modelling approaches aiming to a better understanding of the complex structure of the neural system, as well as to provide an insight in the evolution of the healthy primate brain.

## ACKNOWLEDGEMENTS

This project has received funding from the European Union’s Horizon 2020 Framework Programme for Research and Innovation under the Specific Grant Agreements 785907 (Human Brain Project SGA2) and 945539 (Human Brain Project SGA3), as well as from the National Institute of Health (NIH) under grant number R01MH122024-02 and the Federal Ministry of Education and Research (BMBF) under project number 01GQ1902.

## AUTHOR CONTRIBUTIONS

**Lucija Rapan:** Investigation, Validation, Visualization, Writing – Original Draft, **Sean Froudist-Walsh:** Conceptualization, Software, Writing – Review and Editing, Funding acquisition, **Meiqi Niu:** Data Curation, Visualization, Writing – Review and Editing, **Ting Xu:** Resources, Data Curation, Writing – Review and Editing, **Thomas Funck:** Formal Analysis, **Karl Zilles:** Conceptualization, Resources, Writing – Review and Editing, Supervision, Funding acquisition, **Nicola Palomero-Gallagher:** Conceptualization, Resources, Data Curation, Supervision, Writing – Review and Editing, Project administration, Funding acquisition.

## REFERENCES

Amiez, C., & Petrides, M. (2009). Anatomical organization of the eye fields in the human and non-human primate. Progress in Neurobiology, 89:220–230.

Andersen, R. A., Asanuma, C., Essick, G., & Siegel, R. M. (1990a). Corticocortical connections of anatomically and physiologically defined subdivisions within the inferior parietal lobule. The Journal of Comparative Neurology, 296:65–113.

Andersen, R. A., Bracewell, R. M., Barash, S., Gnadt, J. W., & Fogassi, L. (1990b). Eye position effects on visual, memory, and saccade-related activity in areas LIP and 7a of macaque. The Journal of Neuroscience, 10:1176–96.

Autio, J. A., Glasser, M. F., Ose, T., Donahue, C. J., Bastiani, M., Ohno, M., … Hayashi, T. (2020). Towards HCP-style macaque connectomes: 24-channel 3T multi-array coil, MRI sequences and preprocessing. NeuroImage, 215:11580.

Baker, J., Patel, G. H., Corbetta, M., & Snyder, L. H. (2006). Distribution of activity across the monkey cerebral cortical surface, thalamus and midbrain during rapid, visually guided saccades. Cerebral Cortex, 16:447–459.

Bakola, S., Passarelli, L., Gamberini, M., P., F., & Galletti, C. (2013). Cortical connectivity suggests a role in limb coordination for macaque area PE of the superior parietal cortex. The Journal of Neuroscience, 33:6648–6658.

Barbas, H., & Pandya, D. N. (1987). Architecture and frontal cortical connections of the premotor cortex (area 6) in the thesus monkey. The Journal of Comparative Neurology, 256:211–228.

Belmalih, A., Borra, E., Contini, M., Gerbella, M., Rozzi, S., & Luppino, G. (2009). Multimodal architectonic subdivision of the rostral part (area F5) of the macaque ventral premotor cortex. Research in Systems Neuroscience, 512:183–217.

Benjamini, Y., & Hochberg, Y. (1995). Controlling the false discovery rate: A practical and powerful approach to multiple testing. Journal of the Royal Statistical Society: Series B (Methodological), 57:289–300.

Betz, W. (1874). Anatomischer Nachweis zweier Gehirncentra. Centralblatt für die medizinische Wissenschaften, 12:578–580, 595-599.

Binkofski, F., Fink, G. R., Geyer, S., Buccino, G., Gruber, O., Shah, N. J., … Freund, H. J. (2002). Neural activity in human primary motor cortex areas 4a and 4p is modulated differentially by attention to action. Journal of Neurophysiology, 88:514–519.

Boussaoud, D., Barth, T. M., & Wise, S. P. (1993). Effects of gaze on apparent visual responses of frontal cortex neurons. Experimental Brain Research, 93:423–434.

Brodmann, K. (1905). Beiträge zur histologischen Lokalization der Grosshirnrinde. III. Mitteilung: Die Rindenfelder der niederen Affen. Journal of Neurology and Psychology, 4:177–226.

Brodmann, K. (1909). Vergleichende Lokalisationslehre der Grosshirnrinde in ihren Prinzipien dargestellt auf Grund des Zellenbaues. Leipzig: Barth JA.

Caminiti, R., Borra, E., Visco-Comandini, F., Battaglia-Mayer, A., Averbeck, B., & Luppino, G. (2017). Computational architecture of the parieto-frontal network underlying cognitive-motor control in monkeys. eNEURO, 4:e0306–16.

Caminiti, R., Ferraina, S., & Johnson, P. B. (1996). The sources of visual information to the primate frontal lobe: a novel role for the superior parietal lobule. Cerebral Cortex, 6:319–328.

Caspers, J., Palomero-Gallagher, N., Caspers, S., Schleicher, A., Amunts, K., & Zilles, K. (2015). Receptor architecture of visual areas in the face and word-form recognition region of the posterior fusiform gyrus. Brain Structure and Function, 220:205–219.

Caspers, S., Schleicher, A., Bacha-Trams, M., Palomero-Gallagher, N. A., & Zilles, K. (2012). Organization of the human inferior parietal lobule based on receptor architectonics. Cerebral Cortex, 23:615–628.

Cipolloni, P., & Pandya, D. N. (1999). Cortical connections of the frontoparietal opercular areas in the rhesus monkey. The Journal of Comparative Science, 403:431–458.

Colby, C. L., Duhamel, J. R., & Goldberg, M. E. (1993). Ventral intraparietal area of the macaque: anatomic location and visual response properties. Journal of Neurophysiology, 69:902–914.

Cooke, D. F., & Graziano, M. S. (2003). Defensive movements evoked by air puff in monkeys. Journal of Neurophysiology, 90:3317–3328.

Dauguet, J., Peled, S., Berezovskii, V., Delzescaux, T., Warfield, S. K., Born, R., & Westin, C.-F. (2007). Comparison of fiber tracts derived from in-vivo DTI tractography with 3D histological neural tract tracer reconstruction on a macaque brain. Neuroimage, 37:530–538.

di Pellegrino, G., & Wise, S. P. (1993). Visuospatial versus visuomotor activity in the premotor and prefrontal cortex of a primate. The Journal of Neuroscience, 13:1227–1243.

di Pellegrino, G., Fadiga, L., Fogassi, L., Gallese, V., & Rizzolatti, G. (1992). Understanding motor events: a neurophysiological study. Experimental Brain Research, 91:176–180.

Donahue, C. J., Sotiropoulos, S. N. S. J., Hernandez-Fernandez, M., Behrens, T. E., Dyrby, T. B., … Glasser, M. F. (2016). Using diffusion tractography to predict cortical connection strength and distance: A quantitative comparison with tracers in the monkey. The Journal of Neuroscience, 36:6758–70.

Dum, R. P., & Strick, P. L. (2002). Motor areas in the frontal lobe of the primate. Physiology & Behavior, 77:677–682.

Ferrari, P. F., Gallese, V., Rizzolatti, G., & Fogassi, L. (2003). Mirror neurons responding to the observation of ingestive and communicative mouth actions in the monkey ventral premotor cortex. European Journal of Neuroscience, 17:1703–1714.

Ferrari, P., Gerbella, M., Coudé, G., & Rozzi, S. (2017). Two different mirror neuron networks: the sensorimotor (hand) and limbic (face) pathways. Neuroscience, 358:300–315.

Fogassi, L., Gallese, V., Buccino, G., Craighero, L., Fadiga, L., & Rizzolatti, G. (2001). Cortical mechanism for the visual guidance of hand grasping movements in the monkey: A reversible inactivation study. Brain, 124:571–586.

Fogassi, L., Gallese, V., Fadiga, L., Luppino, G., Matelli, M., & Rizzolatti, G. (1996). Coding of peripersonal space in inferior premotor cortex (area F4). Journal of Neurophysiology, 76:141–157.

Fogassi, L., Raos, V., Franchi, G., Gallese, V., Luppino, G., & Matelli, M. (1999). Visual responses in the dorsal premotor area F2 of the macaque monkey. Experimental Brain Research, 128:194–199.

Froudist-Walsh, S., Browning, P. G., Young, J. J., Murphy, K. L., Mars, R. B., Fleysher, L., & Croxson, P. L. (2018). Macro-connectomics and microstructure predict dynamic plasticity patterns in the non-human primate brain. Elife, 7:e34354.

Fujii, N., Mushiake, H., & Tanji, J. (1998). An oculomotor representation area within the ventral premotor cortex. Proceedings of the National Academy of Sciences of the United States of America, 95:12034–12037.

Fujii, N., Mushiake, H., & Tanji, J. (2000). Rostrocaudal distinction of the dorsal premotor area based on oculomotor involvement. Journal of Neurophysiology, 83:1764–1769.

Gallese, V., Fadiga, L., Fogassi, L., & Rizzolatti, G. (1996). Action recognition in the premotor cortex. Brain, 119:593–609.

Gentilucci, M., Fogassi, L., Luppino, G., Matelli, M., Camarda, R., & Rizzolatti, G. (1988). Functional organization of inferior area 6 in the macaque monkey I. Somatotopy and the control of proximal movements. Experimental Brain Research, 71:475–490.

Geyer, S., Ledberg, A., Schleicher, A., Kinomura, S., Schormann, T., Bürgel, U., … Roland, P. E. (1996). Two different areas within the primary motor cortex of man. Nature, 382:805–807.

Geyer, S., Matelli, M., Luppino, G., & Zilles, K. (1998). A new microstructural map of the macaque monkey lateral premotor cortex based on neurofilament protein distribution. European Journal of Neuroscience, 10:83–83.

Geyer, S., Matelli, M., Luppino, G., & Zilles, K. (2000). Functional neuroanatomy of the primate isocortical motor system. Anatomy and Embriology, 202:443–474.

Ghosh, S., & Gattera, R. (1995). A comparison of the ipsilateral cortical projections to the dorsal and ventral subdivisions of the macaque premotor cortex. Somatosensory and Motor Research, 12:359–78.

Gould, H. J., Cusick, C. G., Pons, T. P., & Kaas, J. H. (1986). The relationship of corpus callosum connections to electrical simulation maps of motor, supplementary motor, and the frontal eye fields in owl monkeys. The Journal of Comparative Neurobiology, 247:297–325.

Graziano, M. S., & Cooke, D. F. (2006). Parieto-frontal interactions, personal space, and defensive behavior. Neuropsychologia, 44:845–859.

Graziano, M. S., Yap, G. S., & Gross, C. G. (1994). Coding of visual space by premotor neurons. Science, 266:1054–1057.

Gregoriou, G. G., & Savaki, H. E. (2003). When vision guides movement: a functional imaging study of the monkey brain. NeuroImage, 19:959–967.

Hoshi, E., & Tanji, J. (2000). Integration of target and body-part information in the premotor cortex when planning action. Nature, 408:466–470.

Hoshi, E., & Tanji, J. (2004). Differential roles of neuronal activity in the supplementary and presupplemetary motor areas: from information retrieval to motor planning and execution. Journal of Neurophysiology, 92:3482–3499.

Huerta, M. F., & Kaas, J. H. (1990). Supplementary eye field as defined by intracortical microstimulation: connections in macaques. The Journal of Comparative Neurology, 293:299–330.

Impieri, D., Zilles, K., Niu, M., Rapan, L., Schubert, N., Galletti, C., & Palomero-Gallagher, N. (2019). Receptor density pattern confirms and enhances the anatomic-functional features of the macaque superior parietal lobule areas. Brain Structure and Function, 224:2733–2756.

Jbabdi, S. L., Haber, S. N., & Behrens, T. E. (2013). Human and monkey ventral prefrontal fibers use the same organizational principles to reach their targets: tracing versus tractography. Journal of Neuroscience, 33:3190–3201.

Kakei, S., Hoffman, D. S., & Strick, P. L. (2001). Direction of action is represented in the ventral premotor cortex. Nature Neuroscience, 4:1020–1025.

Koyama, M., Hasegawa, I., Osada, T., Adachi, Y., Nakahara, K., & Miyashita, Y. (2004). Functional magnetic resonance imaging of macaque monkeys performing visually guided saccade tasks: comparison of cortical eye fields with humans. Neuron, 41:795–807.

Kurata, K., & Tanji, J. (1986). Premotor cortex neurons in macaques: activity before distal and proximal forelimb movements. The Journal of Neuroscience, 6:403–411.

Lacquaniti, F., Guigon, E., Bianchi, L., Ferraina, S., & Caminiti, R. (1995). Representing spatial information for limb movement: role of area 5 in the monkey. Cerebral Cortex, 5:391–409.

Leichnetz, G. R. (2001). Connections of the medial posterior parietal cortex (area 7m) in the monkey. The Anatomical Record, 263:215–236.

Luppino, G. R. (2000). The organization of the frontal motor cortex. News in Physiological Sciences, 15:219–224.

Luppino, G., Matelli, M., Camarda, R. M., & Rizzolatti, G. (1993). Corticocortical connections of area F3 (SMA-proper) and area F6 (pre-SMA) in the macaque monkey. The Journal of Comporative Neurology, 338:114–140.

Luppino, G., Matelli, M., Camarda, R. M., Gallese, V., & Rizzolatti, G. (1991). Multiple representation of body movements in mesial area 6 and the adjacent cingulate cortex: an intracortical microstimulation study in the macaque monkey. The Journal of Comparative Neurology, 311:463–482.

Luppino, G., Murata, A., Govoni, P., & Matelli, M. (1999). Largely segregated parietofrontal connections linking rostral intraparietal cortex (areas AIP and VIP) and the ventral premotor cortex (areas F5 and F4). Experimental Brain Research, 128:181–187.

Mahalanobis, P. C., Majumdar, D. N., & Rao, C. R. (1949). Anthropometric survey of the United Provinces, 1941: a statistical study. Sankhya: The Indian Journal of Statistics, 9:89–324.

Maranesi, M., Roda, F. B., Rozzi, S., & Ferrari, P. F. (2012). Anatomo-functional organization of the ventral primary motor and premotor cortex in the macaque monkey. The European Journal of Neuroscience, 36:3376–3387.

Markov, N. T., Ercsey-Ravasz, M. M., Ribeiro Gomes, A. R., Lamy, C., Magrou, L., Vezoli, J., … Kennedy, H. (2014). A weighted and directed interareal connectivity matrix for macaque cerebral cortex. Cerebral Cortex, 24:17–36.

Mars, R. B., Passingham, P. E., & Jbabdi, S. (2019). Connectivity fingerprints: from areal descriptions to abstract spaces. Trends in Cognitive Sciences, 22:1026–1037.

Matelli, M., Govoni, P., Galletti, C., Kutz, D. F., & Luppino, G. (1998). Superior area 6 afferents from the superior parietal lobule in the macaque monkey. The Journal of Comparative Neurology, 402:327–352.

Matelli, M., Luppino, G., & Rizzolatti, G. (1985). Patterns of cytochrome oxidase activity in the frontal agranular cortex of the macaque monkey. Behavioural Brain Research, 18:125–136.

Matelli, M., Luppino, G., & Rizzolatti, G. (1991). Architecture of superior and mesial area 6 and the adjacent cingulate cortex in the macaque monkey. The Journal of Comparative Neurology, 311:445–462.

Merker, B. (1983). Silver staining of cell bodies by means of physical development. Journal of Neuroscience Methods, 9:235–241.

Milham, M. P., Ai, L., Koo, B., …, Zhou, Y., Margulies, D. S., & Schroeder, C. E. (2018). An open resource for non-human primate imaging. Neuron, 100:61–74.

Milham, M., Petkov, C., …, & Zhou, Y. (2020). Accelerating the evolution of nonhuman primate neuroimaging. Neuron, 105:600–603.

Mitz, A. R., & Wise, S. P. (1987). The somatotopic organization of the supplementary motor area: intracortical microstimulation mapping. The Journal of Neuroscience, 7:1010–1021.

Morecraft, R., Stilwell-Morecraft, K. S., Cipolloni, B., Ge, J., McNeal, D., & Pandya, D. N. (2012). Cytoarchitecture and cortical connections of the anterior cingulate and adjacent somatomotor fields in the rhesus monkey. Brain Research Bulletin, 87:457–487.

Noonan, M. P., Sallet, J., Mars, R. B., Neubert, F. X., O’Reilly, J. X., Andersson, J. L., … Rushworth, M. F. (2014). A neural circuit covarying with social hierarchy in macaques. PLoS Biology, 12:e1001940.

Palomero-Gallagher, N., & Zilles, K. (2018). Cyto- and receptor architectonic mapping of the human brain. In I. Huitinga, & M. J. Webster, Handbook of Clinical Neurology: Brain Banking in Neurologic and Psychiatric Diseases (pp. 150:355–387). Amsterdam: Elsevier.

Palomero-Gallagher, N., Mohlberg, H., Zilles, K., & Vogt, B. A. (2008). Cytology and receptor architecture of human anterior cingulate cortex. The Journal of Comparative Neurology, 508:906–926.

Palomero-Gallagher, N., Zilles, K., Schleicher, A., & Vogt, B. A. (2013). Cyto- and receptor architecture of area 32 in human and macaque brains. The Journal of Comparative Neurology, 521:3272–3286.

Park, M. C., Belhaj-Saif, A., Gordon, M., & Cheney, P. D. (2001). Consistent features in the forelimb representation of primary motor cortex in rhesus macaques. The Journal of Neuroscience, 21:2784–2792.

Passingham, R. E. (1993). The frontal lobes and voluntary action. In R. E. Passingham, The frontal lobes and voluntary action (Oxford Psychology Series). New York: Oxford University Press.

Passingham, R. E., Stephan, K. E., & Kötter, R. (2002). The anatomical basis of functional localization in the cortex. Nature Reviews Neuroscience, 3:606–616.

Petrides, M., & Pandya, D. N. (1994). Comparative architectonic analysis of the human and macaque frontal cortex. In B. a. Grafman, Handbook of Neuropsychology (pp. 17–58). Amsterdam: Elsevier Scinece Publishers.

Petrides, M., & Pandya, D. N. (2006). Efferent association pathways originating in the caudal prefrontal cortex in the macaque monkey. The Journal of Comparative Neurology, 498:227–251.

Porter, R., & Lemon, R. (1993). Corticospinal function and voluntary movement. Oxford: Oxford University Press.

Preuss, T. M., & Goldman-Rakic, P. (1991). Myelo- and cytoarchitecture of the granular frontal cortex and surrounding regions in the strepsirhine primate Galago and the anthropoid primate Macaca. The Journal of Comparative Neurology, 310:429–574.

Preuss, T. M., Stepniewska, I., Jain, N., & Kaas, J. H. (1997). Multiple divisions of macaque precentral motor cortex identified with neurofilament antibody SMI-32. Brain Research, 767:148–153.

R Core Team. (2020). R: A language and environment for statistical computing. Retrieved from R Foundation for Statistical Computing, Vienna, Austria: http://www.r-project.org/index.html

Rathelot, J. A., & Strick, P. L. (2009). Subdivisions of primary motor cortex based on cortico-motoneuronal cells. Proceedings of the National Academy of Science of the United States of America, 106:918–923.

Rizzolatti, G., & Craighero, L. (2004). The mirror-neuron system. Annual Review of Neuroscience, 27:169–192.

Rizzolatti, G., & Fogassi, L. (2014). The mirror mechanism: recent findings and perspectives. Philosophical Transactions of the Royal Society B: Biological Sciences, 369:20130420.

Rizzolatti, G., Camarda, R., Fogassi, L., Gentilucci, M., Luppino, G., & Matelli, M. (1988). Functional organization of inferior area 6 in the macaque monkey II. Area F5 and the control of distal movements. Experimental Brain Research, 71:491–507.

Rizzolatti, G., Fadiga, L., Gallese, V., & Fogassi, L. (1996). Premotor cortex and the recognition of motor actions. Cognitive Brain Research, 3:131–141.

Rizzolatti, G., Gentilucci, M., Fogassi, L., Luppino, G., Matelli, M., & Ponzoni-Maggi, S. (1987). Neurons related to goal-directed motor acts in inferior area 6 of the macaque monkey. Experimental Brain Research, 67:220–224.

Rizzolatti, G., Luppino, G., & Matelli, M. (1998). The organization of the cortical motor system: new concepts. Electroencephalography and Clinical Neurophysiology, 106:283–296.

Rousseeuw, P. J. (1987). Silhouettes: A graphical aid to the interpretation and validation of cluster analysis. Journal of Computational and Applied Mathematics, 20:53–65.

Sakata, H., Shibutani, H., & Kawano, K. (1980). Spatial properties of visual fixation neurons in posterior parietal association cortex of the monkey. Journal of Neurophysiology, 43:1654–1672.

Sanes, J. N., & Donoghue, J. P. (2000). Plasticity and primary motor cortex. Annual Review of Neuroscience, 23:393–415.

Schlag, J., & Schlag-Rey, M. (1987). Evidence for a supplementary eye field. Journal of Neurophysiology, 57:179–200.

Schleicher, A., & Zilles, K. (1988). The use of automated image analysis for quantitative receptor autoradiography. In F. Van Leeuwen, B. Rm, P. Cw, & P. O, Molecular neuroanatomy (pp. 147–157). Amsterdam: Elsevier.

Schleicher, A., & Zilles, K. (1990). A quantittative approach to cytoarchitectonics: analysis of structural inhomogeneities in nervous tissue using an image analyser. Journal of Microscopy, 157:367–381.

Schleicher, A., Amunts, K., Geyer, S., Kowalski, T., Schormann, T., Palomero-Gallagher, N., & Zilles, K. (2000). A stereological approach to human cortical architecture: identification and delineation of cortical areas. Journal of Chemical Neuroanatomy, 20:31–47.

Schleicher, A., Amunts, K., Geyer, S., Morosan, P., & Zilles, K. (1999). Observer-indepentent method for microstructural parcellation of cerebral cortex: a quantitative approach to cytoarchitectonics. NeuroImage, 9:165–177.

Schleicher, A., Morosan, P., Amunts, K., & Zilles, K. (2009). Quantitative architectural analysis: a new approach to cortical mapping. Journal of Autism and Developmental Disorders, 39:1568–81.

Schleicher, A., Palomero-Gallagher, N., Morosan, P., Eickhoff, S. B., Kowalski, T., DeVos, K., … Zilles, K. (2005). Quantitative architectural analysis: a new approach to cortical mapping. Anatomy and Embryology, 210:373–386.

Scholtens, L. H., Schmidt, R., de Reus, M. A., & van den Heuvel, M. P. (2014). Linking macroscale graph analytical organization to microscale neuroarchitectonics in the macaque connectome. Journal of Neuroscience, 34:12192–12205.

Snyder, L. H., Batista, A. P., & Andersen, R. A. (1997). Coding of intention in the posterior parietal cortex. Nature, 386:167–170.

Stepniewska, I., Preuss, T. M., & Kaas, J. H. (1993). Architectonics, somatotopic organization and ipsilateral cortical connections of the primary motor area (M1) of owl monkey. The Journal of Comparative Neurology, 330:238–271.

Strick, P. L., & Preston, J. B. (1982a). Two representations of the hand in area 4 of primate I: motor output organization. Journal of Neurophysiology, 48:139–149.

Strick, P. L., & Preston, J. B. (1982b). Two representations of the hand in area 4 of primate II: somatosensory input organization. Journal of Neurophysiology, 48:150–159.

Tanji, J. (1994). The supplementary motor area in the cerebral cortex. Neuroscience Research, 19:251–268.

van den Heuvel, M. P., de Reus, M. A., Feldman Barrett, L., Scholtens, L. H., Coopmans, F. M., Schmidt, R., … Li, L. (2015). Comparison of diffusion tractography and tract-tracing measures of connectivity strength in rhesus macaque connectome. Human Brain Mapping, 36:3064–3075.

Wiesendanger, M. (1981). The pyramidal tract. In A. L. Towe, & E. S. Luchei, Motor coordination (pp. 401–491). Boston: Springer.

Wiesendanger, M. (2011). Organization of secondary motor areas of cerebral cortex. In V. B. Brooks, pp. Handbook of physiology, the nervous system, motor control (pp. 2:1121–1147). Washington, DC: American Physiological Society.

Wise, S. P. (1985). The primate premotor cortex: past, present and preparatory. Annual Review of Neuroscience, 8:1–19.

Wise, S. P., Moody, S. L., Blomstrom, K. J., & Mitz, A. R. (1998). Changes in motor cortical activity during visuomotor adaptation. Experimental Brain Research, 121:285–299.

Woolsey, C. N., Settlage, P. H., Meyer, D. R., Sencer, W., Pinto Hamuy, T., & Travis, A. M. (1952). Patterns of localization in precentral and “supplementary” motor areas and their relation to the concept of a premotor area. Research Publications-Association for Research in Nervous and Mental Disease, 30:238–264.

Xu, T., Sturgeonghi, D., Ramirezghi, J. S., Froudist-Walshb, S., Marguliesj, D. S., Schroedercde, C. E., … Milhamaf, M. P. (2019). Interindividual variability of functional connectivity in awake and anesthetized rhesus macaque monkeys. Biological Psychiatry: Cognitive Neuroscience and Neuroimaging, 4:543–553.

Zilles, K., & Amunts, K. (2010). Centenary of Brodmann’s map--conception and fate. Nature Reviews: Neuroscience, 11:139–45.

Zilles, K., & Palomero-Gallagher, N. (2017). Multiple transmitter receptors in regions and layers of the human cerebral cortex. Frontiers in Neuroanatomy, 11:78.

Zilles, K., Palomero-Gallagher, N., Grefkes, C., Scheperjans, F., Boy, C., Amunts, K., & Schleicher, A. (2002a). Architectonics of the human cerebral cortex and transmitter receptor fingerprints: reconciling functional neuroanatomy and neurochemistry. European Neuropsychopharmacology, 12:587–599.

Zilles, K., Schleicher, A., Palomero-Gallagher, N., & Amunts, K. (2002b). Quantitative analysis of cyto- and receptor architecture of the human brain. In A. W. Toga, & J. C. Mazziotta, Brain mapping: The Methods (pp. 21:573–602). San Diego: Elsevier.

